# Mechanistic principles of hydrogen evolution in the membrane-bound hydrogenase

**DOI:** 10.1101/2024.03.16.585322

**Authors:** Abhishek Sirohiwal, Ana P. Gamiz-Hernandez, Ville R. I. Kaila

**Affiliations:** Department of Biochemistry and Biophysics, Stockholm University, 10691 Stockholm, Sweden

**Keywords:** Bioenergetics, archaeal metabolism, PCET, H2 production, multi-scale simulations, electric fields, enzyme catalysis

## Abstract

The membrane-bound hydrogenase (Mbh) from *Pyrococcus furiosus* is an archaeal member of the Complex I superfamily. It catalyzes the reduction of protons to H_2_ gas powered by a [NiFe] active site and transduces the free energy into proton pumping and Na^+^/H^+^-exchange across the membrane. Despite recent structural advances (1–4), the mechanistic principles of H_2_ catalysis and ion transport in Mbh remain elusive. Here we probe how the redox chemistry drives the proton reduction to H_2_ and how the catalysis couples to conformational dynamics in the membrane domain of Mbh. By combining large-scale quantum chemical density functional theory (DFT) and correlated *ab initio* wave function methods with atomistic molecular dynamics simulations, we show that the proton transfer reactions required for the catalysis are gated by electric field effects that direct the protons by water-mediated reactions from Glu21_L_ towards the [NiFe] site, or alternatively along the nearby His75_L_ pathway that also becomes energetically feasible in certain reaction steps. These local proton-coupled electron transfer (PCET) reactions induce conformational changes around the active site that provide a key coupling element via conserved loop structures to the ion transport activity. We find that H_2_ forms in a heterolytic proton reduction step, with spin crossovers tuning the energetics along key reaction steps. On a general level, our work showcases the role of electric fields in enzyme catalysis, and how these effects are employed by the [NiFe] active site of Mbh to drive the PCET reactions and ion transport.

**Significance statement:** Hydrogen (H_2_) serves as a crucial solar fuel in renewable energy systems that can be efficiently produced by microbial hydrogenases. Here we probe the elusive mechanistic principles underlying the H_2_ production in the ancient membrane-bound hydrogenase (Mbh) from the thermophilic archaeon *Pyrococcus furiosus*. Distinct from other hydrogenases, Mbh not only produces H_2_, but it couples this activity with ion transport across a membrane that powers the archaeal energy metabolism. Our study elucidates key mechanistic principles underlying H_2_ production and shed light on energy transducing enzymes that led to the evolution of modern mitochondrial respiratory enzymes.

## Introduction

The membrane-bound hydrogenase (Mbh) is an ancient enzyme found in the thermophilic archaeon *Pyrococcus furiosus* and one of the earliest members of the respiratory Complex I superfamily (1–7). Mbh powers the ferredoxin-driven (*E*_m_=*ca.* -480 mV at pH=7) reduction of protons to H_2_ (*E*_m_=-420 mV at pH=7), (1, 8–10) which is catalyzed by its [NiFe] active site. Mbh transduces the free energy (−120 mV=-2.7 kcal mol^-1^) into proton pumping (H^+^) and sodium (Na^+^) / proton (H^+^) exchange, generating a sodium motive force (smf) across the biological membrane that powers the Na^+^-dependent ATP synthesis of *P. furiosus* (11). Mbh is a 300 kDa transmembrane protein comprising 14 subunits, with the hydrophilic domain harboring the [NiFe] active site (MbhL) and three iron-sulfur (FeS) cluster (MbhN, MbhJ), responsible for the electron transfer, whereas the membrane domain catalyzes Na^+^ / H^+^ exchange (MbhA-G) and proton pumping (MbhM/MbhH) (**Figure 1)**. It is closely related to the respiratory Complex I (NADH:ubiquinone oxidoreductase) (5, 7), a large (0.5-1 MDa) redox-driven proton pump that drives electron transport and oxidative phosphorylation in aerobic respiratory chains. Mbh is thus a key enzyme for understanding not only the evolution of complex bioenergetic machineries, but also how H_2_ gas production powers the generation of an ion motive force across a biological membrane. Despite significant molecular understanding on soluble [NiFe] hydrogenases (12–15), its mechanistic principles remain puzzling and poorly understood.

**Figure 1.**
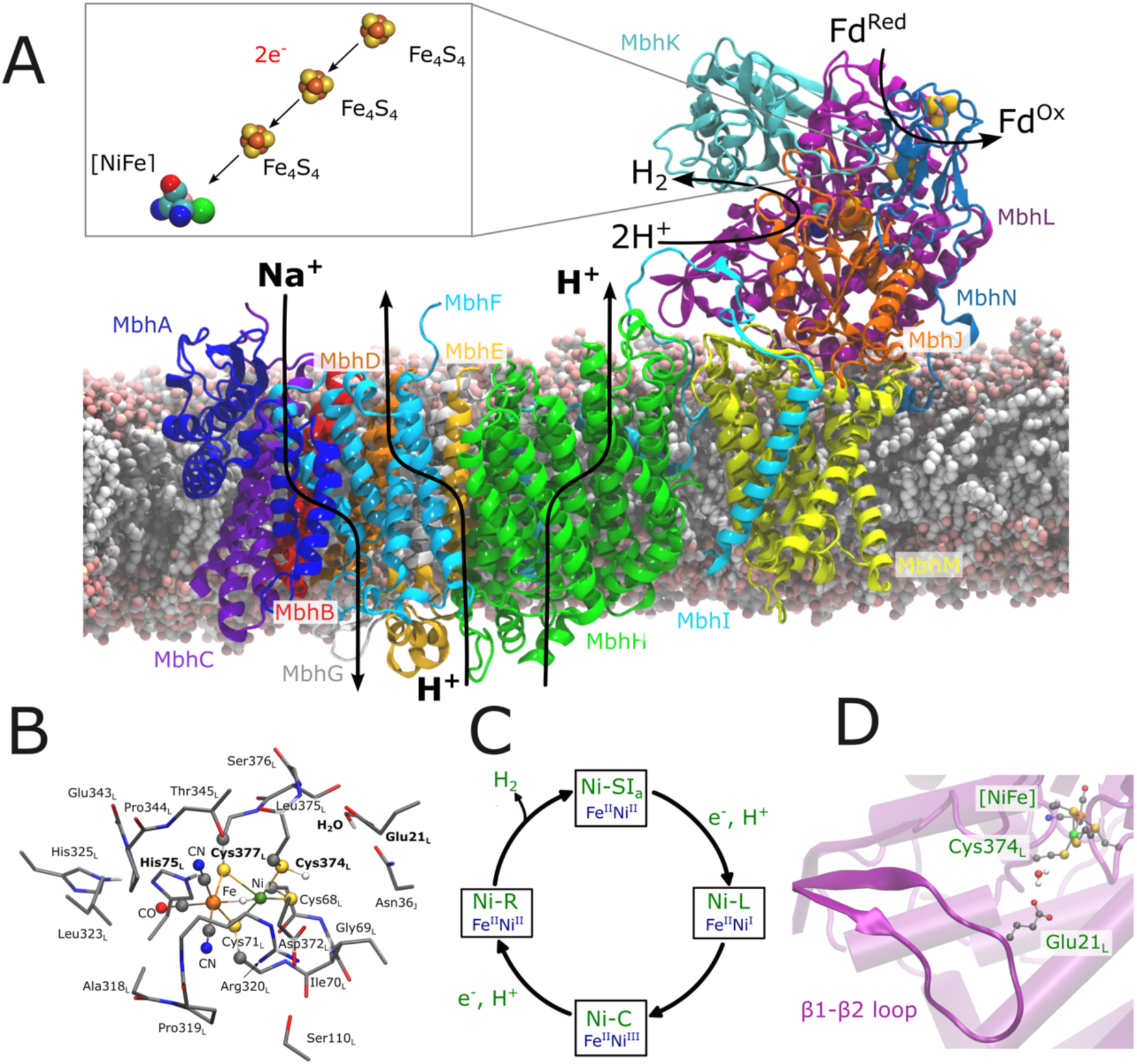
Structure, function, and catalytic cycle of Mbh. (A) Overall structure and function of Mbh, showing the hydrophilic domain, responsible for ferredoxin (Fd)-driven H^+^ reduction to H2 gas, and the membrane domain, responsible for proton pumping and Na^+^/H^+^ exchange across the membrane. Arrows along the transmembrane region show subunits responsible for the ion transport. Redox active cofactors responsible for the electron transfer from Fd to the [NiFe] center is also shown. (B) DFT models used in the present work to study the mechanism of [NiFe] catalysis, and (C) catalytic states associated with the H2 catalysis in [NiFe] hydrogenases. (D) Structure of the [NiFe] active site, the β1-β2 loop, harboring Glu21L, and catalytically active water molecules, hydrogen-bonding with Glu21L and Cys374L of subunit MbhL.

The electrons required for the proton reduction are delivered by ferredoxin (Fd) (16), a small (7.5 kDa) (16) soluble electron transfer protein that docks to the MbhN subunit (**Figure 1A**). Fd is a one-electron donor that stepwise transfers the electrons *via* the three iron-sulfur clusters [4Fe-4S] located in MbhN and MbhJ, to the [NiFe] center. The [NiFe] cluster (located in MbhL, **Figure 1A**) coordinates Cys71_L_, Cys377_L_, Cys68_L_, and Cys374_L_ (𝜎-acceptors), whereas the Fe center binds two CN^−^ (𝜎-donor) and one CO ligand (**Figure 1B**). Interestingly, the [NiFe] active site has evolved into the Q-reduction site of the modern respiratory Complex I, with a striking structural resemblance together with key modular adaptations (5, 7, 17–19) **(Figure 1D)**. Based on data from soluble hydrogenases, the Ni center undergoes redox changes required for the catalysis (Ni^I^, Ni^II^, Ni^III^ (12, 13, 20, 21), and possibly also a Ni^IV^ (22) form), whereas the iron remains in the Fe^II^ state during catalysis, with the diatomic strong field ligands ensuring its low spin (LS) configuration (13–15).

Although the catalytic principles of Mbh are still poorly understood, the related [NiFe] active site in the canonical hydrogenases undergoes proton-coupled electron transfer (PCET) reactions that stepwise reduce the Ni and provide protons for the H_2_ formation (or H_2_ splitting) reactions **(Figure 1B-C)** (13–15, 23–25).

The Ni^II^ has a *d*^8^ electronic configuration, which can result in either low spin (LS, *S*=0) or high spin (HS, *S*=1) configurations in hydrogenases (26–28). In the experimentally characterized Ni-SI_a_ (20) and Ni-R (29) states, the center remains in the Ni^II^ LS form, whereas the nickel is paramagnetic (*S*=1/2) in the Ni-L (30–32) and Ni-C (33) states, with Ni^I^ and Ni^III^, respectively. Recently, also a high valent LS Ni^IV^ (S=0) state was assigned based on Fourier-transform infrared spectroscopy (FTIR) and electron paramagnetic resonance (EPR) spectroscopic studies in a NAD^+^-reducing [NiFe] hydrogenase (22). In this regard, it was proposed that the hexa-coordinated Ni^IV^ can protect the active site from O_2_-induced oxidative damage. However, despite the involvement of multiple spin and oxidation states, the role of spin crossover in H_2_ catalysis is not well understood.

In addition to the unclear molecular principles underlying H_2_ formation, the pathways used for proton delivery to the active site in Mbh also remain debated. The protons required for the H_2_ evolution in the soluble [NiFe] hydrogenases are most likely transferred via a pathway involving Cys546 and Glu34 (*D. vulgaris Miyazaki F.,* hereafter *Dv*MF (34, 35); **Figure S1**) that connect via a hydrogen-bonding network to the bulk solvent (15, 24, 35, 36). Moreover, another pathway, via His82 (His75_L_ in Mbh, His88 in *Dv*MF, also known as “histidine pathway”) was described in O_2_-tolerant membrane-bound [NiFe] hydrogenase from *Ralstonia eutropha* (37, 38). In Mbh, four possible proton pathways lead from conserved Glu21_L_ (see below) to the [NiFe] center, via water networks at the interface between MbhL/M, MbhL/I/J, MbhL/N/J and MbhM/cleft (3). These putative pathways have also resemblance to proton pathways leading to the Q oxidoreduction site in Complex I (39, 40), although their function remain unclear.

In contrast to soluble hydrogenases, the H_2_ catalysis is functionally coupled to the ion pumping activity in the membrane domain of Mbh (1). While the overall metal coordination has remained conserved relative to the soluble [NiFe] hydrogenases, Mbh has certain key structural differences around the [NiFe] center that could mediate the coupling effects. One such modular adaptation is established around the conserved loop structure (linker region Ile10_L_-Lys24_L_) of the β1-β2 subunit, which harbors Glu21_L_, a residue that could shuttle protons to the active site (3, 41) (see **Figure 1D**). The same loop undergoes conformational changes in respiratory Complex I (42–44) and could thus also trigger the proton transport activity in the membrane domain also in Mbh. The distance between Glu21_L_ and the Ni-coordinating Cys374_L_ is rather large (4.8 Å) for a direct proton transfer in the cryo-EM structure of Mbh (PDB ID: 6CFW(1)), and the carboxylate sidechain is poorly resolved. In this regard, recent studies (3) suggested that Glu21_L_ could undergo a conformational change, similar to the proton shuttling Glu-242 in cytochrome *c* oxidase (45), and supports proton transfer to the active site. Interestingly, similar structural motifs around the active site are also present in the membrane-bound formate hydrogenlyase (FHL) (4), which also couples H_2_ formation with ion pumping. In FHL, the distance between Cys531_E_ and Glu193_E_ is around 7 Å in the cryo-EM structure (4), whereas the homologous Glu193_E_ of the soluble [NiFe] hydrogenases is located around 3.4 Å from Cys546 in the high-resolution crystal structure (35), and may also provide the protons for catalysis.

Here we probe the energetics of the catalytic cycle responsible for proton reduction and H_2_ formation in Mbh by combining large-scale quantum chemical models with density functional theory (DFT) and correlated *ab initio* wavefunction-based methods, and classical atomistic molecular dynamics (MD) simulations. Based on our multi-scale approach, we identify the role of conserved residues in the catalysis, and how electric field effects modulate reaction barriers. Our work provides a molecular basis for understanding the link between H_2_ catalysis and the ion transport activity across the membrane domain and highlighting key differences relative to the soluble [NiFe] hydrogenase, with implications on the evolution of the Complex I superfamily.

## Results

### Proton transfer energetics in [NiFe] catalysis

To obtain insight into the redox-triggered proton transfer energetics in Mbh, we first constructed large quantum chemical density functional theory (DFT) models of the [NiFe] active site. The models included in addition to the bimetallic core and its immediate ligands, also all first and second sphere protein ligands, and water molecules obtained from our atomistic molecular dynamics (MD) simulations, and leading to a molecular system with around 260 atoms (see **Figure 1B** and *Methods*). Based on the DFT models, we further optimized reaction pathways for proton transfer leading to H_2_ formation along the four experimentally characterized redox states (Ni-SI_a_, Ni-L, Ni-C, Ni-R) of the catalytic cycle (see **Figure 1C**). Our extensive benchmarking calculations relative to high resolution (0.89 Å) x-ray data (35) as well as *ab initio* wave-function theory (RPA, DLPNO-CCSD(T_1_)) show that the employed DFT methodology provides an accurate description of the geometry, electronic structure, and reaction energetics of the [NiFe] site (**Figures S1-S3**, **Tables S1-S4**).

We first explored the structure of the [NiFe] in the Ni-SI_a_ state, with the metal core modeled in the Fe^II^/Ni^II^ configuration, with the nickel in singlet (*S*=0, LS) or triplet (*S*=1, HS) state (**Figure 2A**). To obtain insight into the proton transfer energetics, we permutated the proton on the possible protonatable residues around the active site (Glu21_L_, Cys374_L_, Cys377_L_, Cys68_L_, His75_L_) and optimized reaction pathways for the proton transfer reaction between the different sites (**Figure S5-S7 and Table S5-S7**).

**Figure 2.**
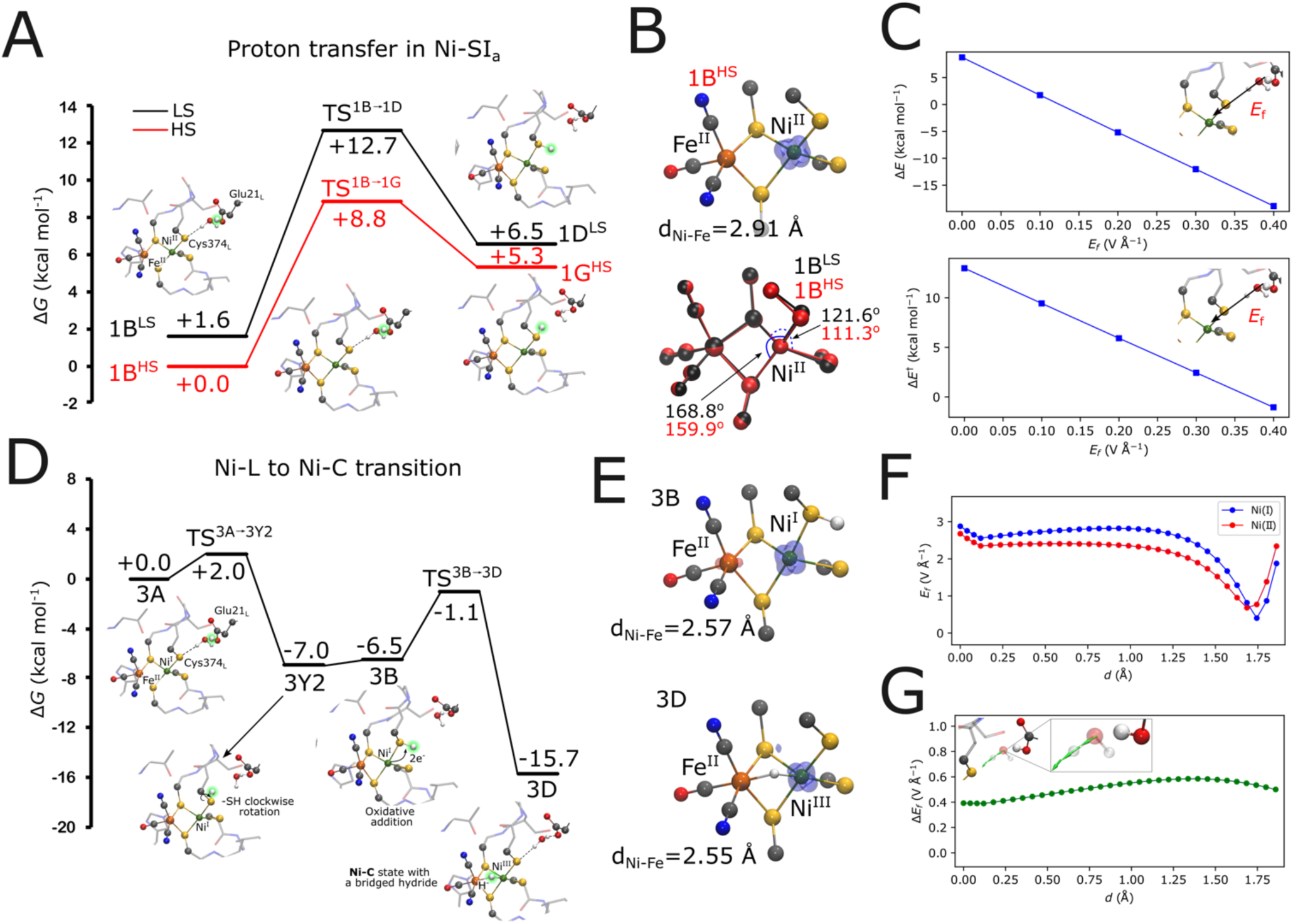
Reaction energetics in the Ni-SIa, Ni-L and Ni-C states. (A) Energetics of proton transfer from Glu21L to Cys374L in the low spin (LS) and high spin (HS) forms of the Ni-SIa state. The transferred proton is highlighted in green (B) *Top*: spin density distribution in the 1B^HS^ state, and *bottom*: overlay of optimized LS and HS models of the Ni-SIa state. (C) Dependence of the reaction energy and barrier on an electric field applied along the proton transfer coordinate. (D) Energy profile for proton transfer from Glu21L to Cys374L in the Ni-L state and subsequent transition to the Ni-C state. The transferred proton is highlighted in green. (E) Spin density distribution in the Ni-L (3B) and Ni-C (3D) states, with marked Ni - Fe distances. (F) Electric field strength along the H2O - Cys374L proton transfer coordinate in the Ni-SIa state (**1B^LS^**), and in the one-electron reduced Ni-L state. (G) Electric field difference in the Ni-SIa *minus* Ni-L states. *Inset*: the electric field vectors point towards the sulfur atom of the Cys374L upon reduction of Ni^II^.

Overall, the optimized geometries (**Figure 2B**, **Figure S5-S6**) suggest that the coordination environment around Ni^II^ in the low- and high-spin forms is almost identical with “see-saw” like coordination (**Figure 2B**); this could arise from the restricted movement of ligands due to structural constraints imposed by the protein scaffold. However, we observed subtle differences in the metal-ligand bond lengths between the spin states (*e.g.* Ni-S_C377_ is 2.19 Å and 2.38 Å in **1B^LS^** and **1B^HS^**, respectively. See Table S6 for a detailed analysis).

The water-mediated proton transfer from Glu21_L_ to Cys374_L_ has a modest free energy barrier (Δ*G*^‡^=+12.7 kcal mol^-1^ in LS, Δ*G*^‡^=+8.8 kcal mol^-1^ in HS), but protonation of Cys374_L_ is energetically unfavorable (Δ*G* = +6.5 kcal mol^-1^ in **1D^LS^**; +5.3 kcal mol^-1^ **1G^HS^**) (**Figure 2A**, see also **Figure S7**). When Glu21_L_ is protonated, both spin states are nearly isoenergetic (ΔΔ*G*=1.6 kcal mol^-1^), with a small overall preference for the HS form (**Figure 2A**). We find that the proton transfer from Glu21_L_ to Cys68_L_ (+15.6/+15.4 kcal mol^-1^ for **1E^LS^/IE^HS^)** or from His75_L_ to Cys377_L_ are also energetically highly unfavorable (**Figure S5-S6**). The energetics compares well with our wave-function based *ab initio* calculations, suggesting that our hybrid density functional treatment captures the reaction energetics of the system within an overall accuracy of a few kcal mol^-1^ for the electronic effects (**Table S3-S4**). Taken together, these calculations show that the overall proton transfer from Glu21_L_ or His75_L_ to the cysteine ligands or to the metal site in the Ni-SI_a_ state is unfavorable (see also **Table S6** for structural analysis).

One-electron reduction of Ni-SI_a_ leads to the Ni-L state, with a Ni^I^/Fe^II^ (*S*=1/2) configuration, with the reduction strongly favoring the proton transfer towards the [NiFe] cluster (**Figure 2D**, **Figure S8-S9, Movie S1**). In this regard, the proton transfer from Glu21_L_ **(3A)** to Cys374_L_ **(3Y2)** becomes strongly exergonic upon formation the Ni-L state (Δ*G*=-7.0 kcal mol^-1^) with a small reaction barrier (ΔG^‡^∼2.0 kcal mol^-1^, **Figure 2D**). Rotation of the protonated thiol of Cys374_L_ towards the [NiFe] site is nearly isoenergetic (see below, **3B**, -6.5 kcal mol^-1^; ∼92° clockwise with respect to -SH group in **3Y2**, see **Figure 2D**), with a two-center Ni-Fe metal-metal bond (**Figure 2E**, **Table S8-S9**) forming at the core (*cf*. also Ref. (31)). The proton transfer to Cys68_L_ (**3G1**, -4.1 kcal mol^-1^, **Figure S8**) is also exergonic, suggesting that the proton could populate different rotameric configurations of both Cys374_L_ and Cys68_L_, consistent with the experimentally observed multiple forms of the Ni-L state (46, 47). However, in contrast to the energetically favored Glu21_L_-mediated pathway, we find that the His75_L_-mediated proton transfer to the cluster is strongly endergonic (**Figure S8**) in the Ni-L state. Taken together, our findings suggest that the initial proton transfer reaction energetically favors the glutamate pathway (but see below).

Subsequent oxidation of the Ni center forms a Ni^III^/Fe^II^ (*S*=1/2) configuration, known as the Ni-C state. This reduces the transferred proton to a hydride (H^-^) ion, mediated by a *metal*-to-*ligand* electron transfer between the Ni and the H^+^ (**Figure 2D**, **Figure S8-S9, Table S8-S9**). Protonation of the metal-core involves further rotation of the Cys-H bond, allowing the H^-^ to coordinate to the bridging (𝜇-H^-^) position (**3B** →**3D** Δ*G*^‡^=+5.4 kcal mol^-1^) with both the Ni^III^ and Fe^II^ (*d*(Ni-H^-^) =1.60 Å / *d*(Fe-H^-^) =1.70 Å, **Figure S9**, **Movie S2**). The established state closely resembles the experimentally characterized Ni-C form of the soluble [NiFe] hydrogenases (48). In contrast, we find that the formation of a terminal Ni-H^-^ configuration (*d*(Ni-H^-^) =1.44 Å / *d*(Fe-H^-^) =3.54 Å) is energetically unfavored **(Figure S8**, see also **Table S9** for geometric analysis). The free energy profiles thus suggest that the conversion of Ni-L to Ni-C state is exergonic (**3Y2**→**3D**, Δ*G*=-8.7 and Δ*G*^‡^ = +5.8 kcal mol^-1^), and support the progression of the reaction towards the reduced metal-bridged (Ni-H^-^-Fe) hydride state (**Figure 2D-2E**), but the additional protonation of the cysteine ligands is energetically unfavored in this state (**Figure S10-S11**), preventing further proton uptake in the Ni-C state (see below).

### Electric field induced proton transfer reactions

To probe the molecular basis for the redox-triggered change in reaction barriers and thermodynamic driving force for proton transfer in the Ni-SI_a_ → Ni-L transition, we quantified electric field (***E***) effects around the active site. Our calculations suggest that the formation of the reduced Ni-L state increases the electric field along the proton pathway leading from Glu21_L_ to Cys374_L_ by |Δ***E***_f_| = 0.6 V Å^-1^ relative to the oxidized Ni-SI_a_ state, with the reduction resulting in an electric field vector that point along the proton pathway towards Cys374_L_ (**Figure 2F-2G**). To further test how the electric fields could modulate the reaction barriers, we applied an external field in the direction of the H_2_O → Cys374_L_ bond in the Ni-SI_a_ state. Interestingly, this applied field similarly lowers the reaction barriers and reaction energy, with a linear shift with increasing field strengths that closely resembles the effects observed upon formation of the Ni-L state. These findings suggest that redox-triggered electric field effects could drive the proton transfer towards the [NiFe] center (**Figure 2C**) by similar effects as observed for other energy-transducing systems such as cytochrome *c* oxidase (49), Complex I (43), and Photosystem II (51) (*cf*. also (50)).

### Mechanism of H_2_ formation in the Ni-R state

Our DFT calculations suggest that the further one-electron reduction of Ni-C leads to a Ni^II^/Fe^II^ configuration, known as the Ni-R state, with energetically accessible LS and HS forms. To probe the thermodynamic and kinetic feasibility of transferring the second proton required for the H_2_ formation in this state, we studied the energetics of proton transfer from both Glu21_L_ and His75_L_ towards the hydride bridging the [NiFe] core, and based on these, we explored the energetics of forming H_2_ within the [NiFe] core.

We find that the proton transfer from Glu21_L_ (**4A^LS/HS^**) to Cys374_L_ (**5E2^LS/HS^**) becomes strongly downhill in the Ni-R state (Δ*G* =-14.7 /-12.0 kcal mol^-1^ in LS / HS, **Figure 3**), whereas the proton transfer to Cys68_L_ is endergonic (by *ca.* Δ*G* = +5.2/+3.2 kcal mol^-1^, see **Figure S12-S16, Table S10-S12**). The protonation of Cys374_L_ has a small reaction barrier (**4A^LS/HS^**→**5E2^LS/HS^**, Δ*G*^‡^ = +7.7 /+5.6 kcal mol^-1^ for LS / HS), which supports that the reaction is kinetically feasible. Moreover, due to the small energy gap between the spin states, the proton transfer step could employ spin crossover between the LS and HS forms, which in turn would reduce the reaction barriers along the HS pathway (**Figure 3**). Similarly, as for the first proton transfer step (see above), we find that the forward progression of the reaction involves rotation of the thiol group of Cys374_L_ (**5E2^LS/HS^**→**4B^LS/HS^**, Δ*G* = +6.1 / +6.0 kcal mol^-1^ for LS / HS) towards the metal-bound hydride. From here, the H_2_ formation takes place by the oxidative addition at the Ni^II^ center, thereby forming a di-hydride bound Ni^IV^/Fe^II^ species in the LS state (**4B1^LS^**, **Movie S3**, **4B^LS^**→**4B1^LS^**Δ*G*^‡^*=* +6.3 kcal mol^-1^), with an octahedral coordination around a putative Ni^IV^ state. Subsequent reduction of the Ni^IV^ to Ni^II^ (**4B1^LS^**→**4D^LS^,** Δ*G*= +0.3, Δ*G*^‡^*=* +2.8 kcal mol^-1^) couples to the formation of the H_2_ in the Ni^II^ form **(4D^LS^**, Ni-H_2_ is 1.59 Å). In contrast, the H_2_ formation along the HS surface is highly endergonic (**4B^HS^**→**4E^HS^,** Δ*G*=+12.2, Δ*G*^‡^ = +17.9 kcal mol^-1^, **Figure 3**) and requires a large rotational movement of the Cys374_L_ thiol group towards the bound hydride, with H_2_ **(4E^HS^)** forming at the bridging position (*d*(Ni-H_2_) = 2.08 Å, *d*(Fe-H_2_) =1.68 Å).

**Figure 3.**
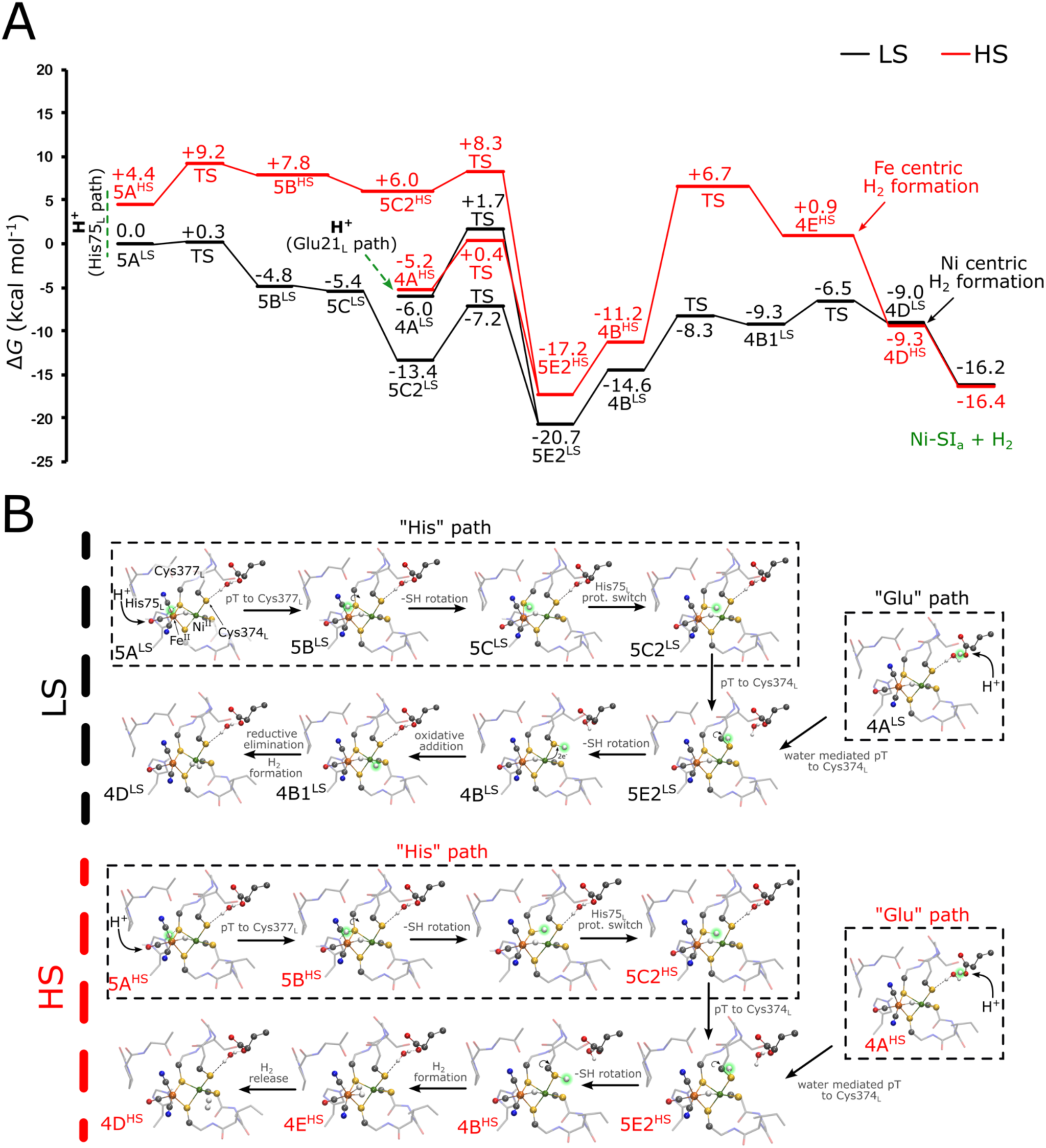
Free energy profile of H2 formation in the Ni-R state. (A) Free energy profile of the H2 formation in the low spin (LS, in black) and high spin (HS, in red) configurations of the Ni-R state. Energetics of proton transfer via the histidine (His75L) and glutamate (Glu21L) pathways. (B) Structure of different intermediates involved in the H2 formation step. The structures of the transition states are shown in **Figures S14-S15**.

Notably, the high-resolution structure of the [NiFe] hydrogenase from *Dv*MF in the Ni-R state (35) also shows a bridging hydride and a protonated Cys374_L_ (referred to as Cys546 in *Dv*MF), with a shorter Ni-H^-^ bond (1.58 Å) relative to the Fe-H^-^ bond (1.78 Å). This state closely resembles our energetically favored LS form of the Ni-R state (**5E2^LS^**, Ni-H^-^ / Fe-H^-^ are 1.58 Å / 1.70 Å), suggesting that the H_2_ formation is strongly favored along the LS pathway.

### Involvement of the histidine pathway in the H_2_ formation

To probe the possible role of His75_L_ in both the H_2_ formation and proton delivery (37, 38), we also optimized all putative states along the catalytic cycle, but with His75_L_ modeled in its protonated (HisH^+^) form (**Figure S17-S25, Table S14-S21**). For the Ni-R state, we find that the proton transfer from His75_L_ **(5A^LS^)** to Cys377_L_ **(5B^LS^)** is exergonic and has a small reaction barrier (**5A^LS^**→**5B^LS^**, Δ*G*=-4.8, Δ*G*^‡^=+0.3 kcal mol^-1^, **Figure 3**). Tilting of the Cys377_L_ **(5C^LS^)** allows the -SH group to move away from the bridging position (**5B^LS^**→**5C^LS^**, Δ*G* = -0.6 kcal mol^-1^) that leads to a subtle shift in the sidechain of His75_L_ (**Figure 3B**). The proton transfer from Cys377_L_ **(5C2^LS^)** to Cys374_L_ **(5E2^LS^)** is exergonic (Δ*G* = -7.3 kcal mol^-1^) with a small reaction barrier (Δ*G*^‡^ = +6.2 kcal mol^-1^) (**Figure S12-S14**), whilst the subsequent proton transfer from Cys374_L_ towards the hydride is also energetically feasible along the same mechanism as discussed above **(5E2^LS^**→**4B^LS^**→**4B1^LS^**→**4D^LS^)**. Interestingly, the proton transfer along this pathway is exergonic in the LS configuration, but endergonic in the HS state (**5A^HS^**→**5B^HS^**, Δ*G*=+3.4 kcal mol^-1^ and Δ*G*^‡^ = +4.8 kcal mol^-1^, **Figure 3**, **Figure S14-S16**).

In addition to its possible role as a proton uptake pathway, we find that the protonation of His75_L_ modulates the proton transfer energetics along the Glu pathway. In this regard, the proximity of the positively charged HisH^+^ next to the [NiFe] center leads to an increase in proton transfer barriers for Glu21_L_ to Cys374_L_ in the Ni-SI_a_ to Ni-L transition (**Figure S17-S21**), as well as an increase in the reaction barrier for hydride binding during the Ni-L to Ni-C transition (see also **Figure S21)**. In contrast, the proton transfer barrier from Glu21_L_ to Cys374_L_ decreases by +5.4 kcal mol^-1^ in the LS Ni-R state (**Figure S23-S25**). The latter effect could arise from a directional electric field (Δ*E*_f_ = 0.1 V Å^-1^) formed along the Glu21_L_-Cys374_L_ pathway upon protonation of His75_L_ (**Figure S26**). We find that the His75_L_ protonation favors the HS over the LS form in the Ni-SI_a_ state (**Figure S19**), but does not affect the spin energetics in the other redox states (**Figure S25**). Taken together, these findings suggest that protonation of His75_L_ could act as a pH-switch that modulates the reaction energetics, consistent with altered [NiFe] activity dependence on the pH and its possible functional “pH sensing” (32).

Taken together, our findings demonstrate that the second proton transfer required for H_2_ catalysis could both energetically and kinetically take place via the “Glu pathway” (in both LS and HS configurations) or the “His pathway” (in the LS configuration). The latter requires protonation of His75_L_, suggesting that the His pathway could be active at low pH upon formation of the Ni-R state, and regulate the activity of the [NiFe] center.

### H_2_ release and active-site recovery to Ni-SI_a_

Our DFT calculations suggest that the binding of H_2_ in the Ni-R state is modulated by the spin state. In the LS form (**4D^LS^**), H_2_ prefers binding in a side-on, 𝜂*^2^*configuration to Ni^II^, whereas H_2_ binds in a bridging position between the Ni and the Fe in the HS form (**4E^HS^**), although the state is energetically highly unfavored (Δ*G*_HS-LS_= +10 kcal mol^-1^, **Figure 3A**). However, dissociation of H_2_ leads to crossing of the spin states, and results in a degenerate *apo*-Fe^II^Ni^II^ state for both the HS and LS forms (**4D^HS^** and **4D^LS^**, Δ*G*= +0.3 kcal mol^-1^). These findings suggest that the H_2_ formation and dissociation could involve spin crossover. We find the H_2_ dissociation is coupled with a rather large entropic (*T*Δ*S*) effect of - 9.3 kcal mol^-1^ that could be relevant for transducing the free energy for ion transport (**Figure S13**). Our calculations further show that the release of H_2_ restores the Ni-SI_a_ state.

### Redox-triggered conformational changes triggers the ion transport machinery

In order to probe the conformational dynamics coupled to catalysis, we performed atomistic molecular dynamics (MD) simulations of Mbh embedded in a membrane-water-ion environment to probe how the redox-coupled proton transfer reaction induce conformational and hydration changes (**Table S22-S23**). In this regard, we derived atomistic force field parameters based on the quantum chemical models for [NiFe] cluster along the key steps of catalytic cycle that allowed us to study the coupling between the redox catalysis and the conformational dynamics (3).

Our MD simulations suggest that reduction of the [NiFe] center changes the conformational dynamics of Glu21_L_, and leads to an increase in the “flipped-in” conformation of the residue. This increases the occupancy of the proton transfer-mediating water molecule, bridging the Glu21_L_-Cys374_L_ gap during Ni-SI_a_ → Ni-L and Ni-C → Ni-R transitions (**Figure 4**) (3). Interestingly, the proton transfer from Glu21_L_ to Cys374_L_ induces a conformational change, which leads to dissociation of the water molecule, and favors the outward flipping of the anionic Glu21_L_ (**Figure 4A-C)**, which could prevent back-transfer of the proton from Cys374_L_ to Glu21_L_/bulk solvent, but also provide a possible coupling element that triggers ion transport in the membrane domain of Mbh (see below). Recent cryo-EM structures of the FHL complex (4) also show a cavity between Cys374_L_ and Glu21_L_ (with a distance of around 7 Å) that could occupy a similar proton transfer-mediating water molecule, while this feature has not been reported for soluble [NiFe] hydrogenases (*Dv*MF). However, we note that although the Cys546 and Glu34 form a strong hydrogen-bonding interaction in *Dv*MF, FTIR experiments (24) suggest that a “dangling” water molecule re-aligns upon changes in protonation states during the Ni-C → Ni-L transition. The outward flip of the protonated Glu21_L_ in our simulations of the Ni-C state furthers stresses the possible functional role of the His75_L_ in providing a second proton for the H_2_ formation (**Figure 3B**). Sequence and structure comparison with the Complex I superfamily (**Figure S30**) show that Glu21_L_ occupies the same position as His38^NDUF2^ (*T. thermophilus* numbering), which is likely to function as a proton donor in the quinone reduction process in Complex I (39) (*cf*. also (52–54)).

**Figure 4.**
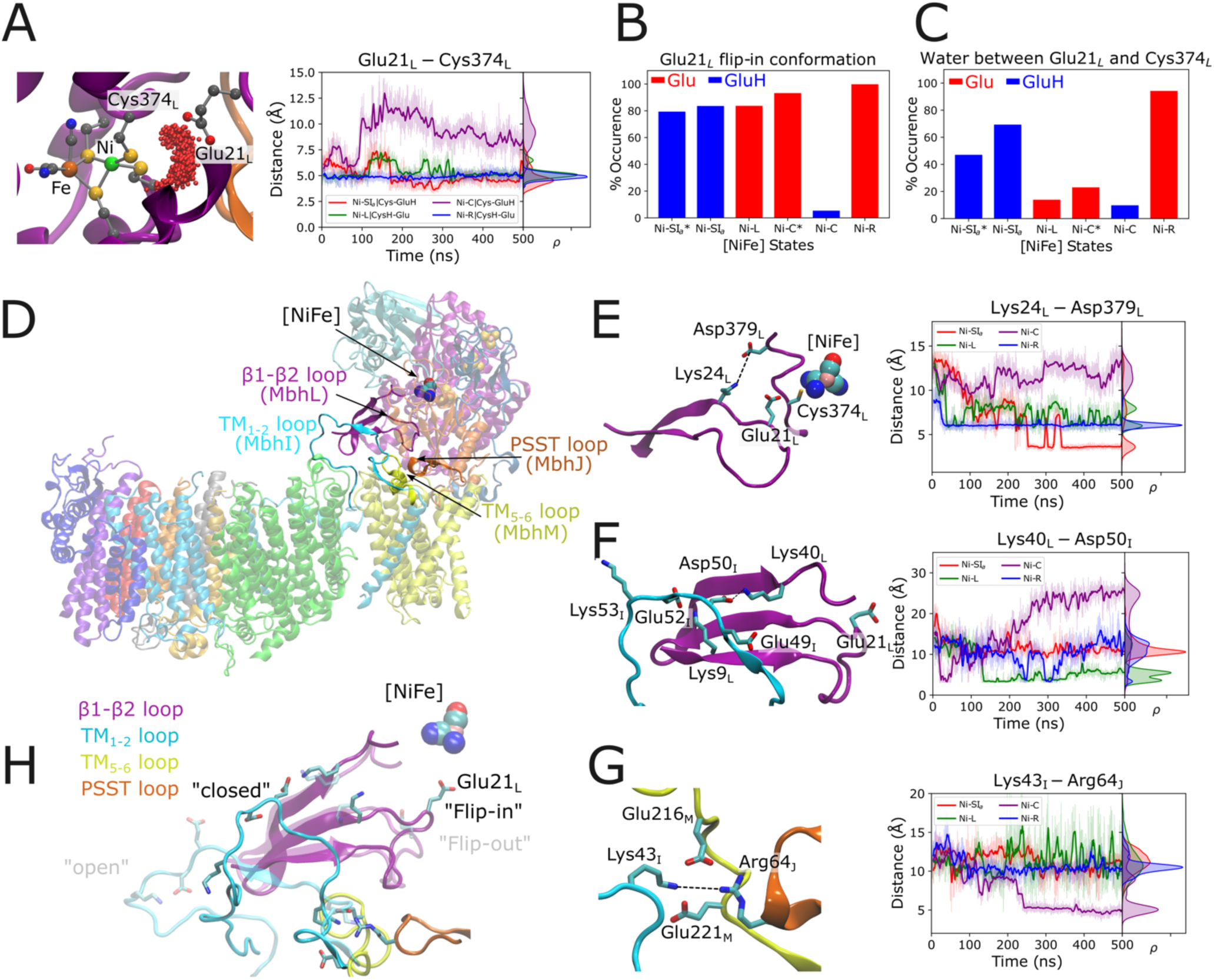
Redox-triggered conformational coupling between redox catalysis and loop dynamics. (A) Clustering of water molecule between the Glu21L and the [NiFe] cluster, and the distance between Cys374L and Glu21L in different catalytic states from MD simulations. (B) Population of “flipped-in” conformation of Glu21L in different catalytic states. (C) Water occupancy between Glu21L and Cys374L in MD simulations. (D) Structure and location of the loop network and the [NiFe] cluster of Mbh. (E) Effect of the Glu21L “flip” on the ion-pair dynamics in different catalytic states. The outward motion of Glu21L in the Ni-C state favors dissociation of the ion-pair with Lys24L and Asp379L, and contraction of the β1-β2 loop (see panel H). (F) Interaction of the TM1-2 loop (MbhI) and the β1-β2 loop (MbhL). The contraction of β1-β2 leads to dissociation of an ion-paired network between different subunit, and an outward motion of the TM1-2 loop (see also panel H). (G) Interactions between the TM1-2 loop, the TM5-6 loop, and the PSST loop. The TM1-2 loop forms an ion-pair between Lys43I, Glu216M, Glu221M, and Arg64J. (H) Conformational changes in loops upon conformational changes in Glu21L (“inward” - opaque and “outward”-transparent orientations). Subsequent conformational changes in the TM1-2 loop are also shown (“open” vs “close”). Details of the redox and protonation state in various state are shown in **Tables S22-S23**.

We next analyzed how the redox and protonation changes in the active site could activate the ion transport activity across the membrane, by probing conformational changes in the membrane domain and surrounding loop structures. In this regard, it was recently suggested (43) that the quinone reduction in Complex I couples to conformational changes in the surrounding loop regions that could trigger a π-to-α transition in the transmembrane helix (TM3^ND6^) enabling proton transport in the membrane domain. Interestingly, many of these loop regions are also conserved in Mbh (**Figure 4D-H**, **Figure S28**). Our MD simulations suggest that loop regions of β1-β2 (MbhL) and TM1-2 (MbhI) undergo conformational changes during the Ni-C → Ni-R transition (**Figure 4**, **Figure S27-S28**). In the Ni-C state, we observe an outward flip of the protonated Glu21_L_ and a contraction of the β1-β2 loop, which results in the dissociation of the Lys24_L_-Asp379_L_ ion-pair. This conformational change destabilizes an ion-paired network, involving Lys40_L_, Lys9_L_, Glu49_I_, and Asp50_I_, similarly as in Complex I (43). Interestingly, when the MD simulations are performed in other catalytic state (Ni-SI_a_, Ni-L, Ni-R), we observe stable interactions between the loop region of TM1-2 and β1-β2. Destabilization of the ion-paired network between the TM1-2 and the β1-β2 loop in the Ni-C state leads to large-scale conformation change of the TM1-2 loop that propagate towards the membrane-arm (**Figure 4H**), consistent with the high B-factors and blurred density of the TM1-2 loop observed in the cryo-EM data (1). Despite common redox-driven electrostatic and conformational changes that resemble the coupling elements suggested for Complex I, we find that many of the charged residues in the TM1-2 (Glu49_I_, Glu52_I_, Lys53_I_) are unique to Mbh. Moreover, the region is more hydrophobic in Complex I relative to MbhM – a feature that could have evolved to support binding of the non-polar quinone substrate.

## Discussion

In this work we have proposed a molecular mechanism of the ancient membrane-bound hydrogenase, which reduces protons to form H_2_ gas and couples this redox-chemistry to ion pumping across the archaeal membrane. To this end, we derived an electronic structure-level understanding of the H_2_ formation at the [NiFe] active site, and probed how the redox-driven large-scale conformational changes in loops that connects the electron transfer domain with to the membrane module.

Despite some catalytic similarities to soluble hydrogenases (14, 15, 23), Mbh also shows notable differences in its proton transfer mechanisms (**Figure 5A**). Two distinct proton transfer pathways are identified in Mbh that are operational based on the [NiFe] redox state. Our results suggest that the first proton transfer (in the Ni-SI_a_ → Ni-L transition) takes place via Glu21_L_ (Glu pathway), while the second proton (Ni-C →Ni-R transition) can occur either via Glu and the His (His75_L_) pathways. Remarkably, the reaction barrier for the proton transfer between Glu21_L_ and Cys374_L_ during the Ni-C → Ni-R transition is higher (7.7 kcal mol^-1^) than in the Ni-SI_a_ → Ni-L transition (2 kcal mol^-1^) (**Figure 2D** and **Figure 3A**). However, the barrier decreases (by *ca*. 5 kcal mol^-1^) in the presence of a protonated His75_L_ due to electric field effects (**Figure S25**). These findings suggest a possible interplay between the two proton transfer pathways in facilitating the delivery of the second proton to the active site.

**Figure 5.**
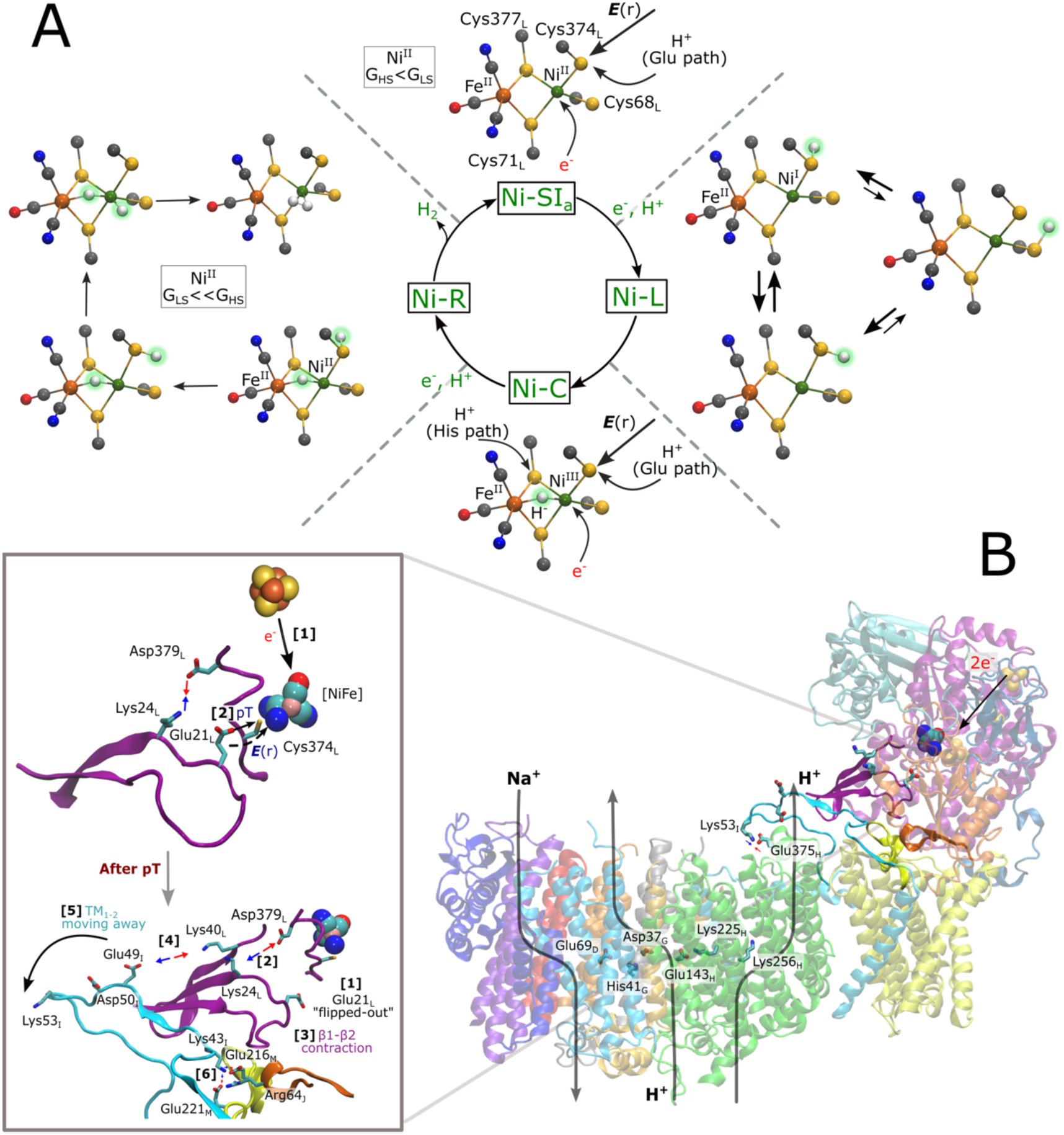
Structural models of intermediate states in the catalytic cycle and proposed coupling elements. (A) Structural models for the Ni-SIa, Ni-L, Ni-C, and Ni-R states in Mbh. The catalysis proceeds via one-electron reduction of Ni-SIa, and subsequent generation of directed electric field gradient that drives the proton transfer from Glu21L to Cys374L/Cys68L, forming the Ni-L state. The high spin (HS) state is more stable relative to the low spin (LS) form for the Ni-SIa state (*G*HS < *G*LS). Different structures for Ni-L, with energetically degenerate rotamers of the protonated Cys374L or protonated Cys68L. Subsequent oxidation of the Ni^I^ leads to the Ni-C state (Ni^III^) with a bridging hydride (μ-H^-^). Subsequent one-electron reduction of the Ni-C state forms the Ni-R state (Ni^II^), followed by proton transfer along the Glu or His pathways that result in the protonation of Cys374L or Cys377L, respectively. Possible representative structures of the Ni-R state are shown along the H2 formation reaction (see also Figure 3). The H2 formation in the Ni-R state is favored by the LS Ni^II^ form (*G*LS << *G*HS). (B) Suggested redox-driven long-range coupling elements in Mbh. *Top inset:* Primary signal transduction steps induced by the [NiFe] redox chemistry involve: [1] one-electron reduction of the [NiFe] by the proximal FeS cluster, which induces a directed electric field toward Cys374L that [2] leads to proton transfer from Glu21L to Cys374L. The “flipped-in” orientation of the Glu21L favors the proton transfer and formation of an ion-paired interaction between Lys24L and Asp379L. *Bottom inset:* Following the proton transfer step: [1] Glu21L flips in its “outward” conformation, which [2] favors dissociation of the Lys24L-Asp379L ion-pair. Subsequently, the β1-β2 loop undergoes a contraction [3], which in turn favors the dissociation of ion-pairs between the TM1-2 and the β1-β2 loop [4]. This cascade leads to conformational changes in the TM1-2 loop, which moves towards the membrane domain. The motion is supported by contacts between Lys53I and Glu375H. [6] These conformational changes also lead to the formation of ion-pairs between TM1-2 (MbhI), PSST (MbhJ), and the TM5-6 (MbhM) loops. The interaction of the TM1-2 loop with the MbhH subunit via Lys53I-Glu375H is shown, together with key residues along the putative proton pathway in MbhH.

It is to be noted that the coordination environment around the Ni and Fe in different catalytic states could also play a critical role in the activation of the two proton pathways. Specifically, in the Ni-L state, we observed a strong coordination between Ni and S (Cys377_L_), with a bond length of 2.30 Å, whilst in the Ni-R state, this bond length increased to 2.74 Å, whilst the Ni-S (Cys374_L_) bond distance remained nearly identical in both states. This suggests a weaker binding of Cys377_L_ to the Ni in the Ni-R state as compared to the Ni-L state, indicating that it could tune the p*K*_a_ values of residues and possibly altering the ligand field environment to facilitate proton transfer along His pathway during the Ni-C → Ni-R transition. The activity of the Glu pathway during Ni-C →Ni-R is likely to also depend on the conformation of the β1-β2 loop, which harbors Glu21_L_, and could function as coupling element during the redox-driven proton pumping.

In soluble hydrogenases, both protons could be transported via the Glu pathway (15). In this regard, Evan *et al.* (55) studied the role of Glu28 (Glu34 and Glu21_L_ in *Dv*MF and Mbh, respectively) in the O_2_-tolerant [NiFe] Hyd1 from *Escherichia coli* during H_2_ oxidation by using an E28Q mutant, which accumulated mainly the Ni-R and Ni-C states. These findings led to the conclusion that Glu28 is not essential for the proton transfer during the transition from Ni-R to Ni-C, whilst the residue was found to be critical in the subsequent Ni-C → Ni-SI_a_ transition. These results are consistent with our work, in which we suggest that the first proton originates exclusively from the “Glu” pathway, whilst the second proton could be transported along either pathway. The two proton transfer pathways in Mbh thus shows a unique resemblance to the proton transfer linked to Q reduction in Complex I, where both His38^NDUF2^ (part of the β1-β2 loop) and Tyr87^NDUF2^ function as the likely proton donors (39).

We note that several structural models for the Ni-L (46, 47) and Ni-R (23, 56, 57) states have been reported based on spectroscopic signatures, and indicating different structural isomers. For instance, temperature-dependent FTIR studies of the soluble [NiFe] hydrogenases observed protonation/deprotonation of the residue homologous to Cys374_L_ (Cys546 in *Dv*MF) with an Δ*H* and Δ*S* of 1.5 ± 0.8 kcal mol^-1^ and 6.1 ± 10.3 kcal mol^-1^ K^-1^, suggesting that the cysteine residue can indeed undergo protonation change in the Ni-L state (46). This compares well with our finding on the exergonic proton transfer from Glu21_L_ to Cys68_L_ or Cys374_L_ in the Ni-L state. Similarly, a recent IR and EPR investigation on a regulatory [NiFe]-hydrogenase from *Cupriavidus necator,* proposed that the interconversion between the Ni-L_1_ and Ni-L_2_ forms does not require the breaking of covalent bonds (47), which is consistent with our finding of two isoenergetic rotamers (**3Y2** and **3B**) of the protonated Cys374_L_ in the Ni-L state (**Figure 5A**). Similar to the Ni-L state, at least three unique spectroscopic signals have been identified for the Ni-R state, implying the co-existence of multiple metastable species. Based on our findings, these states could arise from the different rotamers of the protonated Cys374_L_ (**5E2^LS^** and **4B^LS^**), from the different H_2_ binding modes, and/or the dihydride bound state (**4B1^LS^** and **4D^LS^**) (**Figure 5A**)

As discussed above, our study reveals striking similarities between Mbh and the respiratory Complex I, such as conformational changes in the β1-β2 loop, the TM1-2 (MbhI), TM5-6 (MbhM), and the PSST loop (MbhJ) that are conserved within the superfamily, whilst the MbhH subunit is homologous to the antiporter-like Nqo12 subunit of Complex I. We suggest that the β1-β2 loop plays a critical role in the H_2_ catalysis by shuttling protons from the bulk towards the [NiFe] active site, whilst enabling a redox signal propagation towards the membrane domain that triggers ion pumping across the membrane (3).

The long-range signal transduction between the electron transfer activity in the hydrophilic domain and the ion transport in the membrane domain could be achieved through the redox state of the [NiFe] cluster and protonation state of Cys374_L_ and Glu21_L_. For each catalytic turnover, the conformational changes could be triggered during the Ni-C → Ni-R transition via motion in the network of loops (**Figure 5B**). The primary coupling event could involve an outward flip of Glu21_L_ and the subsequent contraction of the β1-β2 loop, which further leads to changes in the ion-pairs in TM5-6 (MbhM), PSST (MbhJ), and the TM1-2 loop (**Figure 4H**). We note the TM1-2 loop shows a lower sequence conservation (*ca.* 20%) relative the same region in Complex I (**Figure S31**). In this regard, we note that Asp379_L_ is conserved in Mbh and Complex I, but not present in hydrogenases, and suggesting that the Lys24/Asp379 interaction of MbhL could be important for the energy transduction (**Figure 4E**). In contrast, Glu49_I_ is unique for Mbh (**Figure 4F**), and forms an ion-pair with Lys40_L_, whilst MbhM (NuoH in Complex I) comprises the unique Glu216_M_ that interacts with Arg64_J_ (**Figure 4G**). The identified electrostatic network provides an important basis for future mutagenesis studies.

## Conclusions

The membrane-bound hydrogenase (Mbh) from *Pyrococcus furiosus*, a member of the Complex I super-family, serves as an intriguing system for studying the interplay between [NiFe]-enabled H_2_ catalysis and ion transport and its evolutionary relation to Complex I, which catalyzes quinone reduction. By combining density functional theory (DFT) with correlated *ab initio* methods, and atomistic molecular dynamics simulations, we derived key insight into the molecular mechanism of H_2_ evolution in Mbh, and the energetics and dynamics linked to this process. We suggested that Mbh employs similar redox-driven conformational changes as Complex I, particularly around the conserved β1-β2 loop that could be mechanistically important for transducing the redox energy into ion transport across the membrane. We suggest that Mbh employs electric field effects formed by redox and protonation changes – a mechanism that could serve as a general energy transduction principle in nature.

## Materials and Methods

### Quantum Chemical Models

Quantum chemical DFT models were built to probe the geometric and electronic structure of [NiFe] active site of Mbh. The DFT models were built by combining conserved parts of a high-resolution x-ray structure (at 0.89 Å resolution, PDB ID: 4U9H) of the soluble [NiFe] hydrogenase from *Dv*MF, resolved in the Ni-R state (35), together with our MD-relaxed system of Mbh (3), where we substituted all residues that are unique for Mbh (**Figure 1B**). The central-core of the model contained the Fe, Ni, the hydride, two CN^-^ and one CO ligand, and all residues in the first and second coordination sphere (Asn36J, Glu21L, Cys68L-Gly69L-Ile70L-Cys71L, His75L, Ser110L, Ala318L-Pro319L-Arg320L, Leu323L, His325L, Glu343L-Pro344L-Thr345L, Asp372L, Cys374L-Leu375L-Ser376L-Cys377L and a water molecule) (**Figure 1B**). The residues with a truncated backbone were terminated by a methyl group (-CH3) at the Cɑ atom. The Cɑ atoms and the linker hydrogen atoms of residues were fixed at their x-ray position during the geometry optimization. The model contains a total of 264 atoms. All geometry optimizations (ground state, transition state, product states) were performed at the TPSSh level with the central core of the active site comprising of Fe, Ni, S, two CN^-^ and CO assigned def2-TZVP basis-set, while where the rest of the atoms were described with the def2-SVP basis sets. We also applied the multipole accelerated resolution of identity (MARI-J) approximation during the optimization protocol, whereas dispersion effects were included using the empirical dispersion correction with Becke-Johnson damping (D3-BJ). A higher DFT integration grids (m4) and tighter SCF thresholds were used throughout (scfconv 8) in all computations. We also employed the implicit solvation COSMO scheme with dielectric constant set to 4.0. The molecular Hessian used for estimation of free energies were computed numerically at the TPSSh/def2-TZVP/def2-SVP level of theory (same as optimization protocol), with scaling factors (0.9615) of the vibrational frequencies based on,

The reaction pathways were optimized between reactant and product states using a method related to zero-temperature string method, as implemented in TURBOMOLE 7.5.1(58). The final single-point energy calculations were performed with B3LYP* (with 15% Hartree-Fock exchange) and scalar relativistic effects were included using the exact two-component (X2C) Hamiltonian. We used x2c-TZVPP for Fe and Ni, and x2c-TZVP basis-set for rest of the atoms. The choice of B3LYP* functional was chosen based on its performance against the random phase approximation (RPA) method and domain-based pair natural orbital coupled cluster with singles, doubles, and full iterative triples (DLPNO-CCSD(T1)) calculations to predict spin-state energetics accurately. The final single-point energy and molecular Hessian calculations were performed on a DFT model (Figure S4). Results of the benchmark (geometry, spin-state and reaction energetics) are shown in the Supporting Information. All the calculations were performed with the TURBOMOLE v. 7.5.1 (58).

### Molecular dynamics simulations

Atomistic molecular dynamics simulations of the Mbh (PDB ID: 6CFW) (1) model were performed using the CHARMM36 force field for protein/lipids and water. Parameters for the FeS clusters and [NiFe] in different catalytic states (see Table S22-S23) were derived from *in-house* DFT calculations (https://github.com/KailaLab/ff_parameters), with the remaining system treated using the CHARMM36m force field (59). The system was embedded in a 1-palmitoyl-2-palmitoleoyl-sn-glycero-3-phosphoinositol (PYPI) membrane using CHARMM-GUI (60) modelling unresolved sidechains (3), as well as different protonation states with lipids in the cleft between MbhM/MbhH. The model was embedded in a 200 x 100 x 168 Å^3^ box comprising TIP3P water molecules and ions to mimic a 250 mM NaCl concentration. The [NiFe] catalytic site was modelled into Mbh using the high resolution soluble hydrogenase (PDB ID: 4U9H) (35). Initial protonation states were obtained from Poisson-Boltzmann electrostatic calculations with Monte Carlo sampling (3), based on models with the cofactors treated in their oxidized state (see SI Appendix, Table S24, for a list of residues with non-standard protonation state). All MD simulations were performed in an *NPT* ensemble at *T* = 310 K and *p* = 1 atmosphere, using a 2 fs integration timestep, and with electrostatics modelled using the particle mesh Ewald (PME) method. The system was gradually relaxed for 10 ns with harmonic restraints of 2 kcal mol^-1^ Å^-1^ on all protein and cofactor heavy atoms followed by 10 ns with harmonic restraints of 2 kcal mol^-1^ Å^-1^ on all protein backbone heavy atoms and by 10 ns equilibration with weak (0.5 kcal mol^-1^ Å^-1^) restraints on all Cα atoms, and 0.5 μs production runs. All classical MD simulations (3.5 μs in total) were performed using NAMD2 (v. 2.12/2.13) for equilibration and NAMD3 (61) for production runs. The simulations were analyzed using VMD (62). Multiple sequence alignment (MSA) for soluble hydrogenases and complex I superfamily was done with ClustalW (63) and visualized with Jalview (64).

## Acknowledgements

A.S. acknowledges the EMBO Long-Term Fellowship (ALTF 952-2022) for support. This work was supported by European Research Council under the European Union’s Horizon 2020 research and innovation program/grant agreement 715311 (V.R.I.K.), the Knut and Alice Wallenberg Foundation (V.R.I.K. grant: 2019.0251), and the Swedish Research Council (V.R.I.K.). V.R.I.K. also acknowledges DFG for support within the Mercator Fellow Program to SFB1078. This work was also supported by the National Academic Infrastructure for Supercomputing in Sweden (NAISS 2023/1-31) and the Swedish National Infrastructure for Computing (SNIC 2022/1-29 and SNIC 2022/13-14) at the Center for High Performance Computing (PDC) Center, partially funded by the Swedish Research Council through a grant agreement no. 2018-05973 and the Leibniz Rechenzentrum (LRZ, project:pr83ro), Germany.

## Authors contribution

V.R.I.K. and A.S. designed research; A.S. and A.P.G.H. performed research; A.P.G.H. developed new methods: A.S., A.P.G.H., and V.R.I.K analyzed results; A.S. and V.R.I.K. wrote the manuscript with contributions from all authors.

## Data, Materials, and Software Availability

Cartesian coordinates of all quantum cluster DFT models (115 structures) are deposited in the Zenodo repository (10.5281/zenodo.10823885). The force field parameters for the [NiFe] center are available at https://github.com/KailaLab/ff_parameters.

## Table of contents (TOC) graphics

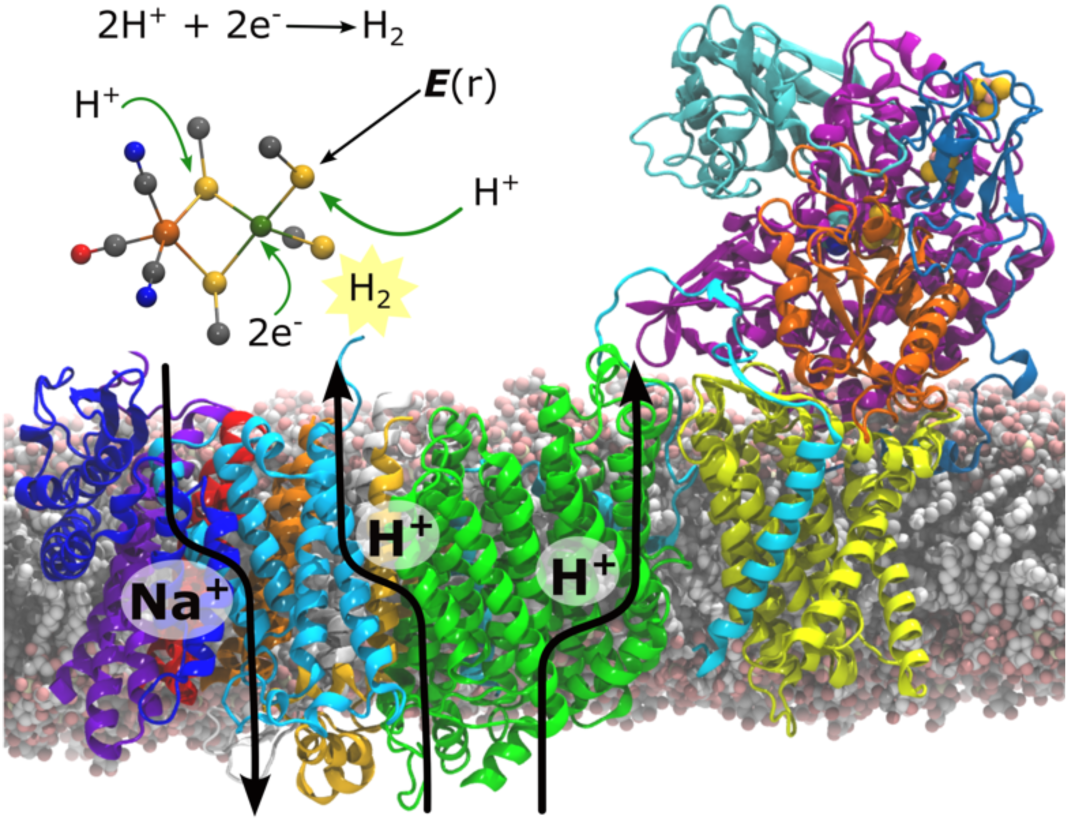

## Notes

https://zenodo.org/doi/10.5281/zenodo.10823884

### Competing Interest Statement

The authors have declared no competing interest.

