## Supplementary material for "Mechanistic principles of hydrogen evolution in the membrane-bound hydrogenase": SI Appendix

#### Table of Contents

##### Extended Methods

1. Benchmarking the [NiFe] geometry, spin states, and reaction energetics
    1.1 Assessment of DFT functionals for the [NiFe] active site geometry
    1.2 Evaluation of spin energetics
    1.3 Evaluation of reaction energetics

##### List of Figures

**Figure S1.** Cluster models for benchmarking the DFT methodology and comparison to high resolution x-ray data.
**Figure S2.** Benchmarking the quantum chemical methodology for spin state energetics.
**Figure S3.** Benchmarking the quantum chemical methodology for reactions energetics.
**Figure S4.** DFT models of Mbh.
**Figure S5.** DFT models of the Ni-$SI_a$ state with low spin Ni(II).
**Figure S6.** DFT models of the Ni-$SI_a$ state with high spin Ni(II).
**Figure S7.** Reaction energy profiles for water-mediated proton transfer from $Glu21_L$ to $Cys374_L$ in the Ni-$SI_a$ state.
**Figure S8.** DFT models of the Ni-L and Ni-C states.
**Figure S9.** Reaction energy profiles for water-mediated proton transfer from $Glu21_L$ to $Cys374_L$ and hydride binding step involving protonated $Cys374_L$ and $Ni^{III}$.
**Figure S10.** DFT models for the Ni-C catalytic state for the second proton transfer step.
**Figure S11.** Reaction energy profiles for water-mediated proton transfer from $Glu21_L$ to $Cys374_L$ in the Ni-C state.
**Figure S12.** DFT models of the Ni-R state with low spin Ni(II).
**Figure S13.** DFT models of the Ni-R state with high spin Ni(II).
**Figure S14.** Reaction energy profile for proton transfer via the His pathway during $H_2$ formation in the LS and HS forms.
**Figure S15.** Reaction energy profile for proton transfer via the Glu pathway during $H_2$ formation in the LS and HS forms.
**Figure S16.** Combined reaction energy profile for proton transfer along the His and Glu pathways in the LS and HS forms.
**Figure S17.** DFT models of the Ni-$SI_a$ state with low spin Ni(II) in the presence of $His75_L^+$.
**Figure S18.** DFT models of the Ni-$SI_a$ state with high spin Ni(II) in the presence of $His75_L^+$.
**Figure S19.** Reaction energy profiles for water-mediated proton transfer from $Glu21_L$ to $Cys374_L$ in the Ni-$SI_a$ state in the presence of $His75_L^+$.
**Figure S20.** DFT models of the Ni-L and Ni-C states in the presence of $His75_L^+$.
**Figure S21.** Reaction energy profiles for water-mediated proton transfer from $Glu21_L$ to $Cys374_L$ and hydride binding step involving protonated $Cys374_L$ and $Ni^{III}$ in the presence of $His75_L^+$.
**Figure S22.** DFT models for the Ni-C catalytic state for the second proton transfer step in presence of $His75_L^+$.
**Figure S23.** DFT models of the Ni-R state with low spin Ni(II) in the presence of $His75_L^+$.
**Figure S24.** DFT models of the Ni-R state with high spin Ni(II) in the presence of $His75_L^+$.
**Figure S25.** Reaction energy profile for the proton originating from the "Glu" pathway for the LS and HS forms in the presence of $His75_L^+$.
**Figure S26.** Electric field effects on the proton transfer in the Ni-R state and their influence on the protonation state of $His75_L$.
**Figure S27.** Conformational dynamics of key residues involved in the loop network.
**Figure S28.** Conformational sampling of loops involved in the network.
**Figure S29.** Multiple sequence alignment of MbhM.
**Figure S30.** Multiple sequence alignment of MbhL.
**Figure S31.** Multiple sequence alignment of MbhI.
**Figure S32.** Multiple sequence alignment of MbhJ.

#### List of Tables


**Table S1.** Key bond distances of DFT optimized [NiFe] active site from the soluble [NiFe] hydrognease and comparison to a high-resolution x-ray structure.

**Table S2.** Mulliken spin population of key atoms in the Ni-R state of the [NiFe] soluble hydrogenase with various DFT functionals.

**Table S3.** Comparison of adiabatic singlet-minus-triplet (S-T) gaps for the soluble [NiFe] hydrogenase obtained with various DFT functionals and wavefunction-based approaches.

**Table S4.** Reaction energetics of the $H_2$ formation step in Mbh obtained with various DFT functional and wavefunction-based approaches.

**Table S5.** Adiabatic spin state gap between low and high spin of the Ni-$SI_a$ models of Mbh.

**Table S6.** Key bond distances of the DFT optimized Ni-$SI_a$ state of Mbh.

**Table S7.** Mulliken spin populations of key atoms in the Ni-$SI_a$ state of Mbh.

**Table S8.** Key bond distances of the DFT optimized Ni-L and Ni-C states of Mbh.

**Table S9.** Mulliken spin populations of the key atoms in the Ni-L and Ni-C states of Mbh.

**Table S10.** Adiabatic spin state gap between low and high spin form of the Ni-R state of Mbh.

**Table S11.** Key bond distances of the DFT optimized Ni-R state of Mbh.

**Table S12.** Mulliken spin populations of the key atoms in the Ni-R state of Mbh.

**Table S13.** $H_2$ binding free energy in the Ni-R state of Mbh.

**Table S14.** Adiabatic spin state gap between low and high spin form of the Ni-$SI_a$ models in the presence of $His75_L^+$.

**Table S15.** Key bond distances of the DFT optimized Ni-$SI_a$ state in the presence of $His75_L^+$.

**Table S16.** Mulliken spin population of key atoms in the Ni-$SI_a$ state in the presence of $His75_L^+$.

**Table S17.** Key bond distances of the DFT optimized Ni-L and Ni-C states in the presence of $His75_L^+$.

**Table S18.** Mulliken spin populations of the key atoms in the Ni-L and Ni-C state in the presence of $His75_L^+$.

**Table S19.** Adiabatic spin state gap between low and high spin forms of the Ni-R state in the presence of $His75_L^+$.

**Table S20.** Key bond distances of the DFT optimized Ni-R state in the presence of $His75_L^+$.

**Table S21.** Mulliken spin populations of key atoms in the Ni-R state in the presence of $His75_L^+$.

**Table S22.** Redox states of the [NiFe] active site in MD simulations.

**Table S23.** Complete list of classical MD simulations.

**Table S24.** List of non-standard protonation states in the MD simulations.


#### List of Movies


**Movie S1.** Catalytic progression during the Ni-$SI_a$ → Ni-L transition.

**Movie S2.** Catalytic progression during the Ni-L → Ni-C transition.

**Movie S3.** $H_2$ formation in the Ni-R state.

### 1. Benchmarking the [NiFe] geometry, spin states, and reaction energetics

#### 1.1 Assessment of DFT functionals for the [NiFe] active site geometry

To test the performance of different density functionals for describing the geometry of the [NiFe] active site, we constructed quantum chemical model based on the x-ray structure of the soluble [NiFe] hydrogenase in the Ni-R state resolved at 0.89 Å (PDB ID: 4U9H) (1). Importantly, the experimentally resolved positions of the protons makes this structure ideal for the quantum chemical benchmarking purposes. To this end, we constructed a large quantum chemical cluster model, comprising of catalytic [NiFe] core, the hydride, two $CN^-$ and one CO ligands, together with all residues in the first and second coordination sphere (including Thr18, Glu34, Cys81, Gly82, Val83, Cys84, His88, Ser92, Asp123, Ala477, Pro478, Arg479, His484, Val500, Pro501, Ser502, Asp544, Cys546, Ile547, Ala548, Cys549 (**Figure S1**). The protons resolved in the X-ray structure were kept in our model, leading to a system with 243 atoms. Truncated backbone atoms were terminated by methyl groups, whereas the $C_\alpha$ atoms of all residues were fixed at their X-ray position during the geometry optimization.

The quantum cluster model (**Figure S1**) were optimized at four different rungs along the "Jacobs Ladder", including BP86 (GGA)(2), TPSS (meta-GGA) (3), TPSSh (hybrid meta-GGA) (4) and at the B3LYP (hybrid GGA) (5) level. During the geometry optimization, the central core of the active site comprising the Fe, Ni, S, $2CN^-$ and CO were assigned def2-TZVP basis sets (6), whereas the other atoms were described at the def2-SVP level. We also performed geometry optimization using scalar-relativistic X2C Hamiltonian (exact two-component) (7-9) and corresponding basis sets (x2c-TZVPall and x2c-SVPall). The multipole accelerated resolution of identity (MARI-J) approximation (10) was used during the geometry optimizations to lower the computational cost. Dispersion effects were included using the D3-BJ approximation,(11) together with a dense DFT integration grids (m4) and tight SCF convergence thresholds (scfconv 8) used during the optimizations. The surroundings beyond the large explicit model were described with the implicit solvation COSMO model using a dielectric constant of 4.0. The iron and nickel were modeled in the +2 oxidation state, with the models optimized in both the low spin (S=0) and high-spin (S=1) configurations. All the calculations were performed using TURBOMOLE v. 7.5.1 (12).

Key geometric parameters obtained with different density functionals are listed in **Table S1**. All tested DFT functionals gave reasonable predictions of the metal-ligand distances, and we find that inclusion of relativistic effects does not improve the results. Our structural assessment of the [NiFe] crystal structure suggests that some of the key bridging ligands are asymmetrically bound with the Ni and Fe atoms, *e.g.*, the bond between Ni-Cys549 is longer than the Fe-Cys549 bond, while the bond between Ni-$H^-$ is shorter than the Fe-$H^-$ bond. Interestingly, we find that the structures optimized in the low spin configuration, irrespective of the employed density functional, reproduced this trend, whereas the HS structures show opposite trend **(Tables S1 and S2)**. Our analysis thus reveals that the resolved high-resolution x-ray structure of the Ni-R state is a low spin (singlet) ground state configuration. In the next step, we benchmarked how different DFT methods and wavefunction based approaches estimate the spin energetics.

##### 1.2 Evaluation of spin state energetics

The divalent nickel ion of the [NiFe] center takes singlet (S=0, low-spin, diamagnetic) or triplet (S=1, high-spin, paramagnetic) configuration. Low-spin $Ni^{II}$ complexes often prefer a square-planar coordination environment, whereas high spin $Ni^{II}$ complexes prefer a tetrahedral coordination. For the high-resolution x-ray structure of [NiFe] hydrogenase from *Desulfovibrio vulgaris Miyazaki F*, the coordination of $Ni^{II}$ is square-pyramidal (1), hence indicative of a low-spin state.

However, there is not consensus about the ground state spin state of the Ni-$SI_a$ and Ni-R states. Experimental studies using EPR (13), saturation magnetization (14) and UV-visible MCD investigations (15) suggest that these states reside in singlet ground state, whereas EXAFS (16) and some DFT studies suggests that the triplet ground state is preferred (17). Our geometric analysis (See **Section 1.1**) show that the structure optimized in the $Ni^{II}$ low spin form accurately reproduce the structural features of the high-resolution structure.

To probe the spin energetics of the [NiFe] active site, we next performed DFT calculations with different functionals. It is well known that that preferred spin configuration, correlated with the amount of exact exchange (HFX) present in the functional, with hybrid functional stabilizing the HS states, whereas pure functionals tend to stabilize the LS state. In this regard, Reiher and co-workers (18) found that 8-15% often provide accurate description of the spin energetics for bio-inorganic complexes (*cf.* also Refs. (19-21)

To study how different DFT methods affect the single-triplet splitting (S-T) gaps for the NiFe center, we constructed smaller quantum chemical models **(Figure S2A)** consisting of Ni, Fe, hydride, 2CN, 1CO and 4 cysteine residues based on the optimized large model systems **(Figure S1)**. After optimization at the TPSSh/def2-SVP/def2-TZVP level (see **Section 1.1**), we performed single point calculations on these structures to determine the S-T gaps using various DFT functionals **(Figure S2B)** and compared these to data obtained with the Random Phase Approximation (RPA) (22) and DLPNO-CCSD($T_1$) (23), which provide an accurate description for spin-state energetics when benchmarked against CCSD(T) and CASSCF/CASPT2. (24)

The DFT and DLPNO-CCSD($T_1$) calculations were performed with the ORCA5.0 program suite.(25) The DFT calculations were performed at the BP86, TPSS, TPSSh, B3LYP, B3LYP* (15% HFX) and B3LYP** (10% HFX) levels. The relativistic effects were included using the Douglas-Kroll-Hess (DKH) Hamiltonian of order 2 (DKH2) (26, 27) with DKH-def2-TZVPP basis sets on Ni, Fe, and DKH-def2-TZVP on the remaining atoms, together with tight SCF convergence and high DFT integration grid (verytightscf and defgrid3 in the ORCA).

The DLPNO-CCSD($T_1$) calculations were based on the DFT/ TPPSh solution as a reference guess for the initial wave function. For the open-shell systems, Quasi-restricted orbitals (QRO) obtained from unrestricted Kohn-Sham (UKS) calculations were used as reference. Moreover, we used NormalPNO setting (with $T_{CutPairs}$ set to $10^{-5}$) due to large system size and corresponding large number of PNOs. The 1s, 2s and 2p electrons in Fe and Ni and 1s electrons of S were kept frozen (total of 28 electrons). All DFT and DLPNO-CCSD($T_1$) computations were performed with the CPCM ($\varepsilon$=4).

RPA calculations (based on the TPSSh reference determinant) were performed with Turbomole 7.5.1. (12) Scalar relativistic effects were included using DKH2 Hamiltonian. Fe and Ni were assigned x2c-TZVPPall basis-set and x2c-TZVPall on rest of the atoms. Implicit solvation in case of RPA was incorporate using the COSMO model ($\varepsilon$=4). All computations were performed using resolution of identity approximation (RI-J) and corresponding basis-sets. Non-relativistic calculations in case of DFT, RPA and DLPNO-CCSD($T_1$) were performed with def2-TZVPP (Fe and Ni) and def2-TZVP (C, N, O, S and H atoms) basis-sets.

A detailed comparison of the S-T gaps is provided in **Figure S2 and Table S3.** We note that wavefunction based approaches predict low-spin form (singlet) to more stable compared to the high-spin form (triplet), which agrees well with the results from geometrical analysis **(Section 1.1)**.

##### 1.3 Evaluation of reaction energetics

The $H_2$ formation steps in low-spin configuration of $Ni^{II}$ (S=0) was used for benchmarking the reaction energetics using different DFT functionals against RPA and DLPNO-CCSD(T) calculations. The structures of the reactant, transition state and product were derived from the reaction-optimization pathway computation performed on the large model. The system size was reduced by deleting residues in the second sphere and shortening the sidechains. The trimmed models contained 154 atoms **(see Figure S3A)**.

For the DFT and DLPNO-CCSD(T) calculations, we used DKH-def2-TZVPP on Fe, Ni and DKH-def2-TZVP on sulfur atoms and DKH-def2-SVP on rest of the atoms. Very tight SCF convergence (verytightscf in the ORCA convention) and higher DFT integration grid (defgrid3 in the ORCA convention) was used. In case of DLPNO-CCSD(T) calculations, we used DFT determinant (TPSSh functional) as a reference. We used "NormalPNO" setting (with $T_{CutPairs}$ set to $10^{-5}$) due to large system size and corresponding large number of PNOs. The relativistic effects were included using the DKH2 Hamiltonian. The 1s, 2s and 2p electrons in Fe and Ni and 1s electrons of S were kept frozen (total of 28 electrons). All DFT and DLPNO-CCSD(T) computations were performed with the CPCM (ε=4) module implemented in ORCA 5.0.(25)

The RPA calculations (with reference TPSSh determinant) were performed using Turbomole 7.5.1. Scalar relativistic effects were included using DKH2 Hamiltonian. Iron and nickel were assigned x2c-TZVPPall basis-set, x2c-TZVPall for sulphur and x2c-SVPall for rest of the atoms. Implicit solvation in case of RPA was incorporated using the COSMO model (ε=4). All computations were performed using resolution of identity approximation (RI-J) and corresponding basis-sets. The relative electronic energies of the reactant (R), transition state (TS) and product (P) are reported in the **Figure S3B and Table S4.**

#### Supplementary figures

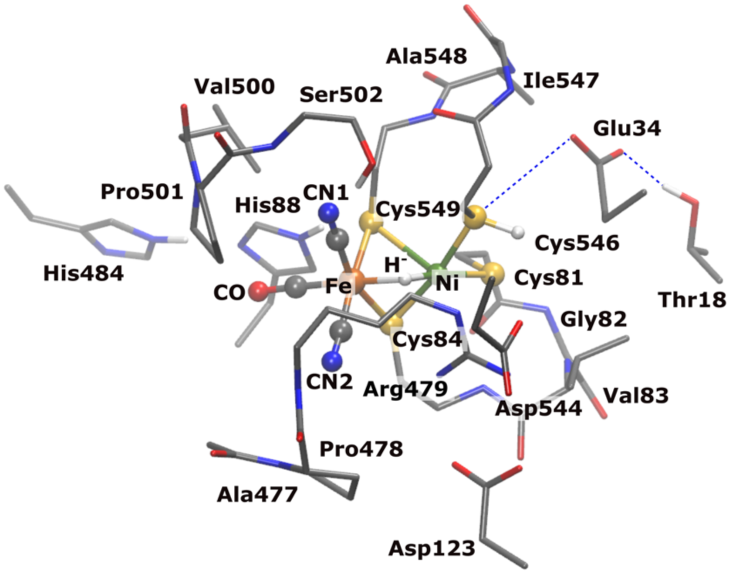


**Figure S1. Cluster models for benchmarking the DFT methodology and comparison to high resolution x-ray data.** The model was built from the soluble [NiFe] hydrogenases (*Desulfovibrio vulgaris Miyazaki F*) resolved at 0.89 Å resolution (PDB ID: 4U9H)(1). The model contains a total of 243 atoms, and optimized using different density functionals (see Table S1).

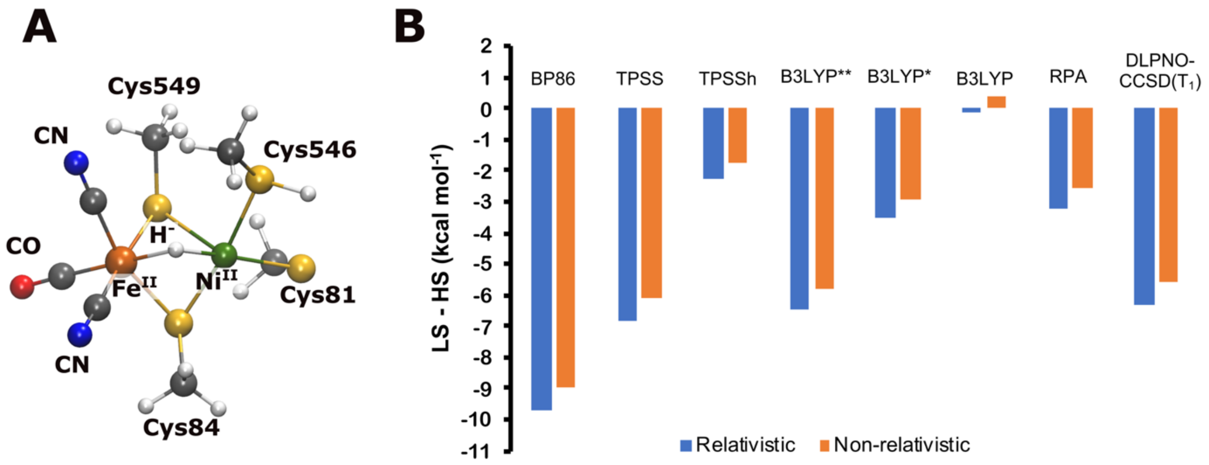


**Figure S2. Benchmarking the quantum chemical methodology for spin-state energetics.** (A) Minimalistic model of [NiFe] cluster (Ni-R state) used for benchmarking spin-state energetics. These models were derived from the corresponding low-spin and high-spin optimized structures (TPSSh functional) of the large-models (see section 1 and Figure S1). (B) Comparison of the Adiabatic singlet-minus-triplet (S-T) gaps for the [NiFe] clusters obtained from various DFT functionals and wavefunction based approaches (RPA and DLPNO-CCSD($T_1$)) using both relativistic and non-relativistic approaches. Absolute values are reported in the Table S3. B3LYP** and B3LYP* are with 10% and 15% exact-exchange, respectively.

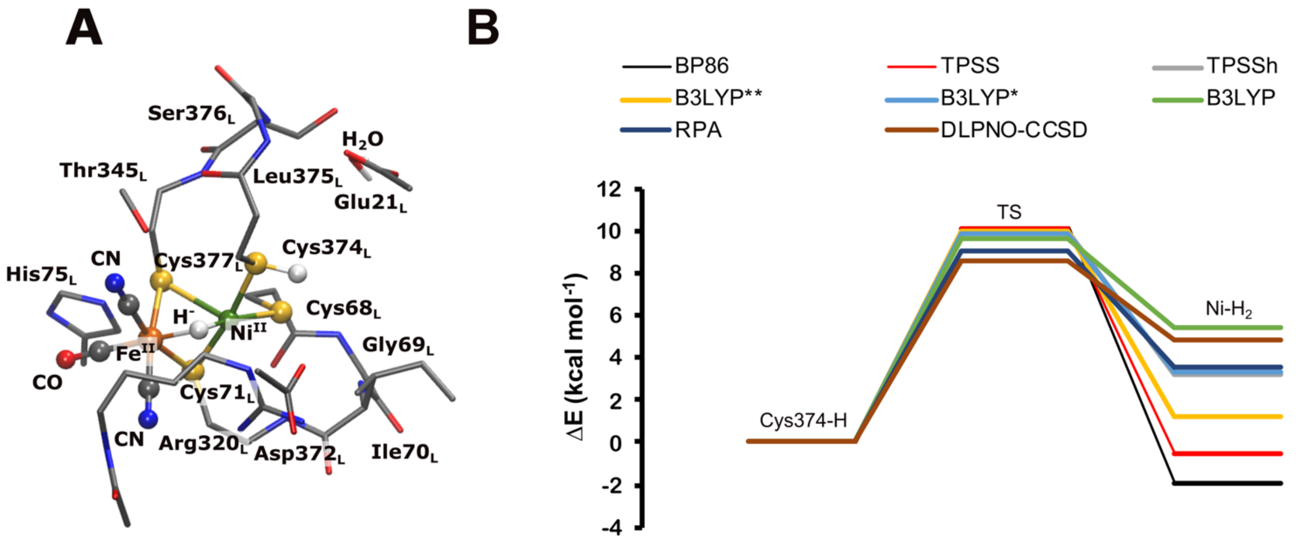


**Figure S3. Benchmarking the quantum chemical methodology for reaction energetics.** (A) Reduced-size cluster model of $H_2$ bound [NiFe] cluster for benchmarking reaction energetics. Corresponding trimmed structures were also created for the reactant and transition states. Reactive hydrogen atoms, that forms $H_2$ eventually, are drawn with larger vDW radius. (B) Comparison of performance of several quantum chemical methods in prediction of the reaction energetics of the $H_2$ formation. B3LYP** and B3LYP* are with 10% and 15% exact-exchange, respectively. Absolute values are provided in Table S4.

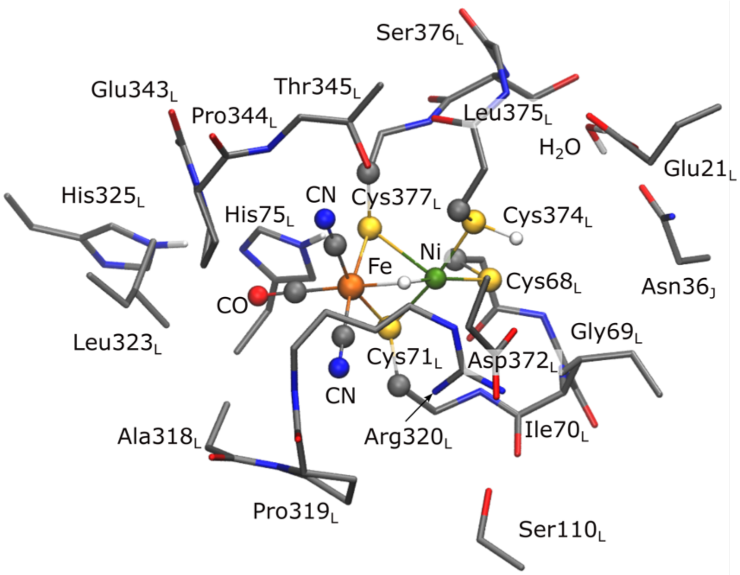


**Figure S4. DFT models of Mbh.** The quantum cluster model derived from membrane bound hydrogenase (Mbh) from *Pyrococcus furiosus*. The model contains up to total of 264 atoms. All geometry optimization computations in this work were performed using this model. The single-point energy computations were performed with the exclusion of $Asn36_J$ residue.

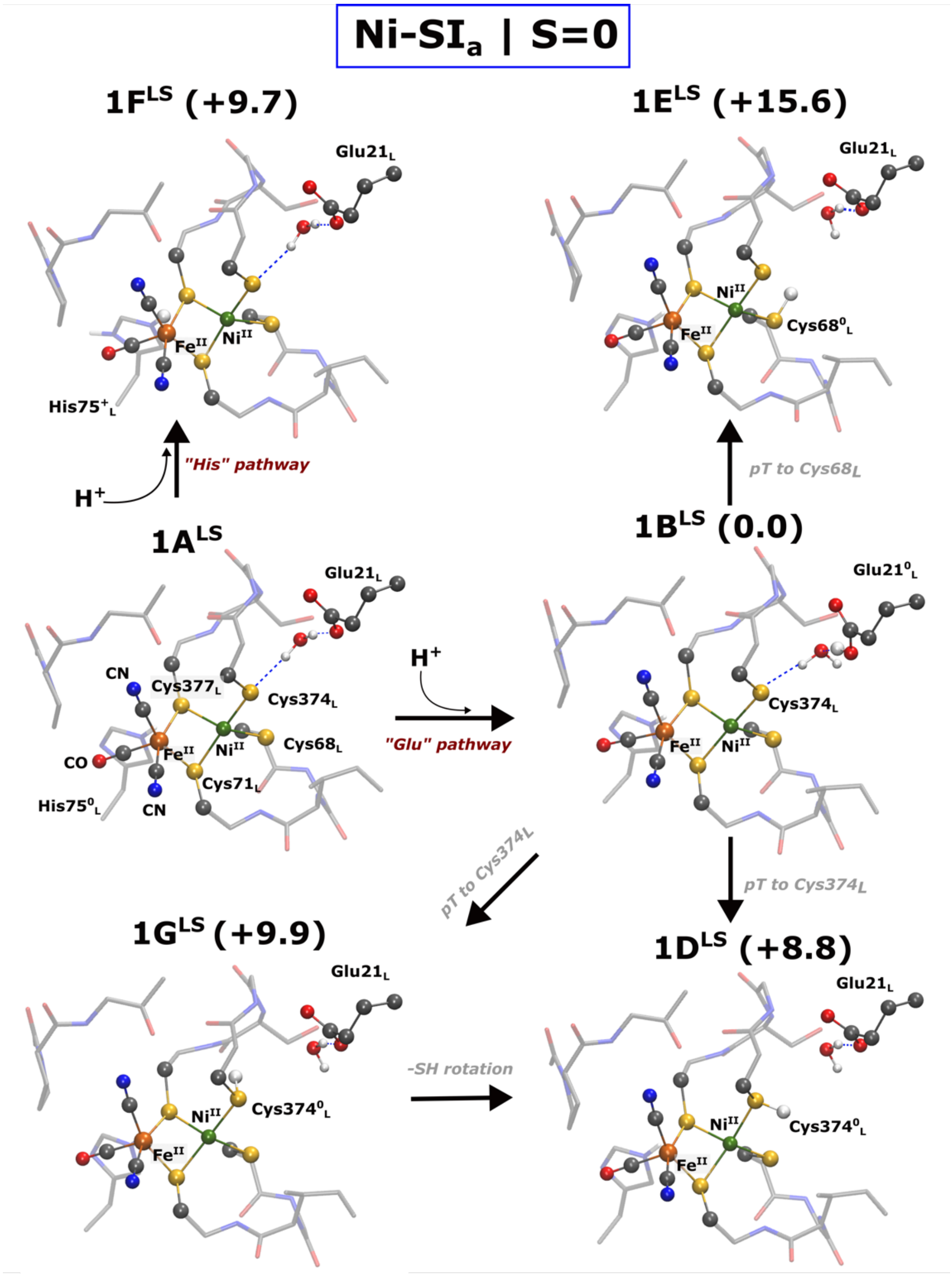


**Figure S5. DFT models of the Ni-SI$_a$ state with low-spin Ni(II).** Core structure of the DFT optimized models of [NiFe] active-site modelled in the Ni-SI$_a$ state (Ni$^{II}$Fe$^{II}$). The Ni$^{II}$ is modeled in low-spin configuration with S=0. Relative electronic energy (in kcal mol$^{-1}$) computed at B3LYP*/x2c-TZVPP/x2c-TZVP level of theory is also depicted. Attempts to protonate Cys377$_L$ resulted in proton jumping back to the His75$_L$. Only the core structure is depicted along with key residues. Key bond distances for all models are shown in Table S6.

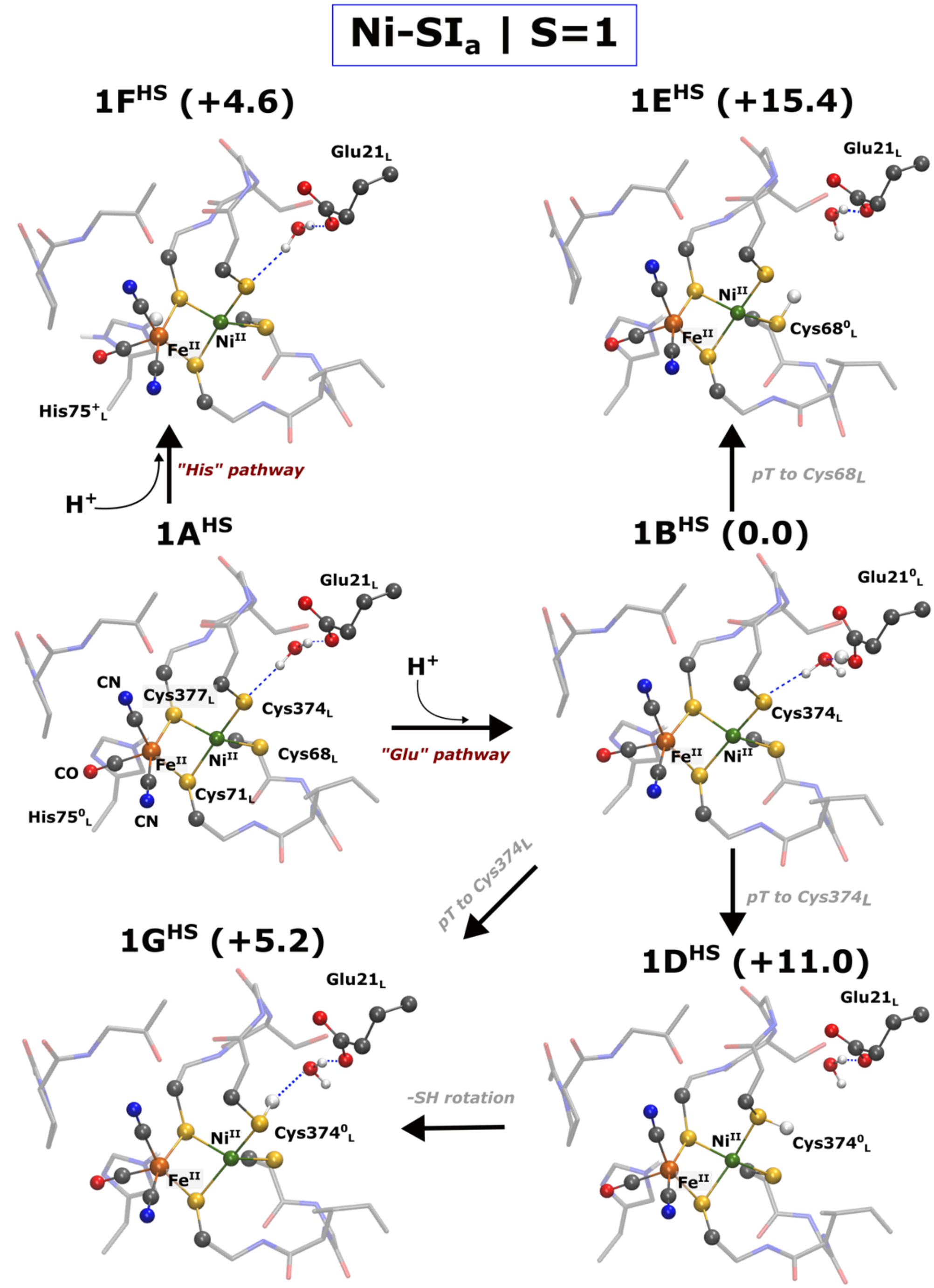


**Figure S6. DFT models of the Ni-SI$_a$ state with high-spin Ni(II).** Core structure of the DFT optimized models of [NiFe] active-site modelled in the Ni-SI$_a$ state (Ni$^{II}$Fe$^{II}$). The Ni$^{II}$ is modeled in high-spin configuration with S=1. Relative electronic energy (in kcal mol$^{-1}$) computed at B3LYP*/x2c-TZVPP/x2c-TZVP level of theory is also depicted. Attempts to protonate Cys377$_L$ resulted in proton jumping back to the His75$_L$. Only the core structure is depicted along with key residues. Key bond distances and Mulliken spin-populations for all models are shown in Table S6 and Table S7, respectively.

A

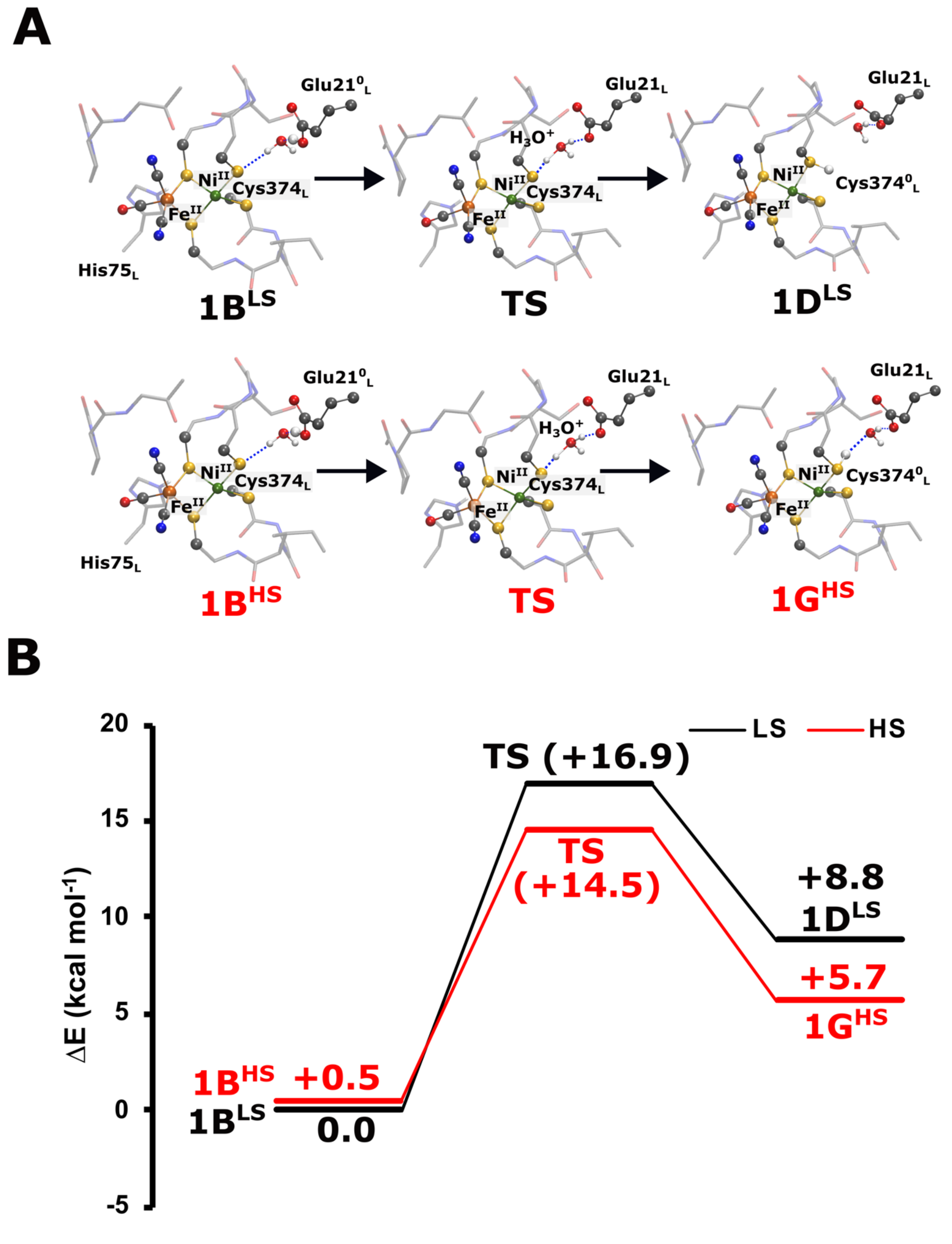


**Figure S7. Reaction energy profiles for water-mediated proton transfer from Glu21$_L$ to Cys374$_L$ in the Ni-SI$_a$ state.** The complete reaction pathway was investigated in both low-spin (LS, black line) and high-spin configurations (HS, red line). The structures were optimized in both LS and HS forms. Single-point energy computations were performed with B3LYP*/x2c-TZVPPall/x2c-TZVPall level of theory. The reactant, product and transition state structures corresponding to both spin-state are depicted.

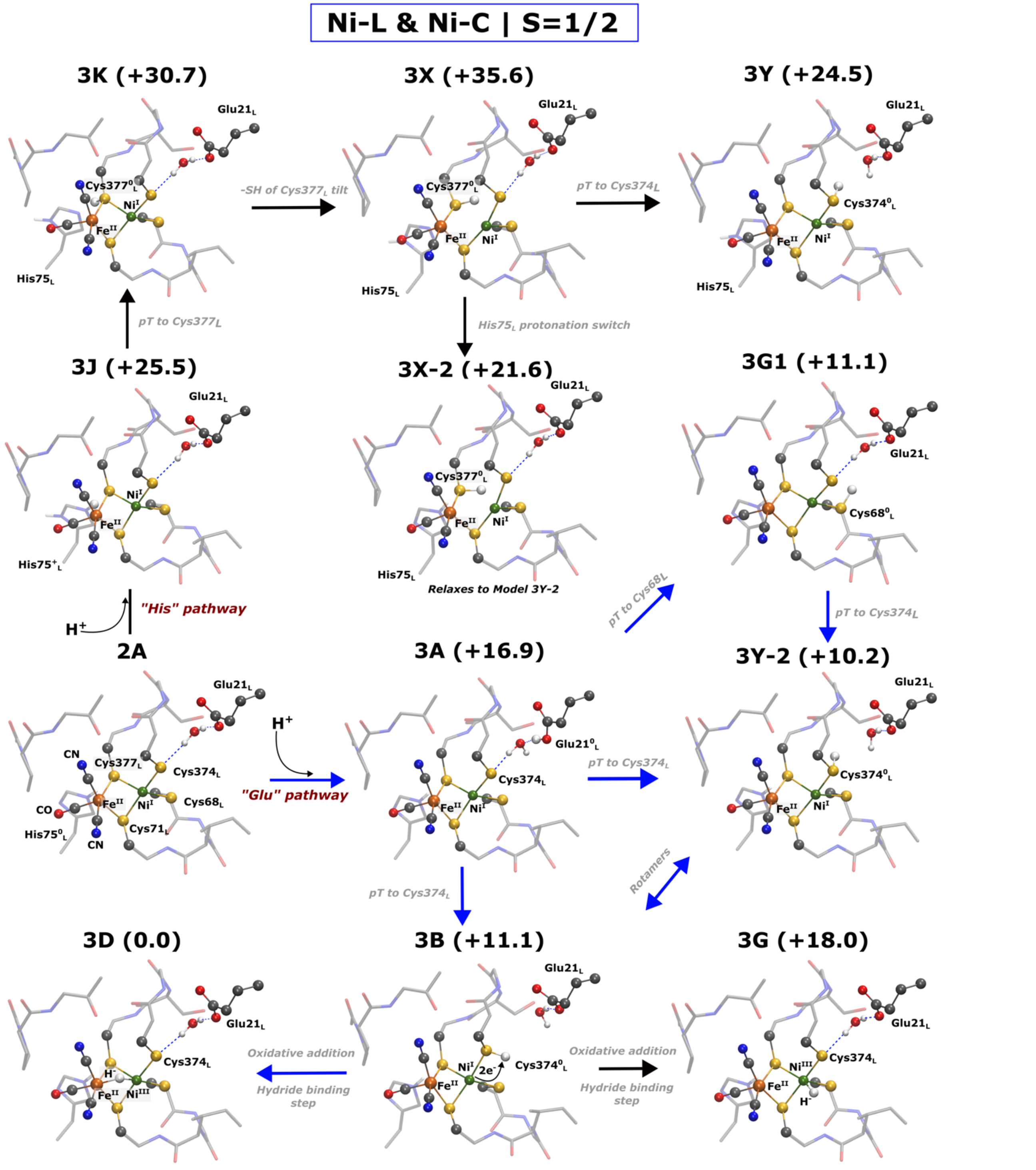


**Figure S8. DFT models of the Ni-L and Ni-C states.** Core structure of the DFT optimized models of [NiFe] active-site modelled in the Ni-L ($Ni^{I}Fe^{II}$) and Ni-C state ($Ni^{III}Fe^{II}$). In both states, $Ni^{I/III}$ has S=1/2. Relative electronic energy (in kcal $mol^{-1}$) computed at B3LYP*/x2c-TZVPP/x2c-TZVP level of theory is also depicted. Blue and black arrows depict exergonic and endergonic reaction progression. Only the core structure is depicted along with key residues. Key bond distances for all models and Mulliken spin-populations are shown in Table S8 and Table S9, respectively.

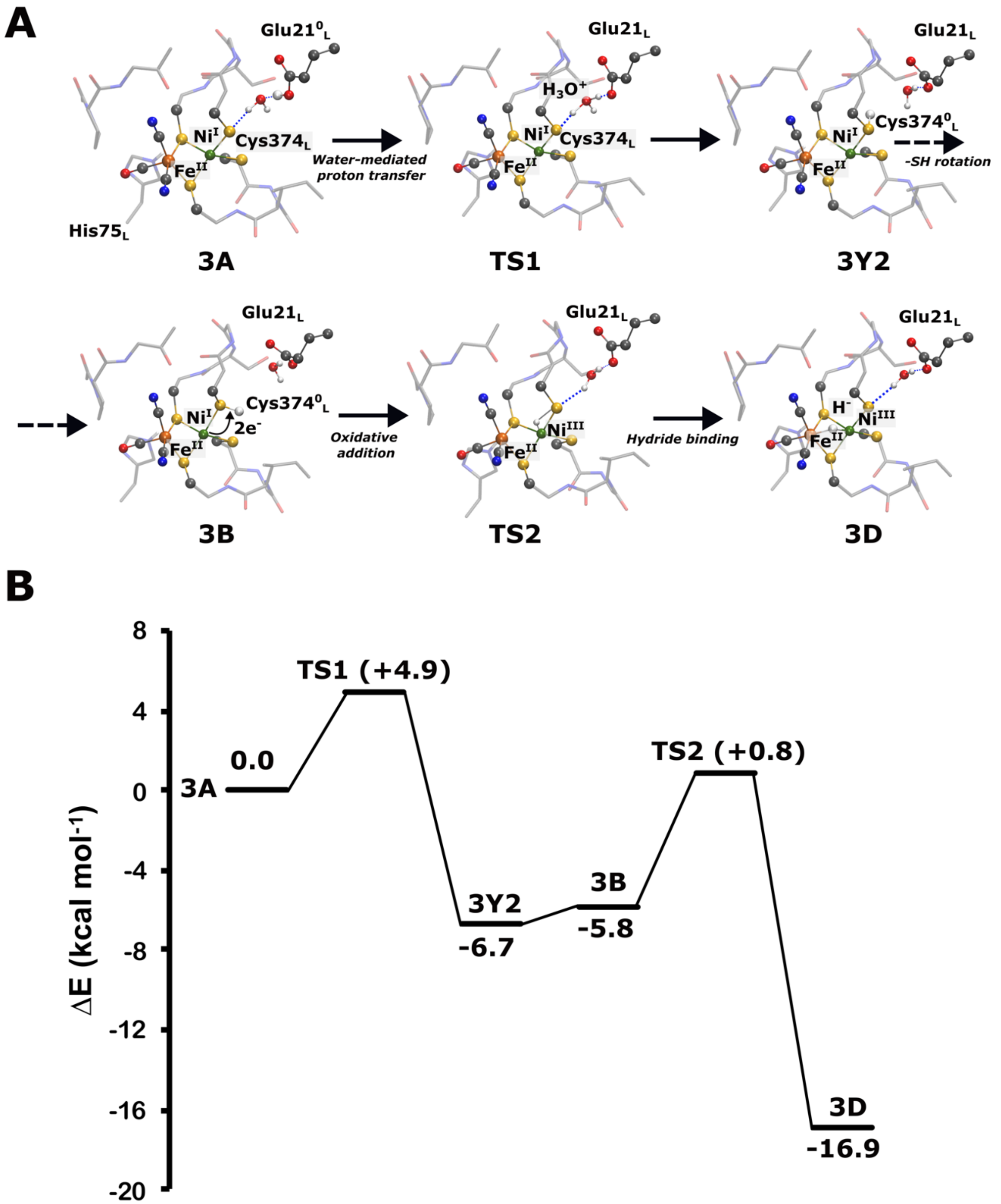


**Figure S9. Reaction energy profiles for water-mediated proton transfer from Glu21$_L$ to Cys374$_L$ and hydride binding step involving protonated Cys374$_L$ and Ni$^{III}$.** Single-point energy computations were performed with B3LYP*/x2c-TZVPPall/x2c-TZVPall level of theory. The reactant, product and transition state structures are also depicted.

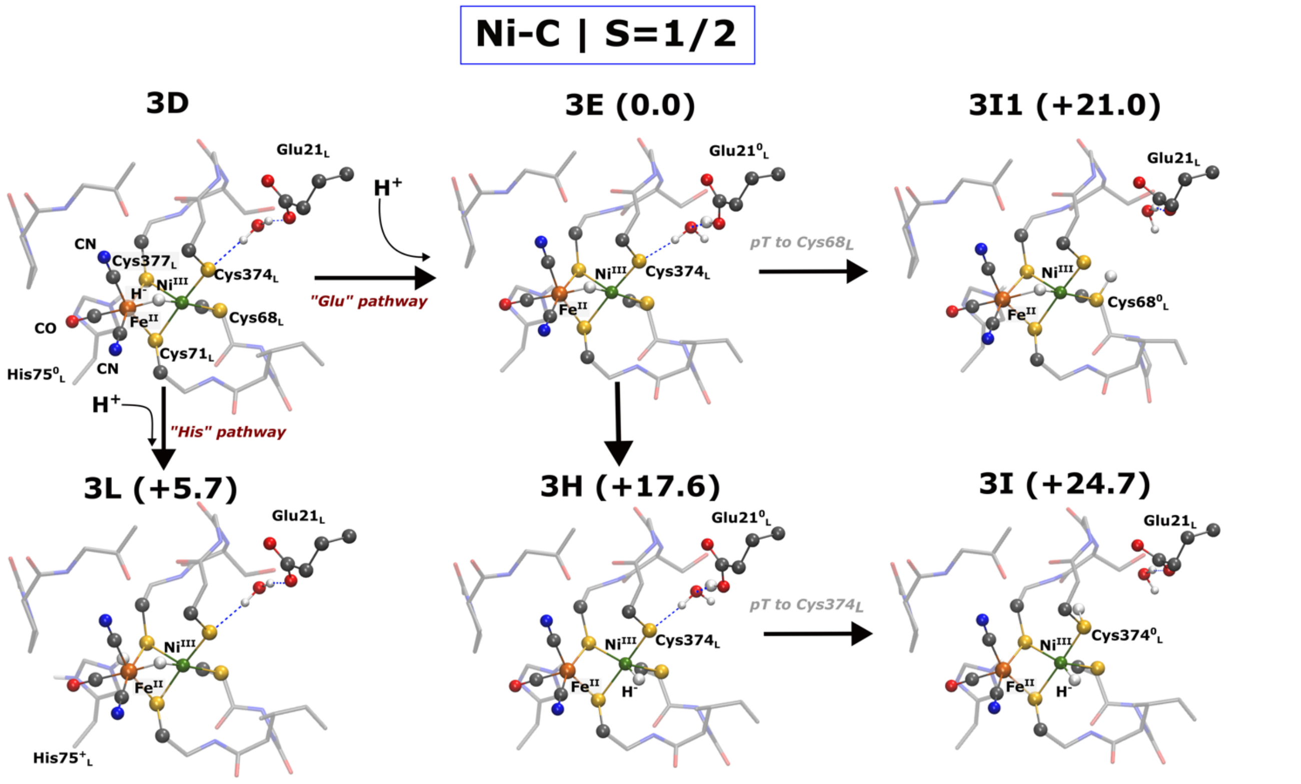


**Figure S10. DFT models for the Ni-C catalytic state for the second proton transfer step.** Core structure of the DFT optimized models of [NiFe] active-site modelled Ni-C state ($Ni^{III}Fe^{II}$, S=1/2). Relative electronic energy (in kcal $mol^{-1}$) computed at B3LYP*/x2c-TZVPP/x2c-TZVP level of theory is also depicted. Only the core structure is depicted along with key residues. Key bond distances for all models and Mulliken spin-populations are shown in Table S8 and Table S9, respectively. Attempts to protonate $Cys377_L$ resulted in proton jumping back to the $His75_L$.

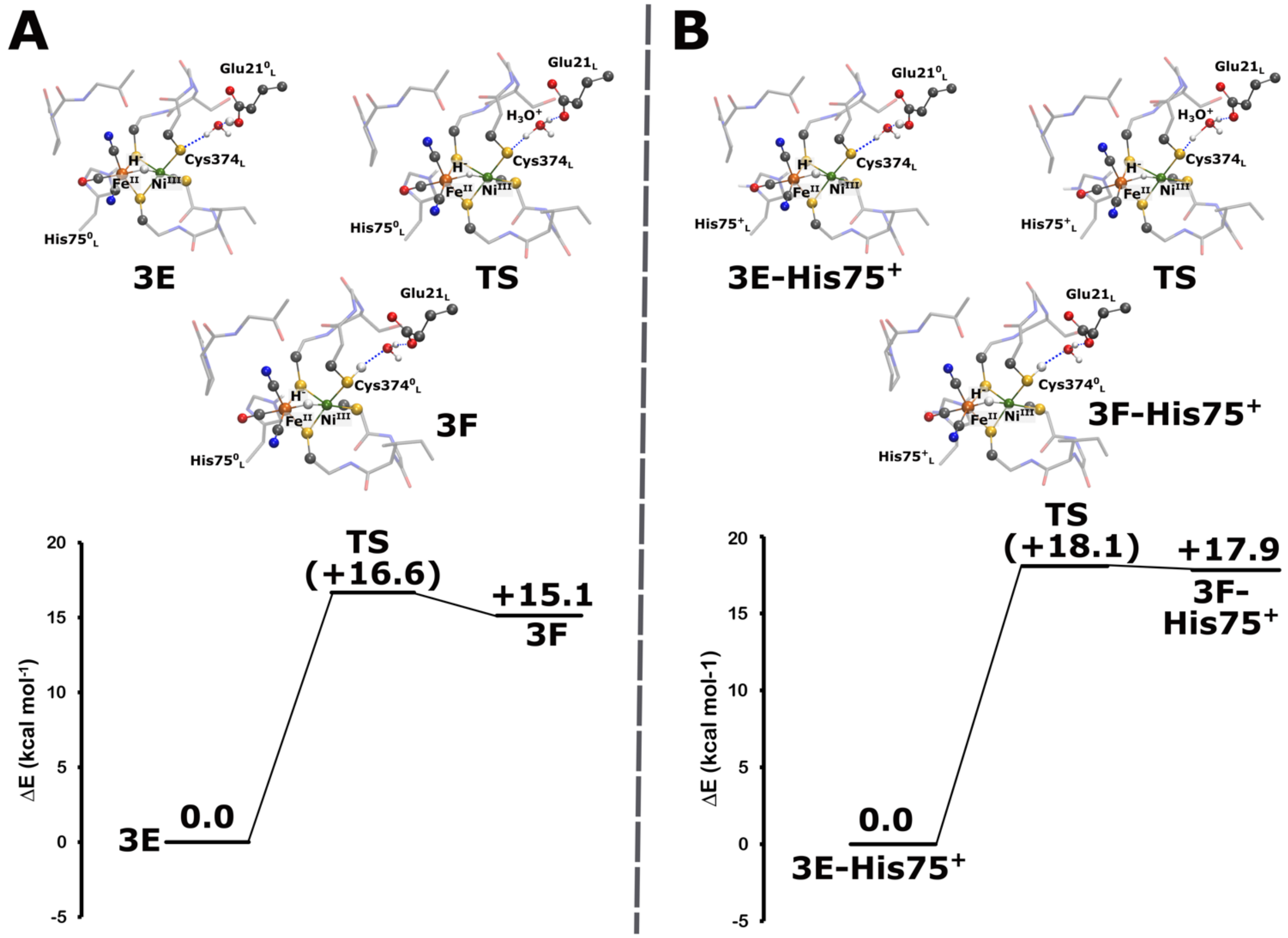


**Figure S11. Reaction energy profiles for water-mediated proton transfer from Glu21$_L$ to Cys374$_L$ in the Ni-C state.** Energetics of water-mediated proton transfer from Glu21$_L$ to Cys374$_L$ in the Ni-C state in the presence of (A) neutral and (B) protonated His75$_L$. Single-point energy computations were performed at the B3LYP*/x2c-TZVPPall/x2c-TZVPall level of theory. The reactant, product and transition state structures are also depicted.

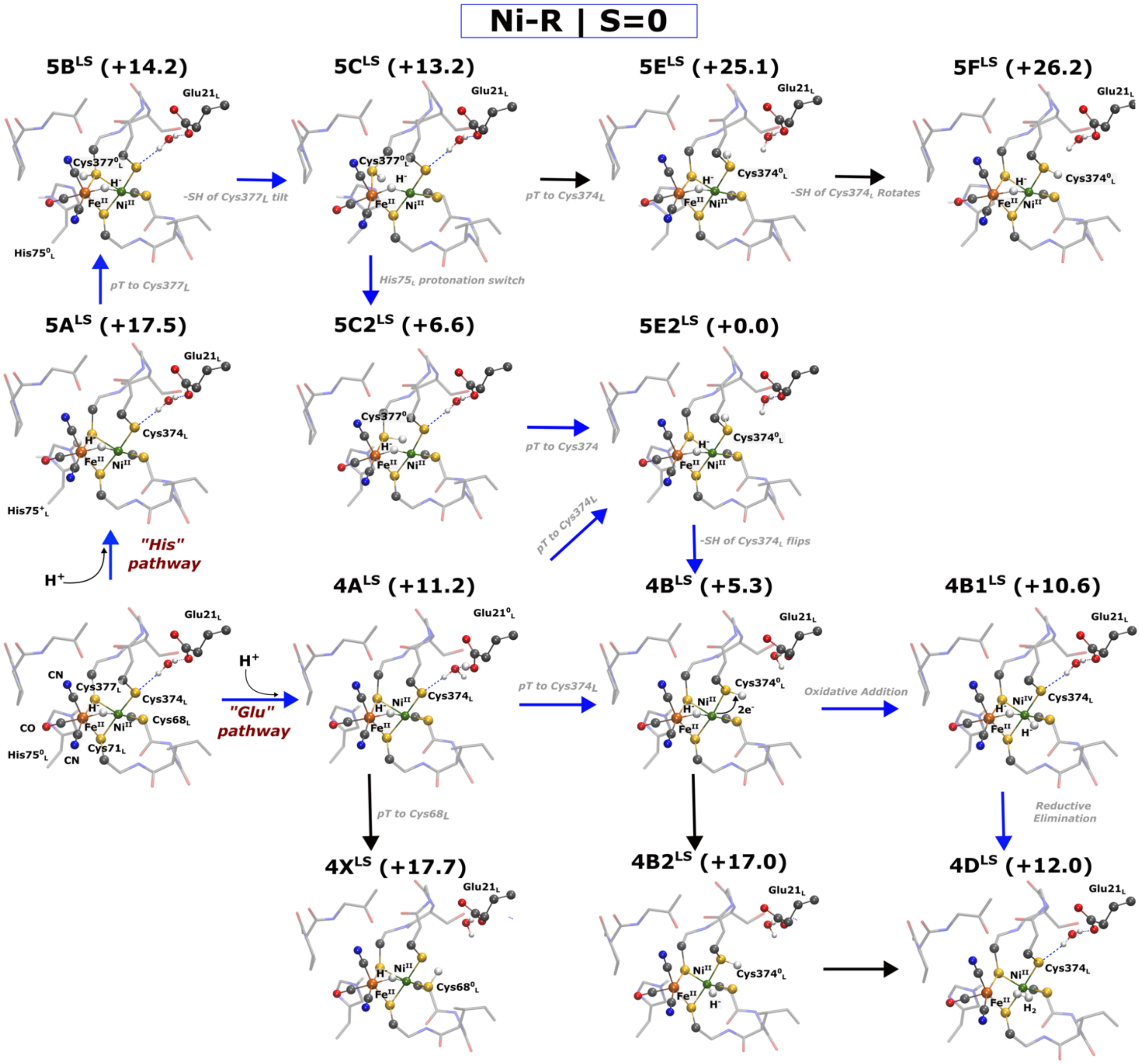


**Figure S12. DFT models of the Ni-R state with low-spin Ni(II).** Core structure of the DFT optimized models of [NiFe] active-site modelled in the Ni-R state ($Ni^{II}Fe^{II}$). The $Ni^{II}$ is modeled in low-spin configuration with S=0. Relative electronic energy (in kcal $mol^{-1}$) computed at B3LYP*/x2c-TZVPP/x2c-TZVP level of theory is also depicted. Blue and black arrows depict exergonic and endergonic reaction progression. Only the core structure is depicted along with key residues. Key bond distances for all models are shown in Table S11.

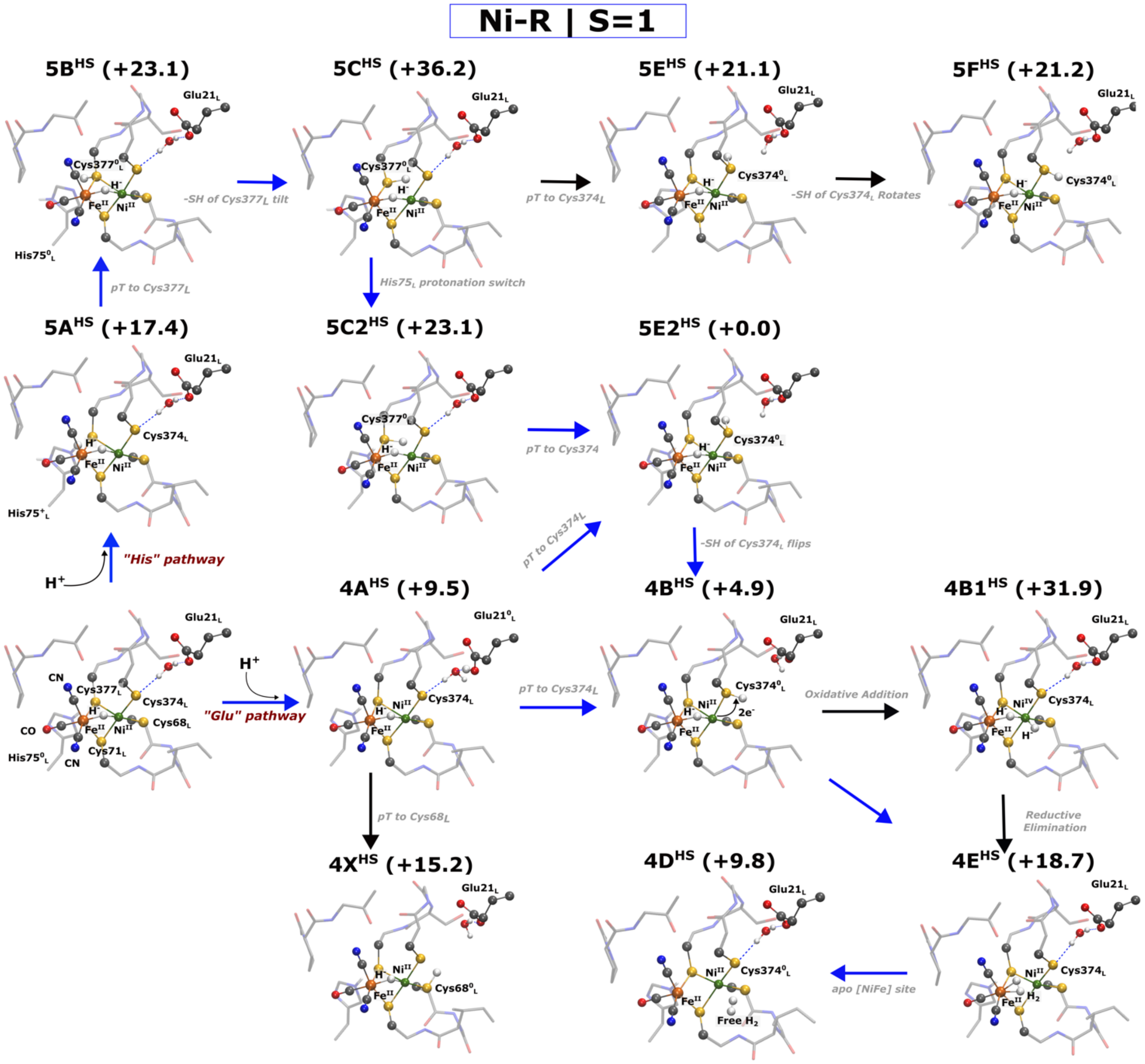


**Figure S13. DFT models of the Ni-R state with high-spin Ni(II).** Core structure of the DFT optimized models of [NiFe] active-site modelled in the Ni-R state ($Ni^{II}Fe^{II}$). The $Ni^{II}$ is modeled in high-spin configuration with S=1. Relative electronic energy (in kcal $mol^{-1}$) computed at B3LYP*/x2c-TZVPP/x2c-TZVP level of theory is also depicted. Blue and black arrows depict exergonic and endergonic reaction progression. Only the core structure is depicted along with key residues. Key bond distances and mulliken spin-populations for all models are shown in Table S11 and Table S12, respectively.

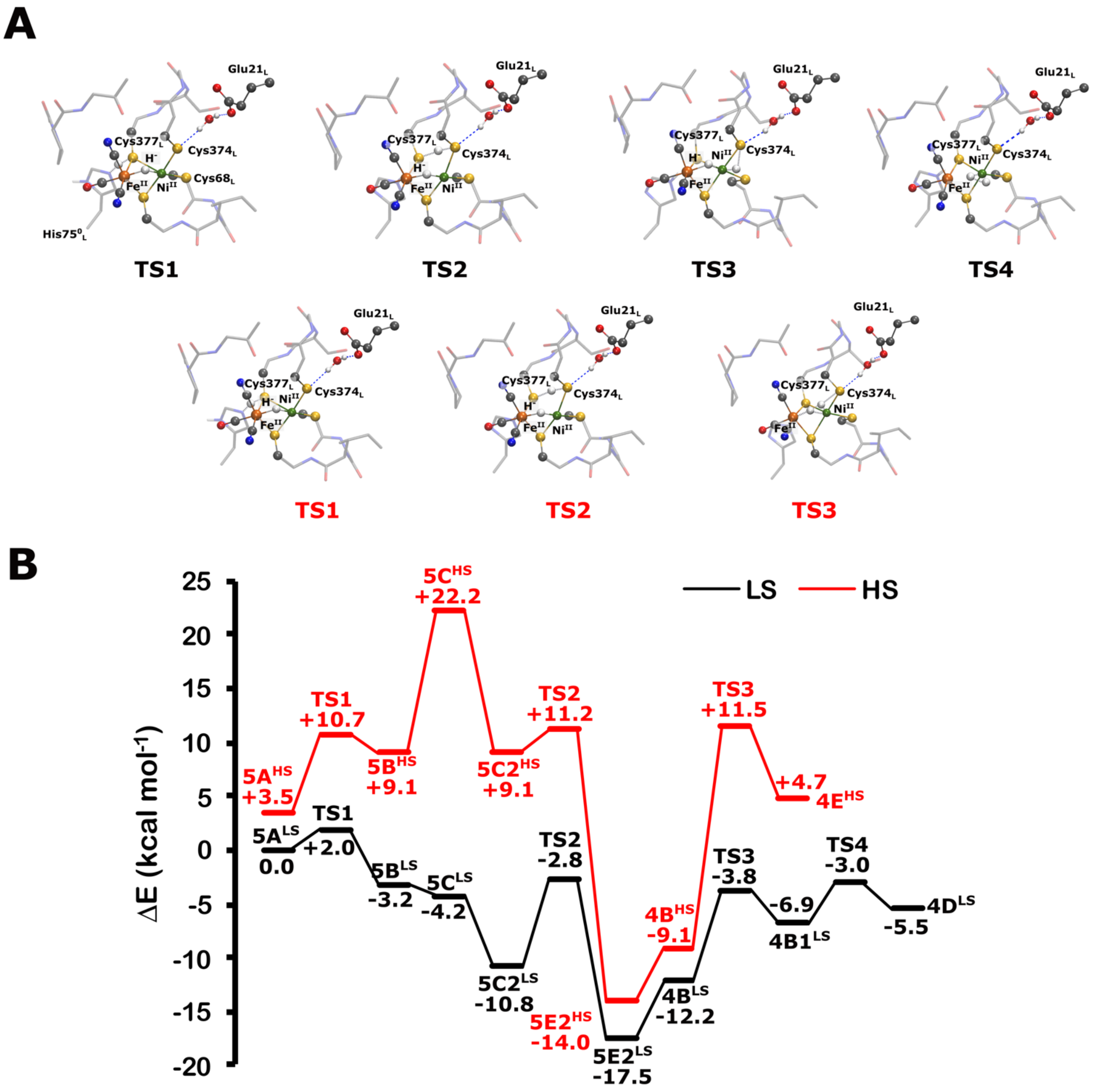


**Figure S14. Reaction energy profile for proton transfer via the His pathway during $H_2$ formation in the LS and HS forms.** The structures were optimized in both LS and HS forms. Single-point energy computations were performed with B3LYP*/x2c-TZVPPall/x2c-TZVPall level of theory. The corresponding transition state structures for both LS and HS configurations are also depicted. Intermediate structures are shown Figures S12 and S13.

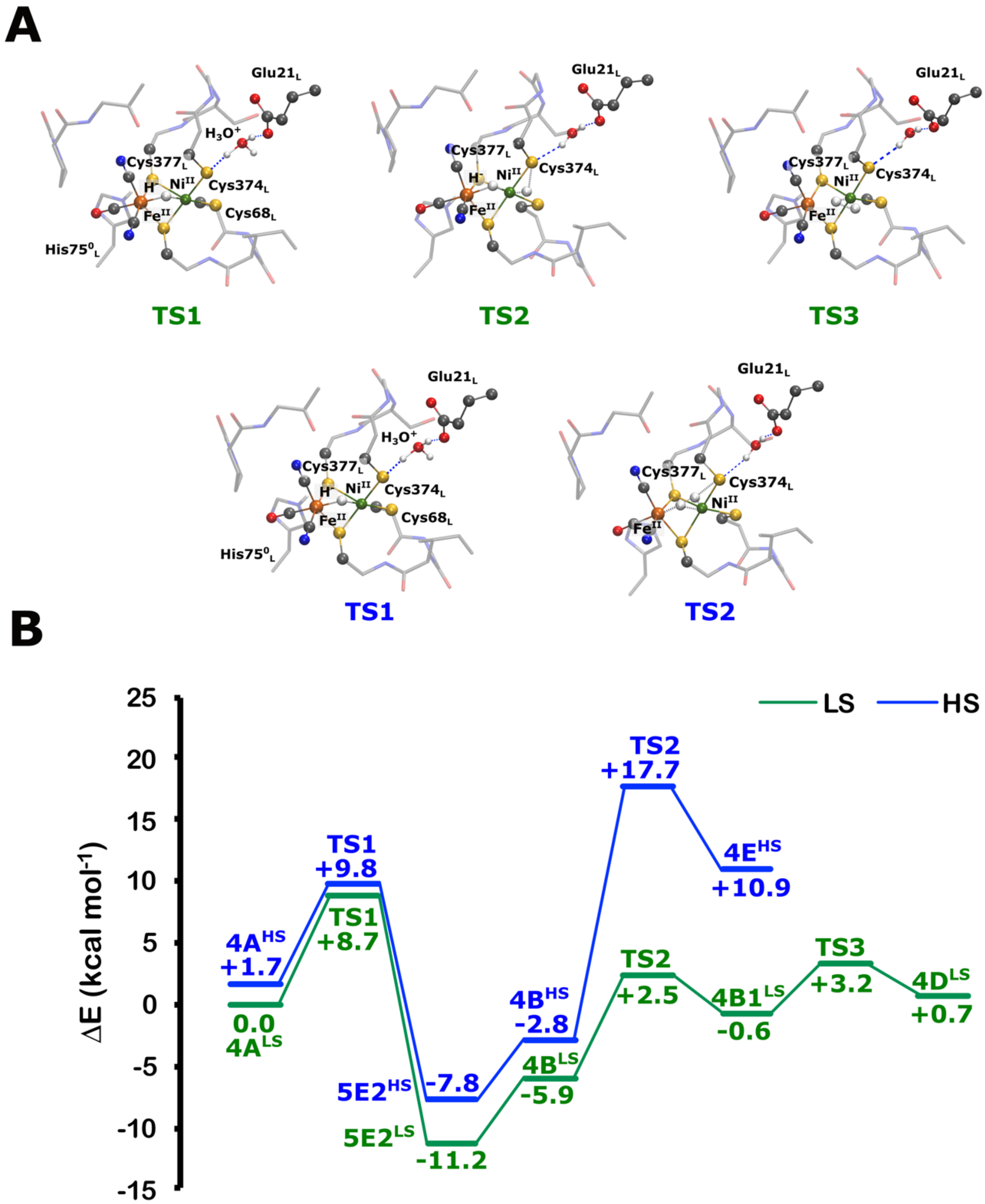


**Figure S15. Reaction energy profile for proton transfer via the Glu pathway during $H_2$ formation in the LS and HS forms.** The structures were optimized in both LS and HS forms. Single-point energy computations were performed with B3LYP*/x2c-TZVPPall/x2c-TZVPall level of theory. The corresponding transition state structures for both LS and HS configurations are also depicted. Intermediate structure is shown Figures S12 and S13.

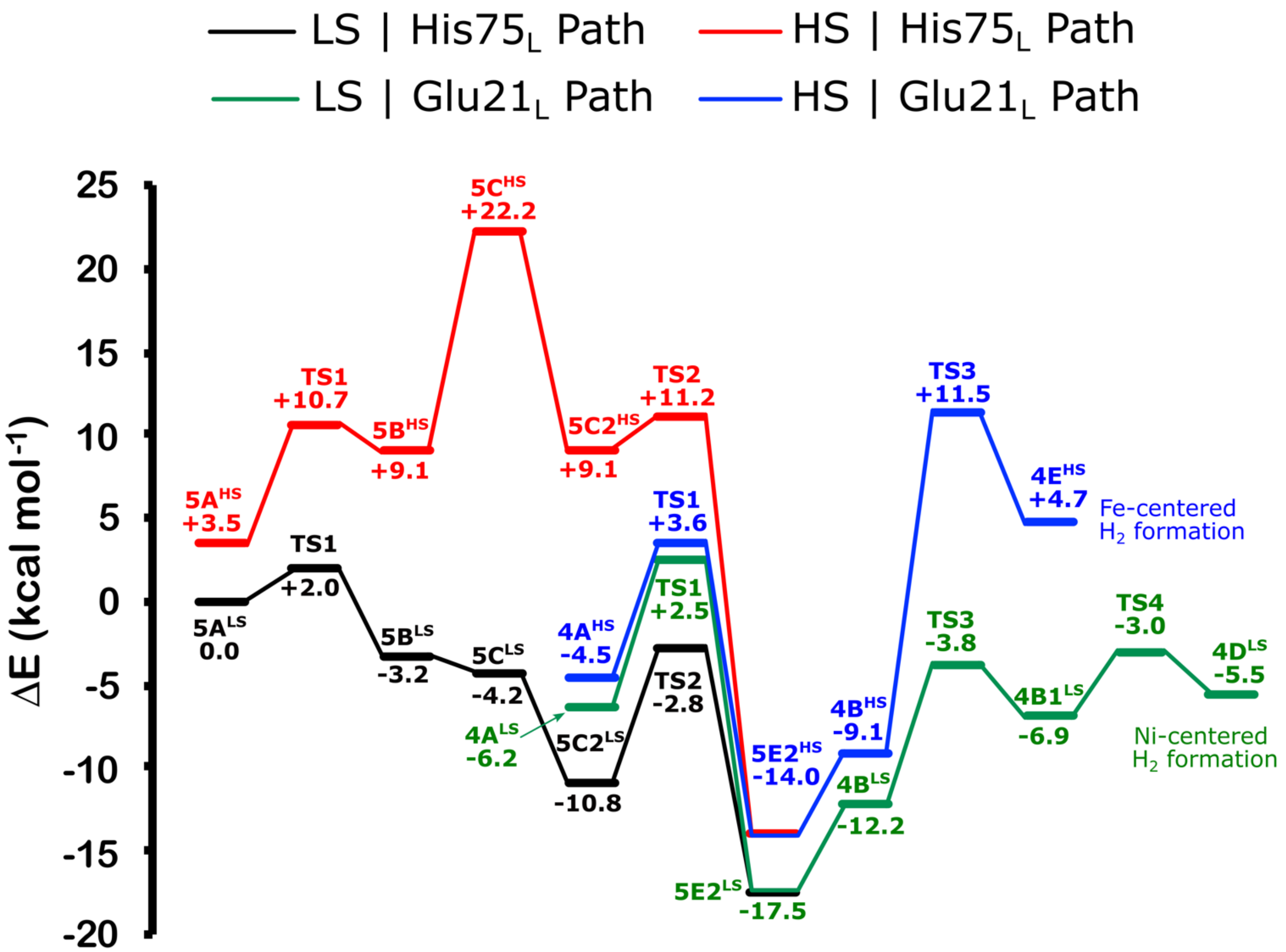


**Figure S16. Combined reaction energy profile for proton transfer along the His and Glu pathways in the LS and HS forms.** The structures were optimized in both LS and HS forms. Single-point energy computations were performed with B3LYP*/x2c-TZVPPall/x2c-TZVPall level of theory. Corresponding TS and intermediate structures were shown in Figures S14-S15 and S12-S13, respectively.

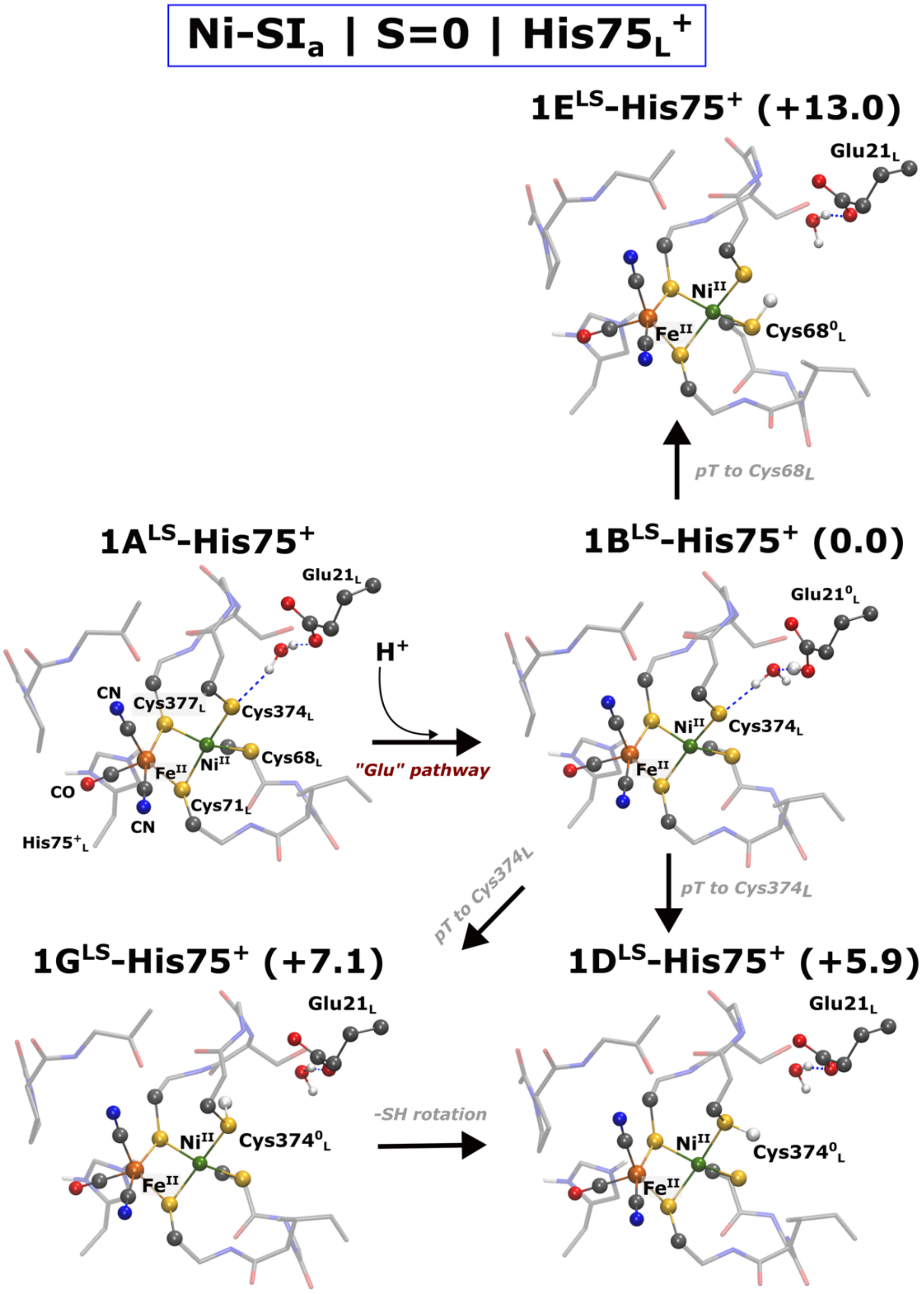


**Figure S17. DFT models of the Ni-SI$_a$ state with low-spin Ni(II) in the presence of His75$_L^+$.** Core structure of the DFT optimized models of [NiFe] active-site modelled in the Ni-SI$_a$ state (Ni$^{II}$Fe$^{II}$). The Ni$^{II}$ is modeled in low-spin configuration with S=0. Relative electronic energy (in kcal mol$^{-1}$) computed at B3LYP*/x2c-TZVPP/x2c-TZVP level of theory is also depicted. Only the core structure is depicted along with key residues. Key bond distances for all models are shown in Table S15.

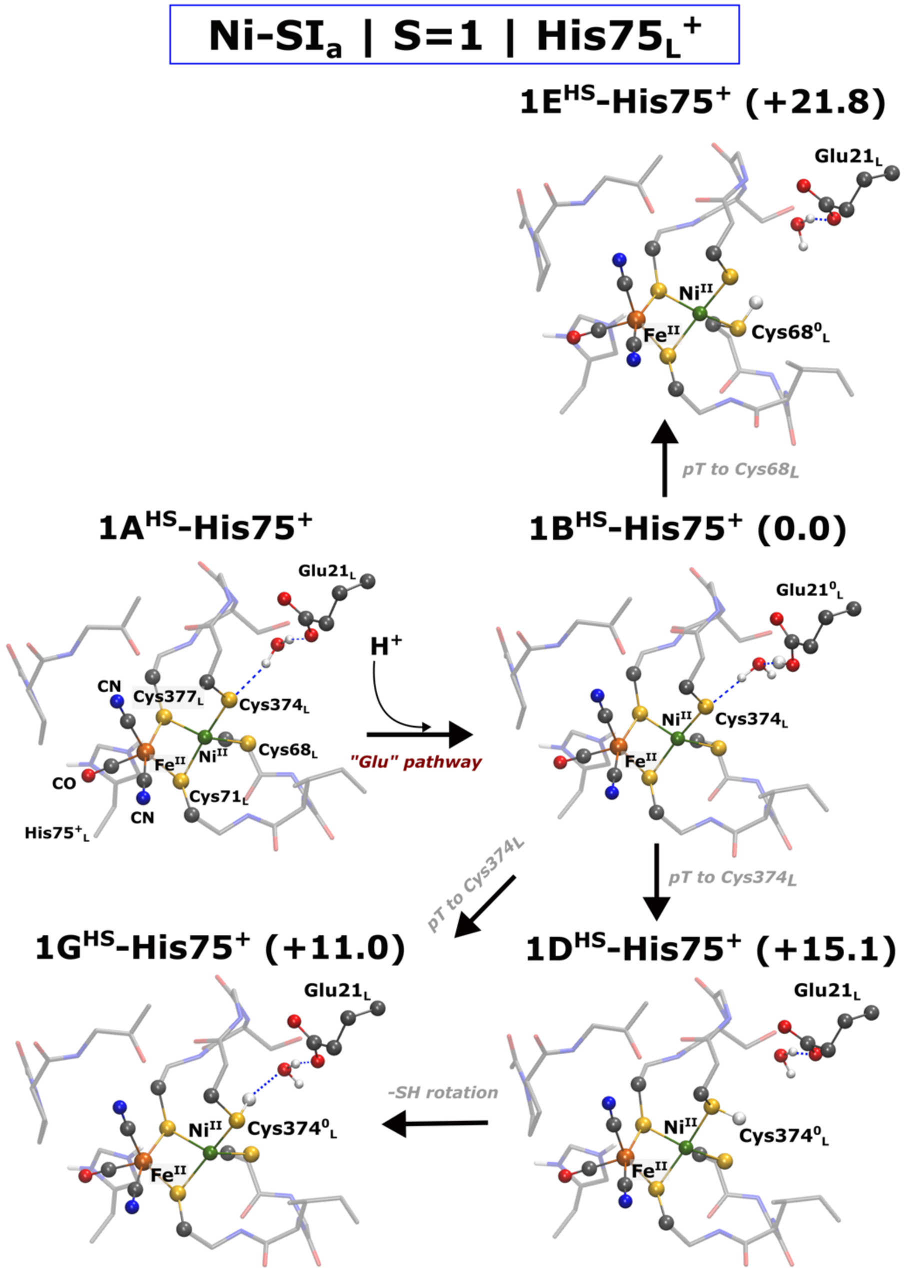


**Figure S18. DFT models of the Ni-SI$_a$ state with high-spin Ni(II) in the presence of His75$_L^+$.** Core structure of the DFT optimized models of [NiFe] active-site modelled in the Ni-SI$_a$ state ($Ni^{II}Fe^{II}$). The $Ni^{II}$ is modeled in high-spin configuration with S=1. Relative electronic energy (in kcal mol$^{-1}$) computed at B3LYP*/x2c-TZVPP/x2c-TZVP level of theory is also depicted. Only the core structure is depicted along with key residues. Key bond distances and Mulliken spin-populations for all models are shown in Table S15 and Table S16, respectively.

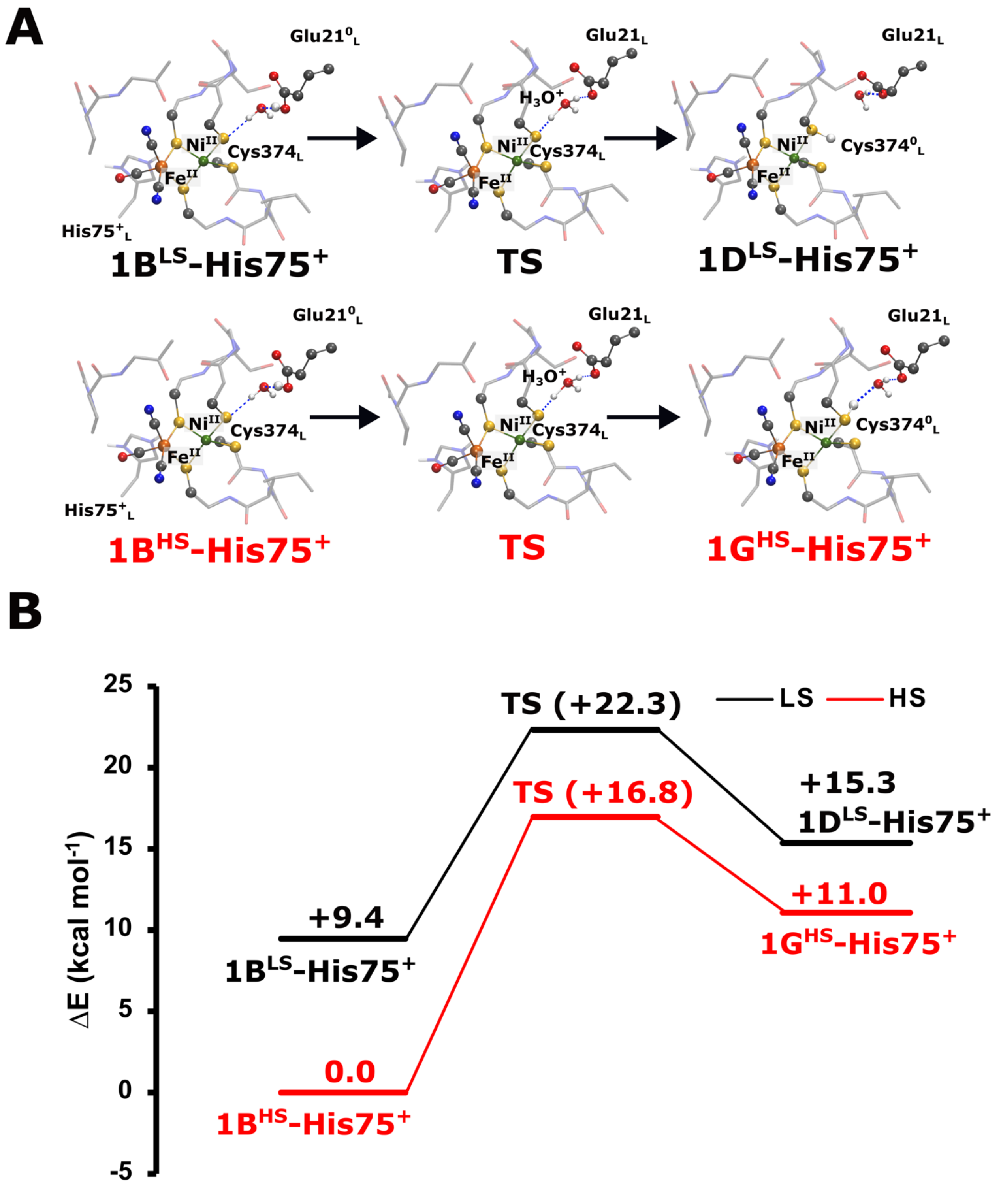


**Figure S19. Reaction energy profiles for water-mediated proton transfer from $Glu21_L$ to $Cys374_L$ in the Ni-$SI_a$ state in the presence of $His75_L^+$.** The complete reaction pathway was investigated in both low-spin (LS, black line) and high-spin configurations (HS, red line). The structures were optimized in both LS and HS forms. Single-point energy computations were performed with B3LYP*/x2c-TZVPPall/x2c-TZVPall level of theory. The reactant, product and transition state structures corresponding to both spin-state are depicted.

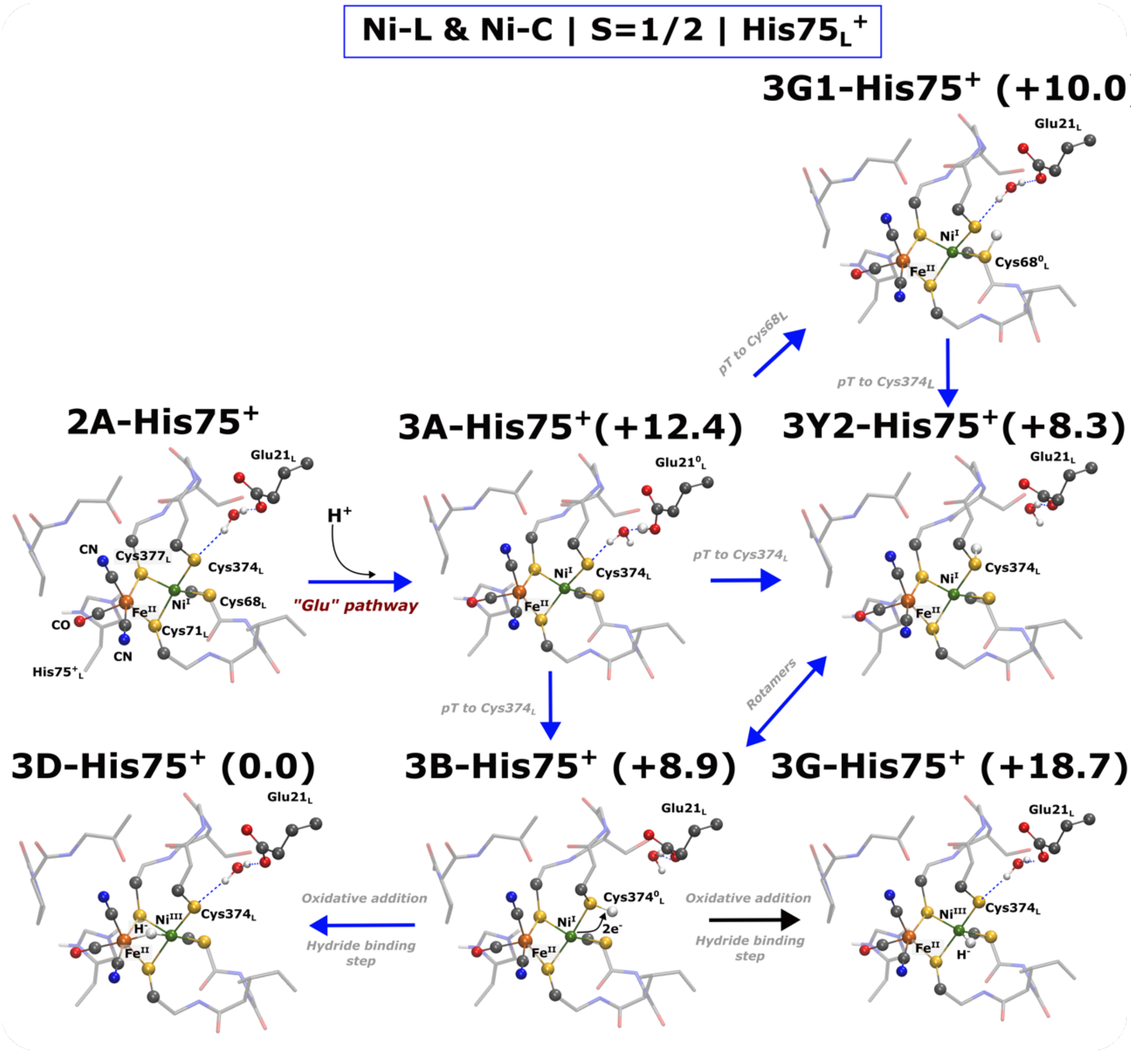


**Figure S20. DFT models of the Ni-L and Ni-C states in the presence of His75$_L^+$.** Core structure of the DFT optimized models of [NiFe] active-site modelled in the Ni-L ($Ni^IFe^{II}$) and Ni-C state ($Ni^{III}Fe^{II}$). In both states, $Ni^{I/III}$ has S=1/2. Relative electronic energy (in kcal $mol^{-1}$) computed at B3LYP*/x2c-TZVPP/x2c-TZVP level of theory is also depicted. Blue and black arrows depict exergonic and endergonic reaction progression. Only the core structure is depicted along with key residues. Key bond distances for all models and Mulliken spin-populations are shown in Table S17 and Table S18, respectively.

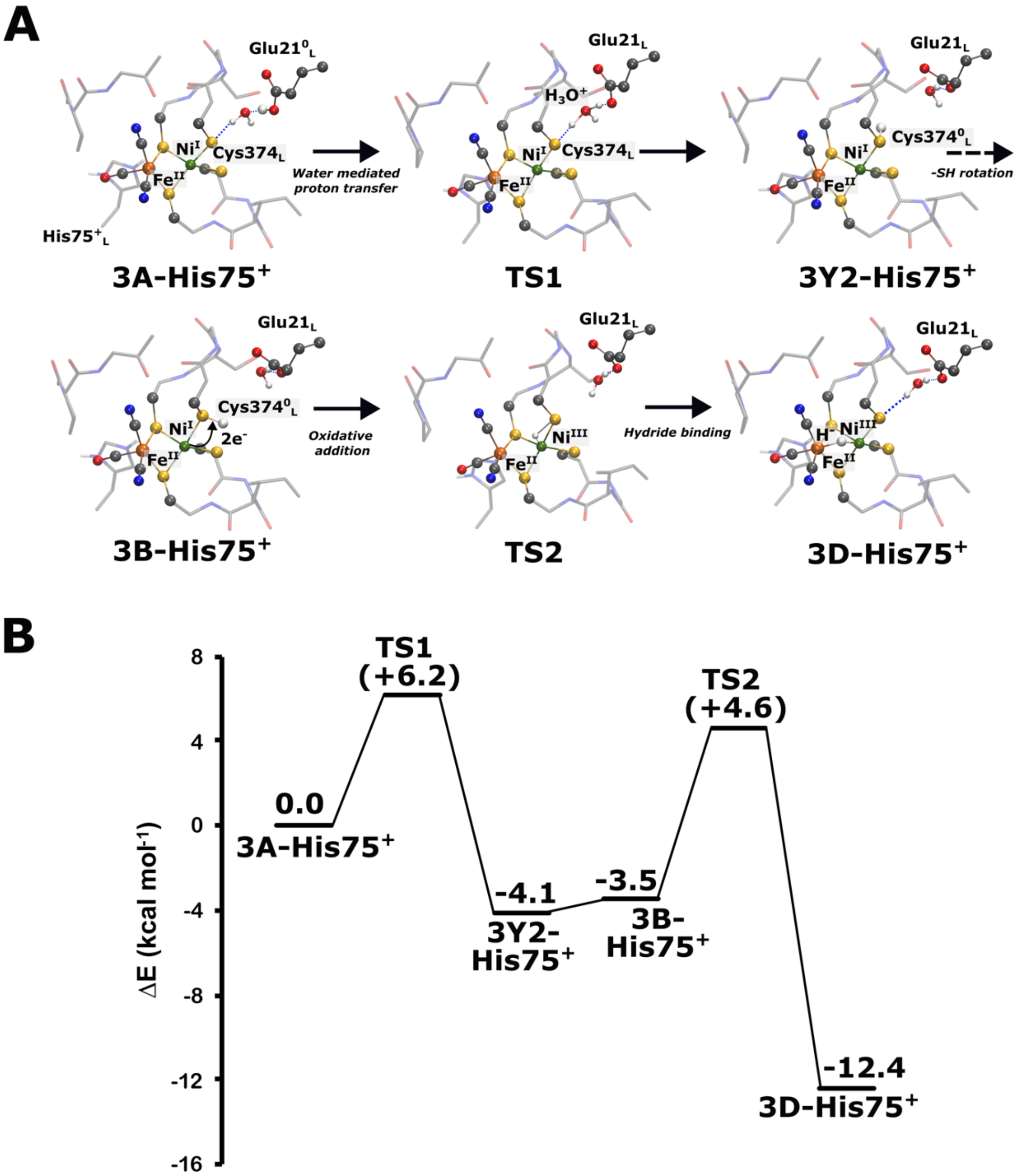


**Figure S21. Reaction energy profiles for water-mediated proton transfer from $Glu21_L$ to $Cys374_L$ and hydride binding step involving protonated $Cys374_L$ and $Ni^{III}$ in the presence of $His75_L^+$.** Single-point energy computations were performed with B3LYP*/x2c-TZVPPall/x2c-TZVPall level of theory. The reactant, product and transition state structures are also depicted.

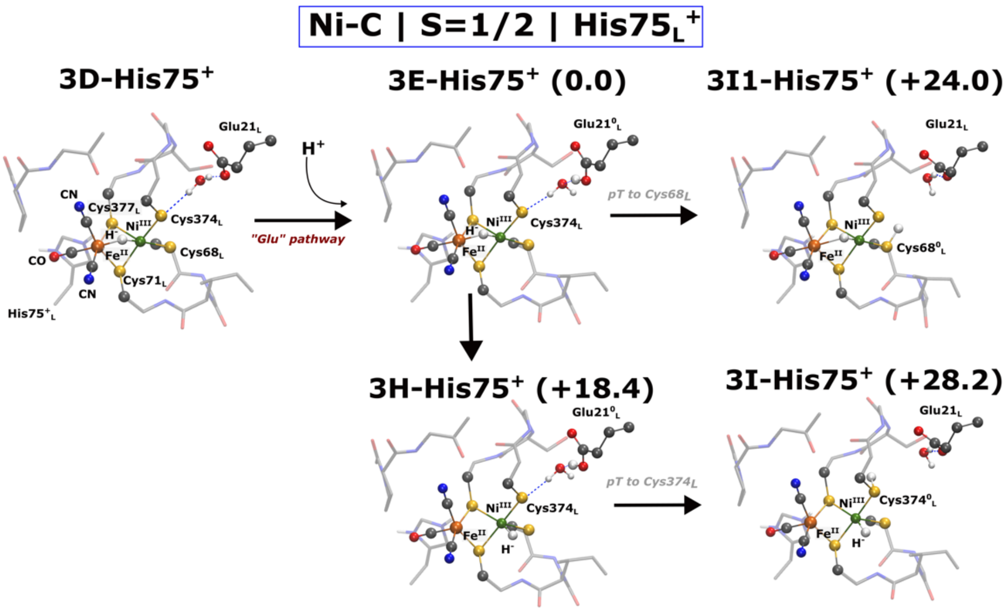


**Figure S22. DFT models for the Ni-C catalytic state for the second proton transfer step in presence of His75$_L^+$.** Core structure of the DFT optimized models of [NiFe] active-site modelled Ni-C state (Ni$^{III}$Fe$^{II}$, S=1/2). Relative electronic energy (in kcal mol$^{-1}$) computed at B3LYP*/x2c-TZVPP/x2c-TZVP level of theory is also depicted. Only the core structure is depicted along with key residues. Key bond distances for all models and Mulliken spin-populations are shown in Table S17 and Table S18, respectively.

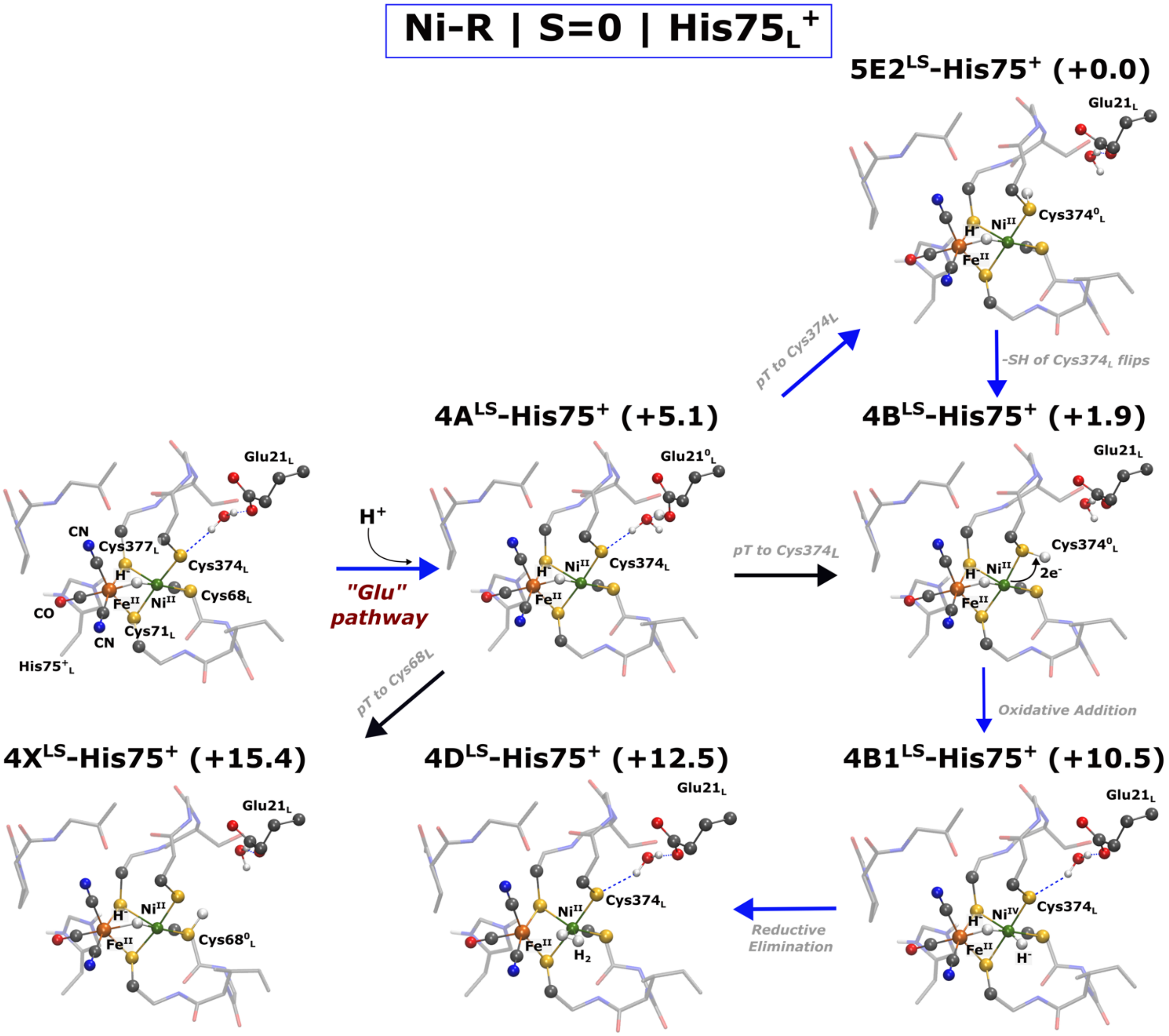


**Figure S23. DFT models of the Ni-R state with low-spin Ni(II) in presence of His75$_L^+$.** Core structure of the DFT optimized models of [NiFe] active-site modelled in the Ni-R state (Ni$^{II}$Fe$^{II}$). The Ni$^{II}$ is modeled in low-spin configuration with S=0. Relative electronic energy (in kcal mol$^{-1}$) computed at B3LYP*/x2c-TZVPP/x2c-TZVP level of theory is also depicted. Blue and black arrows depict exergonic and endergonic reaction progression. Only the core structure is depicted along with key residues. Key bond distances for all models are shown in Table S20.

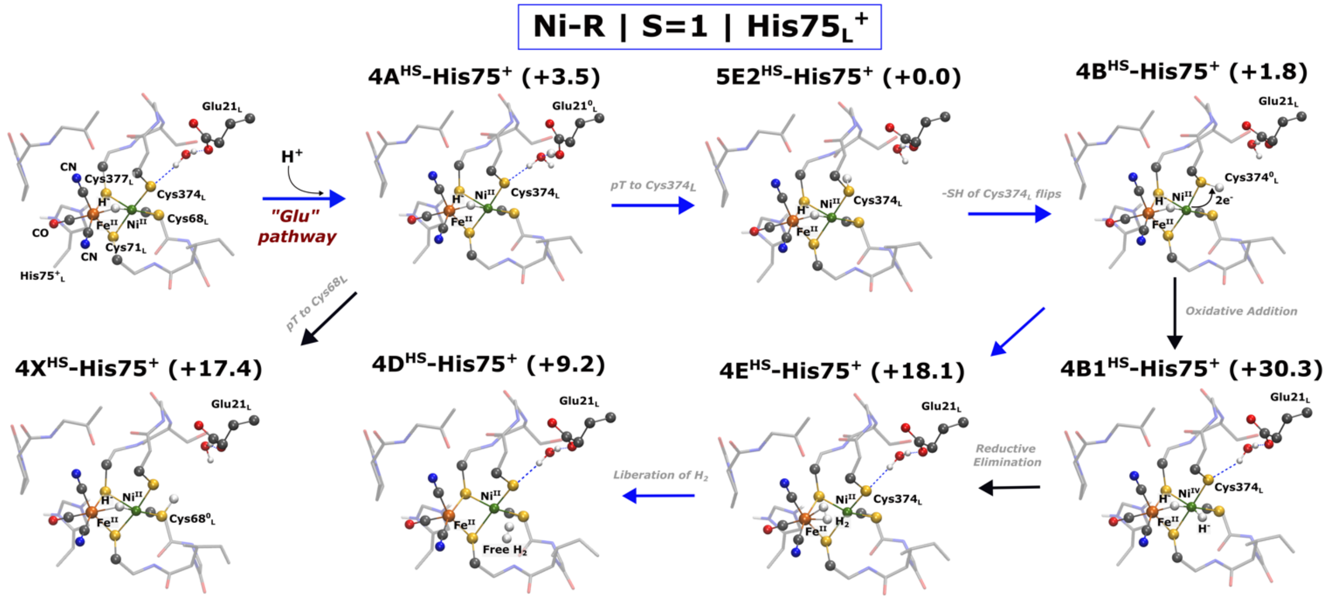


**Figure S24. DFT models of the Ni-R state with high-spin Ni(II) in presence of $His75_L^+$.** Core structure of the DFT optimized models of [NiFe] active-site modelled in the Ni-R state ($Ni^{II}Fe^{II}$). The $Ni^{II}$ is modeled in high-spin configuration with S=1. Relative electronic energy (in kcal $mol^{-1}$) computed at B3LYP*/x2c-TZVPP/x2c-TZVP level of theory is also depicted. Blue and black arrows depict exergonic and endergonic reaction progression. Only the core structure is depicted along with key residues. Key bond distances and Mulliken spin-populations for all models are shown in Table S20 and Table S21.

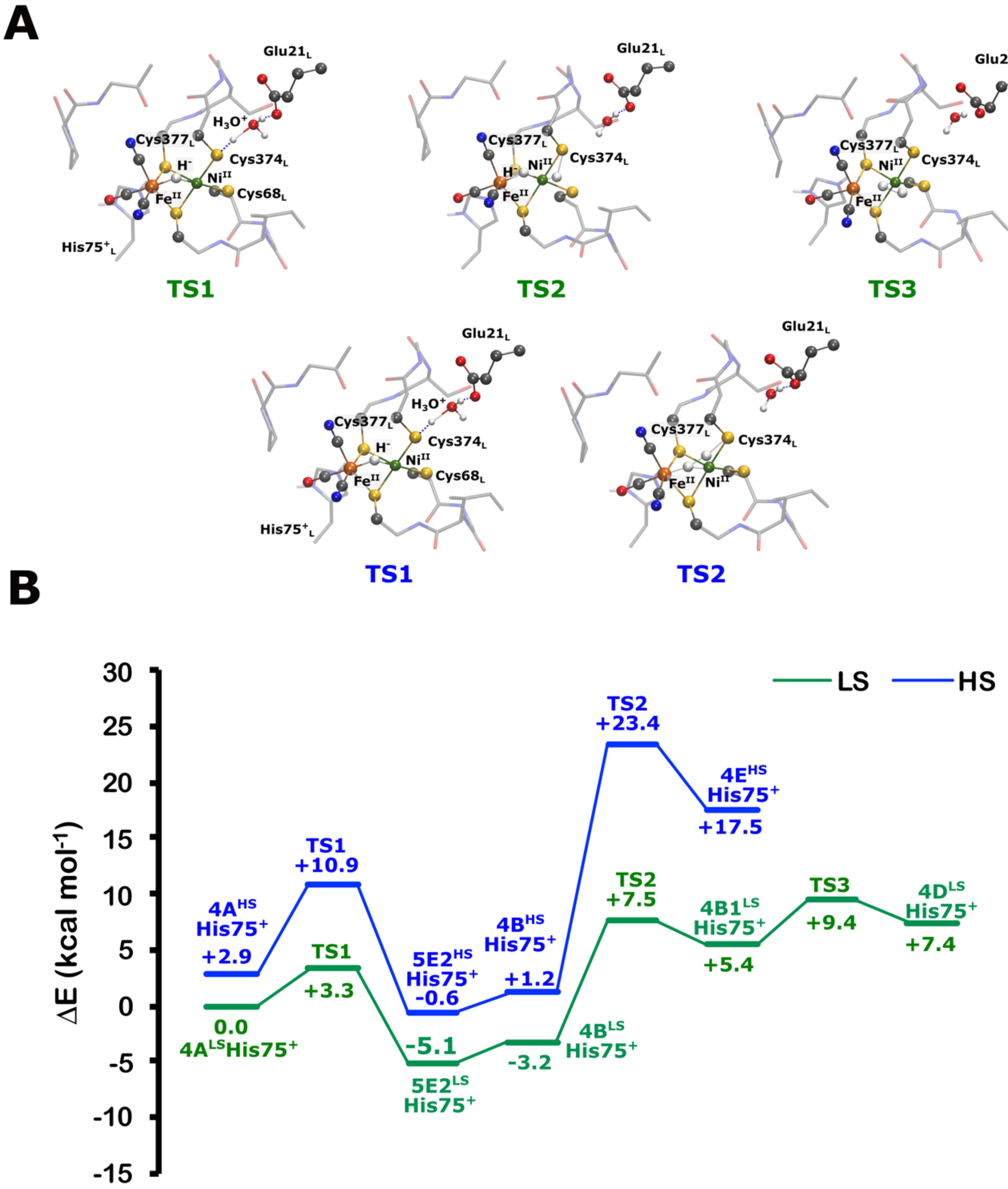


**Figure S25. Reaction energy profile for the proton originating from the "Glu" pathway for the LS and HS forms in the presence of His75$_L$$^+$.** The structures were optimized in both LS and HS forms. Single-point energy computations were performed with B3LYP*/x2c-TZVPPall/x2c-TZVPall level of theory. The corresponding transition state structures are also depicted. Intermediate structure is shown Figures S23 and S24.

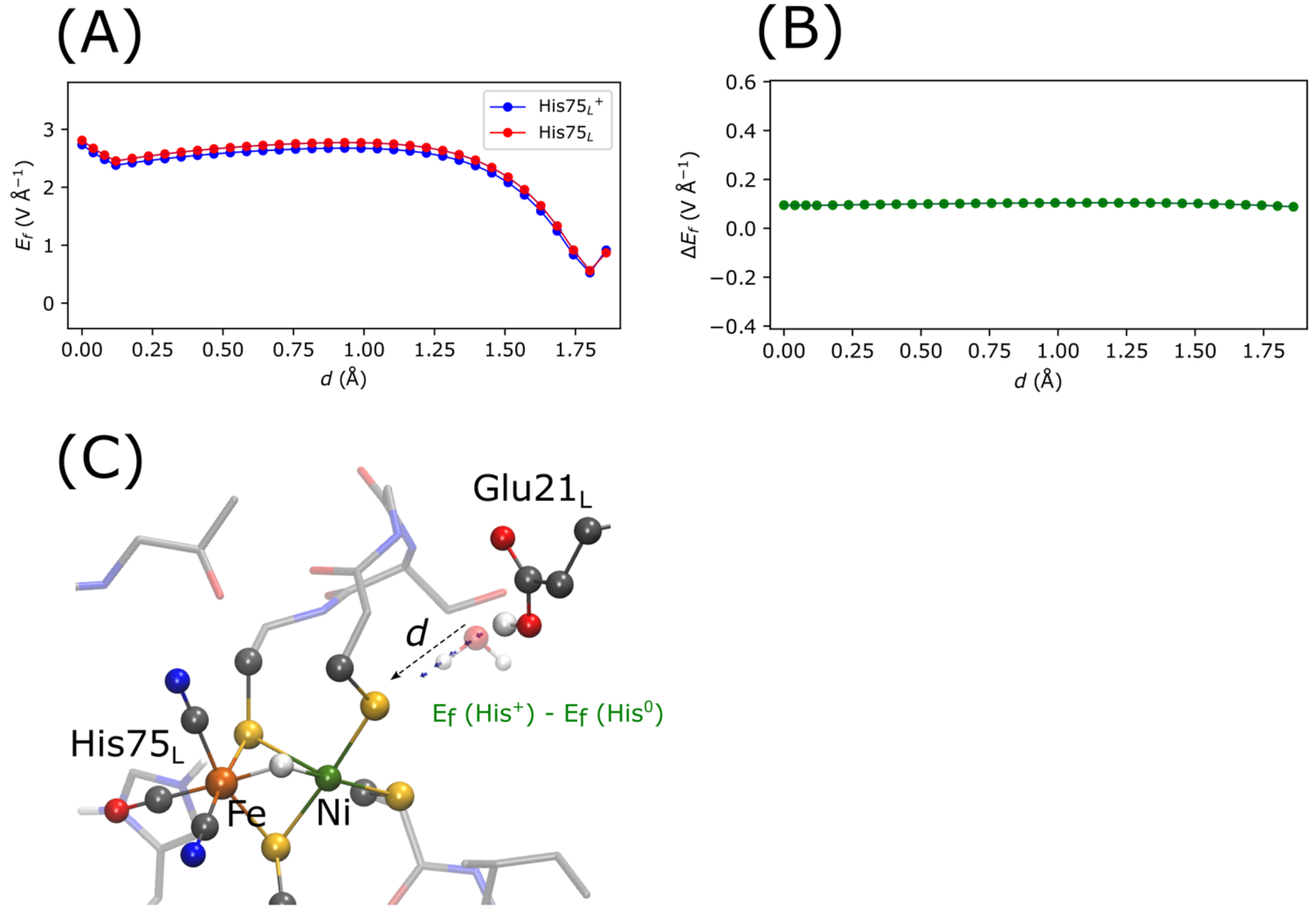


**Figure S26. Electric field effects on the proton transfer in the Ni-R state and their influence on the protonation state of His75$_L$.** (A) Absolute electric field strength along the $H_2O$ - Cys374$_L$ proton transfer coordinate in the Ni-R with neutral (**4A$^{LS}$**) and protonated His75$_L$ (**4A$^{LS}$ His75$^+$**). (B) Electric field difference along the proton transfer coordinate in the models with (**4A$^{LS}$ His75$^+$**) and without protonated (**4A$^{LS}$**) His75$_L$ in the Ni-R state. (C) Depiction of difference electric field vectors and proton transfer coordinate (*d*).

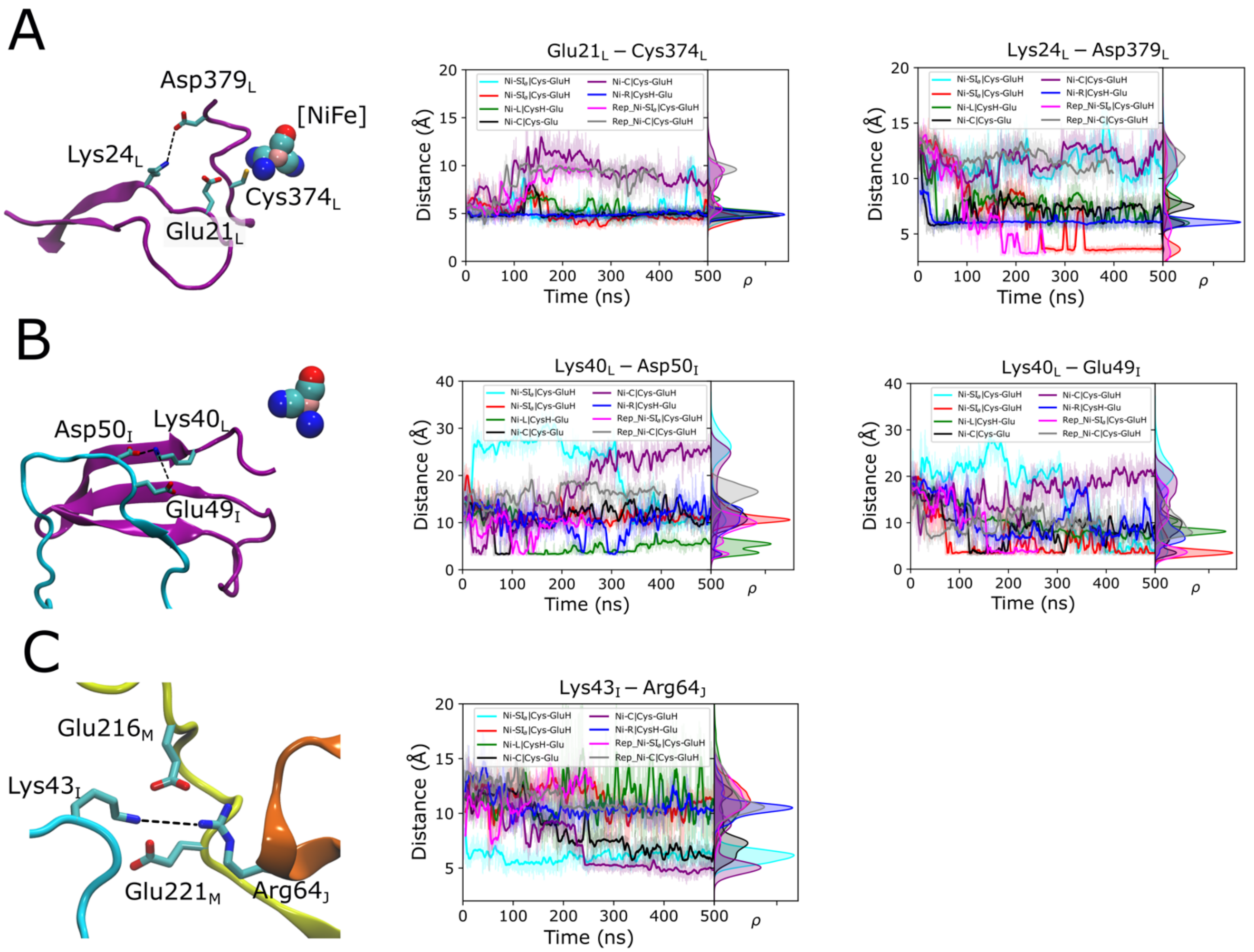


**Figure S27. Conformational dynamics of key residues involved in the loop network.** (A) Dynamics of Glu21$_L$-Cys374$_L$ and Lys24$_L$-Asp379$_L$ ion-pairs in the β1- β2 loop of the MbhL subunit for all catalytic states of the [NiFe] cluster. (B) ion-pair network (Lys40$_L$-Asp50$_I$ and Lys40$_L$-Glu49$_I$) between β1- β2 loop of the MbhL subunit and TM1-2 loop of MbhI subunit. (C) Interaction of MbhI, MbhM and MbhJ loops region via interaction between Lys43$_I$, Arg64$_J$, Glu216$_M$ and Glu221$_M$. Dataset 'Ni-SI$_a$|Cys-GluH' (Cyan) corresponds to the Sim1A, 'Ni-SI$_a$|Cys-GluH' (red) is Sim2A, 'Ni-L|CysH-Glu' (Green) is Sim3A, 'Ni-C|Cys-Glu' (black) is Sim4A, 'Ni-C|Cys-GluH' (maroon) is Sim5A, 'Ni-R|CysH-Glu' (Blue) is Sim6A. Dataset 'Rep_Ni-SI$_a$|Cys-GluH' and 'Rep_Ni-C|Cys-GluH' are replica simulation for the Sim2A and Sim5A, respectively. The core structure of the [NiFe] in the Ni-SI$_a$, Ni-L, Ni-C and Ni-R is same as **1B$^{LS}$**, **3B**, **3D** and **4B$^{LS}$**, respectively. Details on the classical MD set-up is shown in Figure S22-S23.

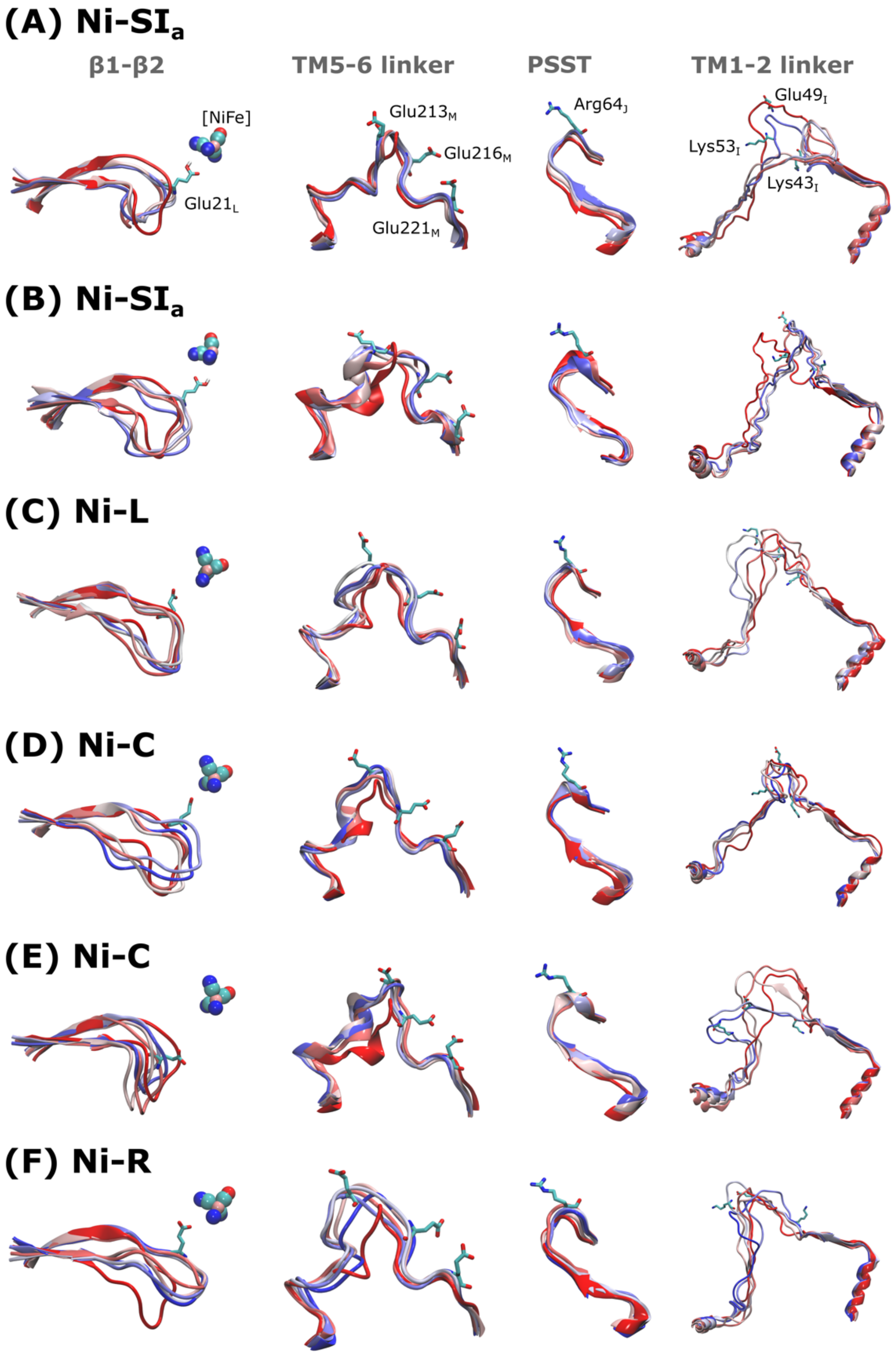


**Figure S28. Conformational sampling of loops involved in the network.** Depiction of spatial clustering of dynamics β1- β2 loop (MbhL subunit), TM5-6 linker loop (MbhM), PSST loop (MbhJ) and TM1-2 linker loop (MbhI) for all [NiFe] catalytic states. Key residues involved in the structural coupling are also showcased. Details on the classical MD set-up is shown in Figure S22-S23.

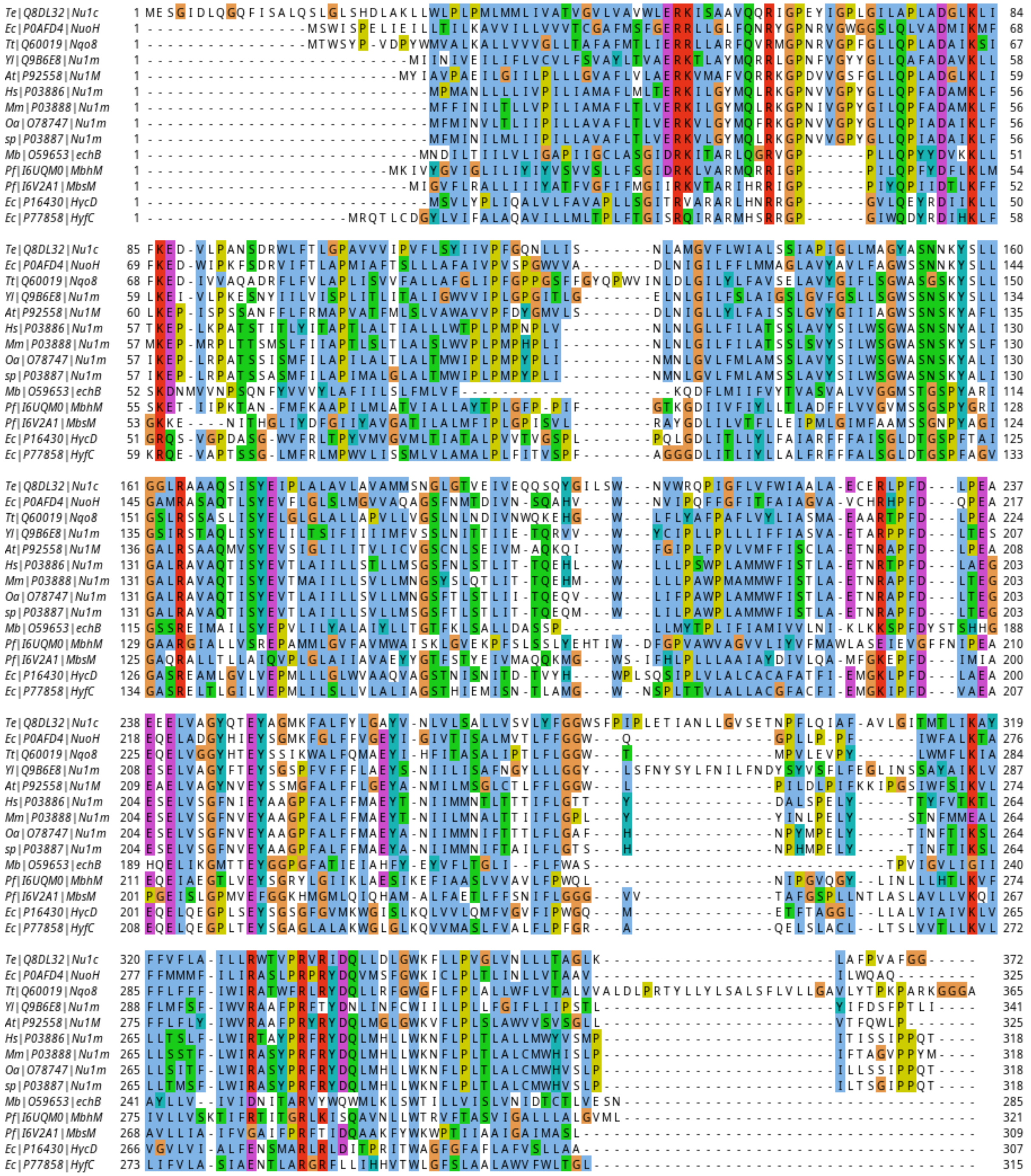

**Figure S29. Multiple sequence alignment of MbhM.** Multiple sequence alignment for MbhM *Pyrococcus furiosus* subunit aligned with proteins from the complex I superfamily: *Ovis aries* (O78747|NU1M), *Bos taurus* (P03887|NU1M), *Mus musculus* (P03888|NU1M), *Homo sapiens* (P03886|NU1M), *Yarrowia lipolytica* (Q9B6E8|NU1M), *Arabidopsis thaliana* (P92558|NU1M), *Thermus thermophilus* (Q60019|NQO8), *E. coli* (P0AFD4|NUOH), *Thermosynechococcus elongatus* (Q8DL32|NU1C), *E. coli* (P16430|HYCD), *E. coli* (P77858|HYFC), *Pyrococcus Furiosus* Mbh (I6UQM0|MbhM), *Pyrococcus furious* Mbs (I6V2A1|MbhM), *Methanosarcina barkeri* (O59653|ECHB).

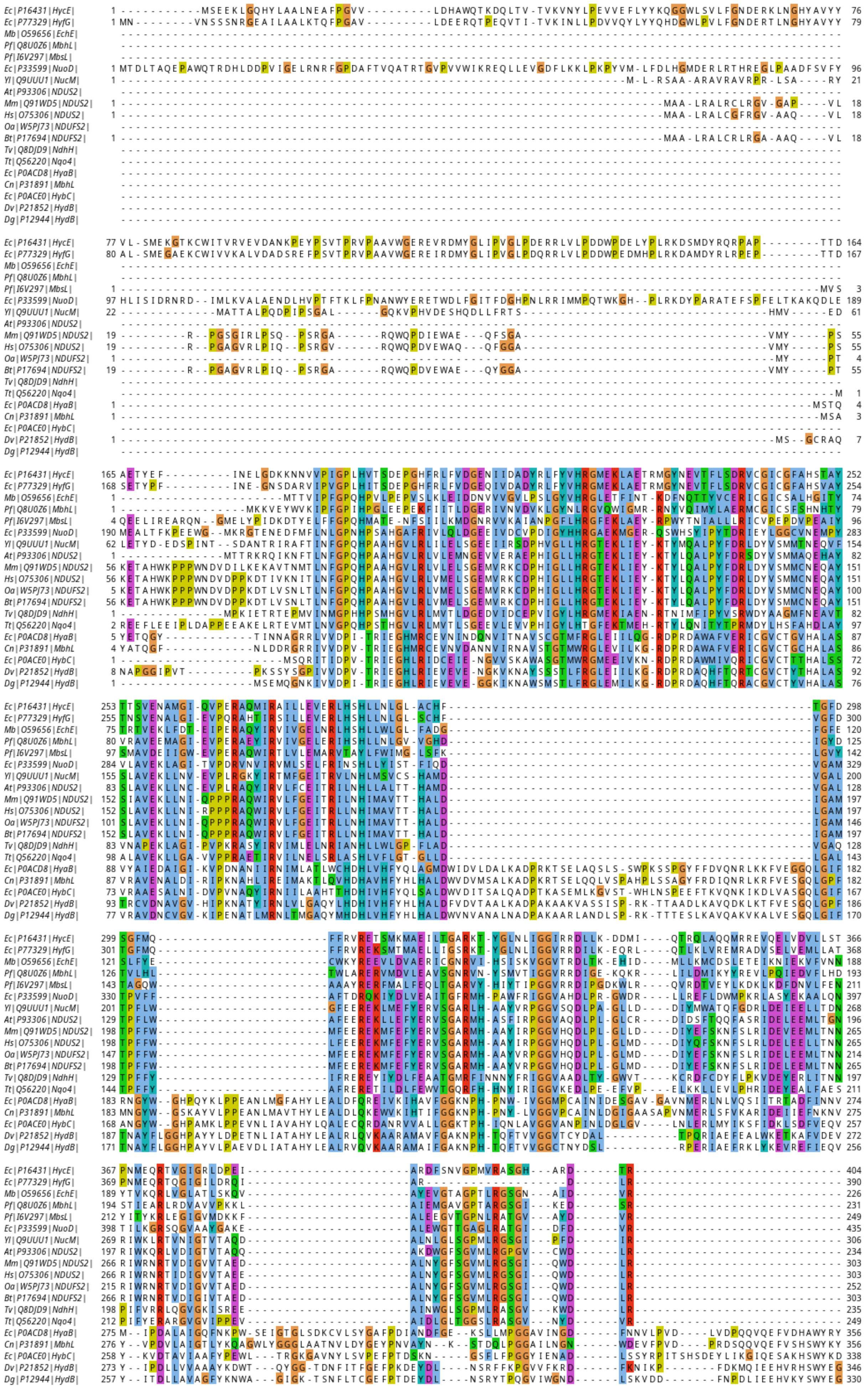
Ec|P16431|HycE|
Ec|P77329|HyfG|
Mb|O59656|EchE|
Pf|Q8U0Z6|MbhL|
Pf|I6V297|MbsL|
Ec|P33599|NuoD|
Yl|Q9UUU1|NucM|
At|P93306|NDUS2|
Mm|Q91WD5|NDUS2|
Hs|O75306|NDUS2|
Oa|W5PJ73|NDUFS2|
Bt|P17694|NDUFS2|
Tv|Q8DJD9|NdhH|
Tt|Q56220|Nqo4|
Ec|P0ACD8|HyaB|
Cn|P31891|MbhL
Ec|P0ACE0|HybC|
Dv|P21852|HydB|
Dg|P12944|HydB|

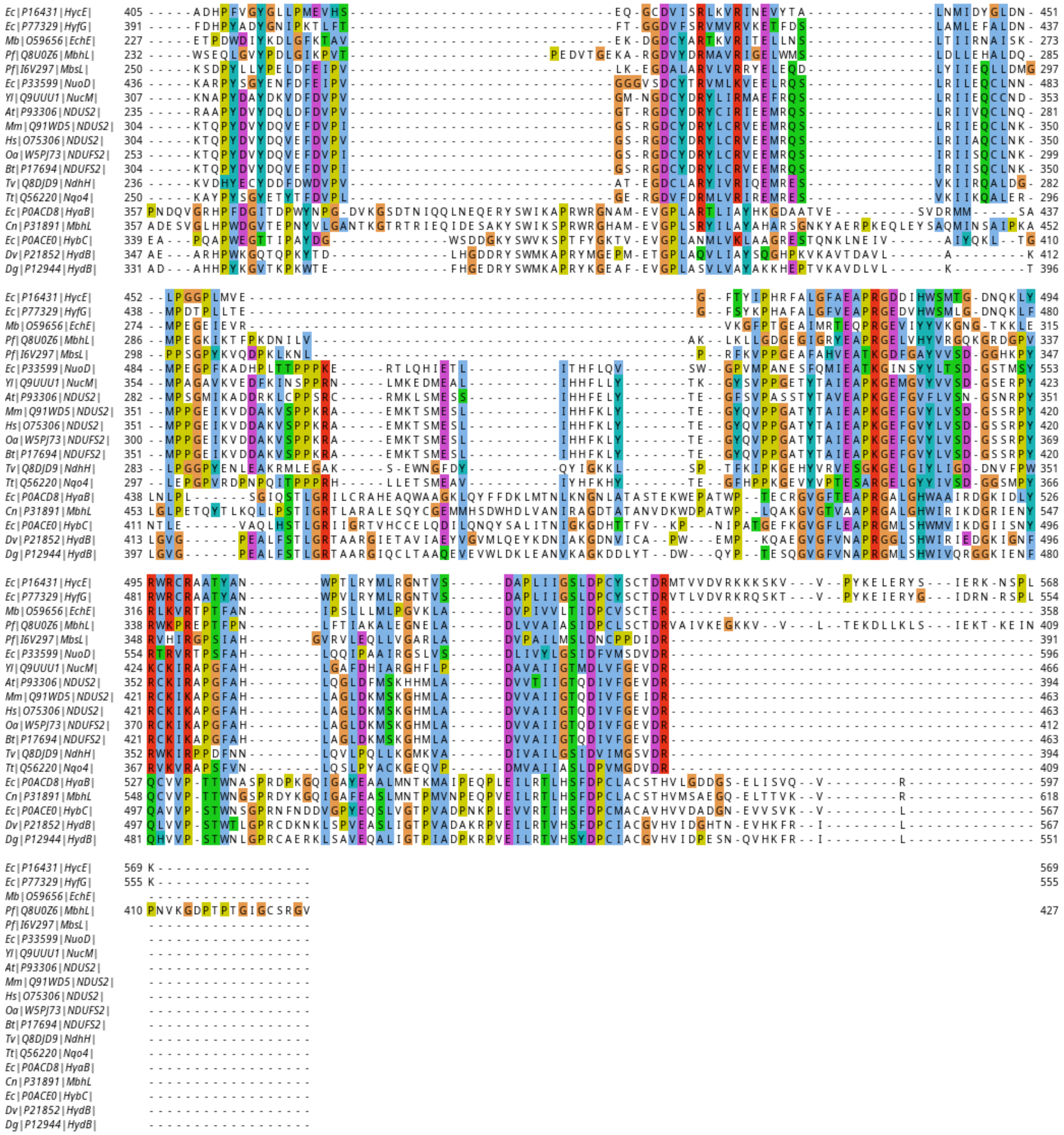

**Figure S30. Multiple sequence alignment of MbhL.** Multiple sequence alignment for MbhL *Pyrococcus furiosus* subunit aligned with proteins from the complex I superfamily and [Ni-Fe] hydrogenases: *Methanosarcina barkeri* (O59656|ECHE), *Pyrococcus furiosus* Mbh (Q8U0Z6|MbhL), *E. coli* Hyd-3 (P16431|HYCE), *E. coli* (P77329|HYFG), *Ovis Aries* (W5PJ73|NDUFS2), *Bos taurus* (P17694|NDUFS2), *Homo sapiens* (O75306|NDUFS2), *Mus musculus* (Q91WD5|NDUFS2), *Arabidopsis thaliana* (P93306|NDUFS2), *Yarrowia lipolytica* (Q9UUU1|NUCM), *Thermosynechococcus elongatus* (Q8DJD9|NDHH), *Thermus thermophillus* (Q56220|NQO4), *E. coli* (P33599|NUOCD_ECOLI), Pyrococcus furiosus Mbs (I6V297|MbhL), *Desulfovibrio vulgaris* (P21852|PHNL), *Desulfovibrio gigas* (P12944|PHNL), *E. coli* Hyd-2 (P0ACE0|MBHM), *E. coli* Hyd-1 (P0ACD8|MBHL), *Cupriavidus_necator* (P31891|MBHL), *Hydrogenpilus_thermoluteolus* (A0A077LAI5|HYDTE).

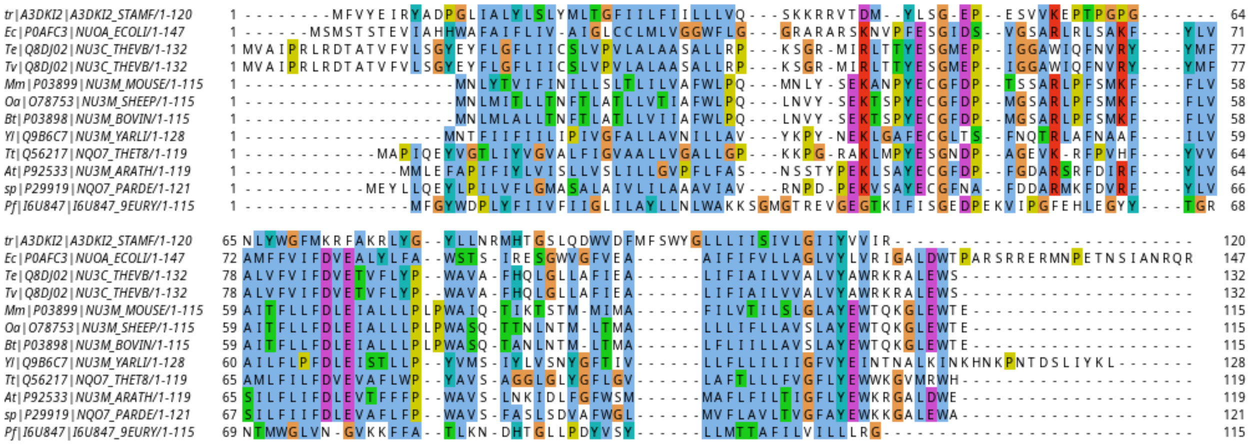


**Figure S31. Multiple sequence alignment of MbhI.** Multiple sequence alignment for MbhI *Pyrococcus furiosus* subunit aligned with proteins from the complex I superfamily: *Pyrococcus furiosus* Mbh (I6U847|MbhI), *E. coli* (P0AFC3|NuoA), *Thermosynechococcus elongatus* (Q8DJ02), *Thermosynecococcus vestitus* (Q8DJ02|Nu3C), Mus Musculus (P03899|Nu3M), Ovis aries (O78753|Nu3M), Bos taurus (P03898|Nu3M), *Yarrowia lipolytica* (Q986C7|Nu3M), *Thermus thermophilus* (Q56217|NQ07), *Arabidopsis thaliana* (P922533|NU3M), *Paracoccus denitrificans* (P29919|NQO7).

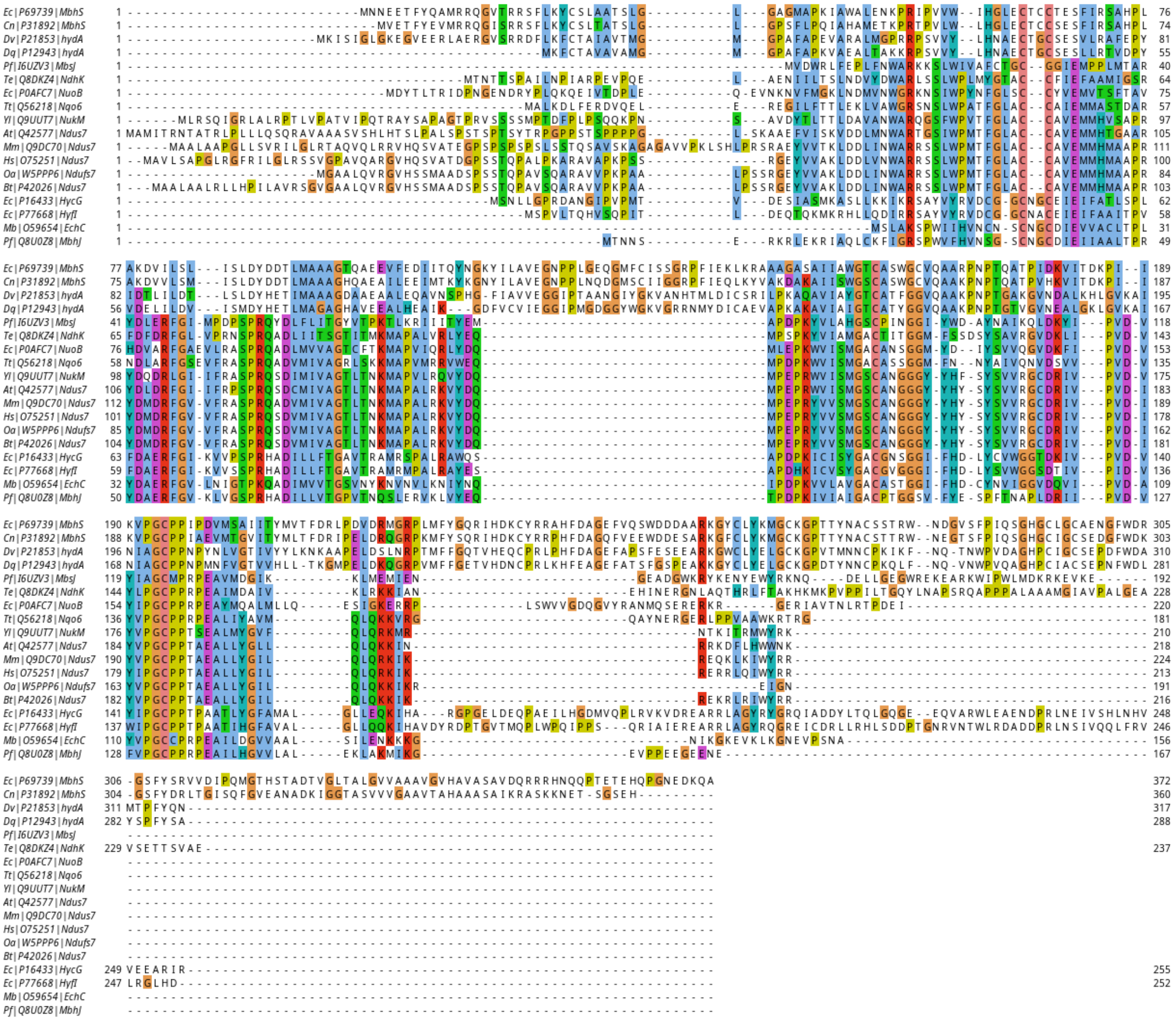

**Figure S32. Multiple sequence alignment of MbhJ.** Multiple sequence alignment for MbhJ *Pyrococcus furiosus* subunit aligned with proteins from the complex I superfamily and [Ni-Fe] hydrogenases: *E. coli* (P16433|HYCG), *E. coli* (P77668|HYFI), *Pyrococcus furiosus* Mbh (Q8U0Z8|MBHJ), *Methanosarcina barkeri* (O59654|ECHC), *Ovis aries* (W5PPP6|NDUS7), *Bos taurus* (P42026|NDUS7), *Homo sapiens* (O75251|NDUS7), *Mus musculus* (Q9DC70|NDUS7), *Arabidopsis thaliana* (Q42577|NDUS7), *Yarrowia lipolytica* (Q9UUT7|NUKM), *Thermus thermophilus* (Q56218|NQO6), *E. coli* (P0AFC7|NUOB), *Thermosynechococcus elongatus* (Q8DKZ4|NDHK), *Pyrococcus furiosus Mbs* (I6UZV3|MbhJ), *Pyrococcus furiosus* Mbh (P21853|PHNS), *Desulfovibrio gigas* (P12943|PHNS), *Cupriavidus_necator* (P31892|MBHS), *Hydrogenophilus thermoluteolus* (A0A2Z6DYQ5| HYDTE), *E. coli* (P69739|MBHS).

#### Supplementary tables

**Table S1. Key bond distances of DFT optimized [NiFe] active site from the soluble [NiFe] hydrogenase and comparison to a high-resolution x-ray structure.** The models were optimized in both singlet (S=0) and triplet (S=1) electronic configuration using different DFT functionals with variable amount of exact exchange. TPSSh-x2c denotes relativistic calculation performed with the X2C Hamiltonian along with the compatible basis-sets. The corresponding distances from the triplet spin-state geometry optimization are provided in the parenthesis.

| Bonds | X-Ray | BP86 | TPSS | TPSSh | TPSSh-x2c | B3LYP |
|---|---|---|---|---|---|---|
| **Ni-Fe** | 2.57 | 2.549 (2.573) | 2.540 (2.565) | 2.547 (2.561) | 2.539 (2.552) | 2.597 (2.588) |
| **Ni-$H^-$** | 1.58 | 1.593 (1.696) | 1.596 (1.694) | 1.588 (1.699) | 1.579 (1.688) | 1.576 (1.700) |
| **Fe-$H^-$** | 1.78 | 1.712 (1.636) | 1.705 (1.639) | 1.694 (1.635) | 1.693 (1.634) | 1.705 (1.639) |
| **Fe-CO** | 1.75 | 1.726 (1.749) | 1.735 (1.755) | 1.737 (1.752) | 1.729 (1.743) | 1.748 (1.764) |
| **Fe-CN1** | 1.88 | 1.859 (1.851) | 1.872 (1.865) | 1.876 (1.869) | 1.869 (1.863) | 1.891 (1.885) |
| **Fe-CN2** | 1.91 | 1.868 (1.863) | 1.880 (1.874) | 1.885 (1.877) | 1.879 (1.871) | 1.903 (1.893) |
| **C-O** | 1.15 | 1.172 (1.172) | 1.169 (1.169) | 1.159 (1.159) | 1.160 (1.161) | 1.152 (1.152) |
| **C-N1** | 1.17 | 1.179 (1.180) | 1.176 (1.177) | 1.169 (1.171) | 1.170 (1.171) | 1.163 (1.164) |
| **C-N2** | 1.15 | 1.179 (1.179) | 1.177 (1.176) | 1.170 (1.170) | 1.170 (1.170) | 1.163 (1.164) |
| **Ni-S($Cys^{81}$)** | 2.24 | 2.231 (2.333) | 2.229 (2.337) | 2.228 (2.338) | 2.218 (2.326) | 2.252 (2.356) |
| **Ni-S($Cys^{84}$)** | 2.21 | 2.182 (2.336) | 2.184 (2.342) | 2.185 (2.352) | 2.176 (2.346) | 2.209 (2.387) |
| **Ni-S($Cys^{546}$)** | 2.18 | 2.169 (2.276) | 2.179 (2.297) | 2.183 (2.322) | 2.172 (2.312) | 2.203 (2.355) |
| **Ni-S($Cys^{549}$)** | 2.54 | 2.578 (2.297) | 2.558 (2.303) | 2.600 (2.319) | 2.587 (2.308) | 2.699 (2.351) |
| **Fe-S($Cys^{84}$)** | 2.26 | 2.256 (2.349) | 2.258 (2.347) | 2.261 (2.347) | 2.255 (2.342) | 2.288 (2.382) |
| **Fe-S($Cys^{549}$)** | 2.31 | 2.338 (2.303) | 2.337 (2.311) | 2.334 (2.326) | 2.329 (2.322) | 2.359 (2.364) |

**Table S2. Mulliken spin population of key atoms in the Ni-R state of the [NiFe] soluble hydrogenase with various DFT functionals.** The calculations were performed for the S=1 (triplet) state. Geometry in each case is optimized using the same functional. TPSSh-x2c denotes relativistic calculation performed with the X2C Hamiltonian along with the compatible basis-sets. UKS (Unrestricted Kohn-Sham) spin-contamination is also reported for each functional (ideal $\langle S^2 \rangle$ for S=1.0 system is 2.00).

| | **BP86** | **TPSS** | **TPSSh** | **TPSSh-x2c** | **B3LYP** |
|---|---|---|---|---|---|
| **Fe** | 0.1503 | 0.1284 | 0.0927 | 0.1002 | 0.0859 |
| **Ni** | 1.0745 | 1.1796 | 1.3155 | 1.3465 | 1.3622 |
| **$S^{546}$** | 0.0836 | 0.0794 | 0.0770 | 0.0757 | 0.0768 |
| **$S^{549}$** | 0.2529 | 0.2250 | 0.1814 | 0.1704 | 0.1611 |
| **$S^{81}$** | 0.1436 | 0.1265 | 0.1067 | 0.1039 | 0.1012 |
| **$S^{84}$** | 0.1981 | 0.1783 | 0.1603 | 0.1565 | 0.1537 |
| **Hydride*** | 0.0222 | 0.0260 | 0.0306 | 0.0385 | 0.0290 |
| **$\langle S^2 \rangle$** | 2.0080 | 2.0094 | 2.0140 | 2.0138 | 2.0154 |

*Hydride atom between the Ni and Fe atom.

**Table S3. Comparison of adiabatic singlet-minus-triplet (S-T) gaps for the soluble [NiFe] hydrogenase obtained with various DFT functionals and wavefunction-based approaches.** Adiabatic singlet-triplet gaps were computed (using same geometry optimized in the singlet and triplet configuration) using variety of DFT functionals and compared with wavefunction based approaches (RPA and DLPNO-CCSD($T_1$)). The computations were performed using both relativistic and non-relativistic approaches. B3LYP** and B3LYP* are with 10% and 15% exact-exchange, respectively. The models used for benchmarking is shown in Figure S2.

| **Method** | **BP86** | **TPSS** | **TPSSh** | **B3LYP**** | **B3LYP*** | **B3LYP** | **RPA** | **DLPNO-CCSD($T_1$)** |
|---|---|---|---|---|---|---|---|---|
| **Relativistic** | -9.730 | -6.841 | -2.301 | -6.464 | -3.492 | -0.119 | -3.238 | -6.352 |
| **Non-Relativistic** | -8.969 | -6.115 | -1.741 | -5.834 | -2.942 | +0.352 | -2.531 | -5.566 |

Note: We have also tested a DLPNO-CCSD($T_1$) calculation with the "TightPNO" settings (using TPSSh reference orbitals) with the cc-pwCVTZ-DK basis-set on Fe and Ni, and cc-PVDZ on rest of the atoms, with a S-T gap of -6.581 kcal $mol^{-1}$.

**Table S4. Reaction energetics of the $H_2$ formation step in Mbh obtained with various DFT functional and wavefunction-based approaches.** The relative energy of the reactant, transition state and product computed using variety of DFT functionals and correlated RPA and DLPNO-CCSD(T) wavefunction approaches. B3LYP** and B3LYP* are with 10% and 15% exact-exchange, respectively. The QM cluster model used for this benchmarking is showcased in Figure S3.

| Method | BP86 | TPSS | TPSSh | B3LYP** | B3LYP* | B3LYP | RPA | DLPNO-CCSD(T) |
|---|---|---|---|---|---|---|---|---|
| **R** | 0 | 0 | 0 | 0 | 0 | 0 | 0.0 | 0 |
| **TS** | +10.109 | +10.076 | +9.937 | +9.973 | +9.807 | +9.639 | +8.990 | +8.637 |
| **P** | -1.973 | -0.563 | +3.183 | +1.255 | +3.346 | +5.475 | +3.599 | +4.820 |

**Table S5. Adiabatic spin state gap between low and high spin of the Ni-SI$_a$ models of Mbh.** The adiabatic low-spin minus high-spin gap (in kcal mol$^{-1}$) were computed using the B3LYP*/x2c-TZVPPall/x2c-TZVPall level of theory.

| Model number | $\Delta E_{LS-HS}$ (kcal mol$^{-1}$) |
|---|---|
| **$1A^{LS} – 1A^{HS}$** | -0.4 |
| **$1B^{LS} – 1B^{HS}$** | -0.5 |
| **$1D^{LS} – 1D^{HS}$** | -2.8 |
| **$1E^{LS} – 1E^{HS}$** | -0.4 |
| **$1F^{LS} – 1F^{HS}$** | +4.6 |
| **$1G^{LS} – 1G^{HS}$** | +4.3 |

**Table S6. Key bond distances of the DFT optimized Ni-SI$_a$ state of Mbh.** Key bond-distances of the Mbh [NiFe] active-site obtained from the DFT optimized quantum cluster models in the Ni-SI$_a$ state. The models were optimized in both singlet (S=0) and triplet (S=1) electronic configuration. The corresponding distances from the high-spin based geometry optimization are provided in brackets.

| Model | Ni-Fe | Fe-S$_{(C377)}$ | Fe-S$_{(C71)}$ | Ni-S$_{(C377)}$ | Ni-S$_{(C71)}$ | Ni-S$_{(C68)}$ | Ni-S$_{(C374)}$ |
|---|---|---|---|---|---|---|---|
| **1A$^{LS}$ [1A$^{HS}$]** | 2.83 [2.91] | 2.22 [2.29] | 2.31 [2.23] | 2.19 [2.37] | 2.28 [2.41] | 2.24 [2.26] | 2.15 [2.16] |
| **1B$^{LS}$ [1B$^{HS}$]** | 2.83 [2.91] | 2.22 [2.29] | 2.30 [2.23] | 2.19 [2.38] | 2.27 [2.39] | 2.23 [2.26] | 2.16 [2.18] |
| **1D$^{LS}$ [1D$^{HS}$]** | 2.84 [2.85] | 2.21 [2.29] | 2.30 [2.24] | 2.19 [2.33] | 2.24 [2.27] | 2.20 [2.23] | 2.21 [2.25] |
| **1E$^{LS}$ [1E$^{HS}$]** | 2.82 [2.90] | 2.24 [2.29] | 2.31 [2.23] | 2.19 [2.33] | 2.28 [2.44] | 2.25 [2.28] | 2.12 [2.13] |
| **1F$^{LS}$ [1F$^{HS}$]** | 2.68 [2.90] | 2.20 [2.30] | 2.28 [2.23] | 2.18 [2.39] | 2.25 [2.40] | 2.26 [2.26] | 2.15 [2.15] |
| **1G$^{LS}$ [1G$^{HS}$]** | 2.84 [2.95] | 2.21 [2.27] | 2.30 [2.25] | 2.19 [2.37] | 2.24 [2.32] | 2.19 [2.22] | 2.22 [2.29] |

**Table S7. Mulliken spin populations of key atoms in the Ni-SIa state of Mbh.** Mulliken spin-population of the key atoms in the core-structure of the [NiFe] cluster (high-spin, S=1) for all key models of the Ni-SI$_a$ state.

| Model | Fe | Ni | $S^{Cys377}$ | $S^{Cys374}$ | $S^{Cys68}$ | $S^{Cys71}$ |
|---|---|---|---|---|---|---|
| **$1A^{HS}$** | 0.07 | 1.33 | 0.12 | 0.22 | 0.18 | 0.04 |
| **$1B^{HS}$** | 0.06 | 1.35 | 0.13 | 0.18 | 0.19 | 0.05 |
| **$1D^{HS}$** | 0.05 | 1.32 | 0.19 | 0.07 | 0.24 | 0.07 |
| **$1E^{HS}$** | 0.02 | 1.31 | 0.11 | 0.33 | 0.12 | 0.08 |
| **$1F^{HS}$** | 0.08 | 1.32 | 0.10 | 0.23 | 0.18 | 0.03 |
| **$1G^{HS}$** | 0.03 | 1.34 | 0.16 | 0.09 | 0.25 | 0.08 |

**Table S8. Key bond distances of the DFT optimized Ni-L and Ni-C states of Mbh.** Key bond-distances of the Mbh [NiFe] active-site obtained from the DFT optimized quantum cluster models in the Ni-L/Ni-C state. The models were optimized with S=1/2 electronic configuration.

| Model | Ni-Fe | Fe-$S_{(C377)}$ | Fe-$S_{(C71)}$ | Ni-$S_{(C377)}$ | Ni-$S_{(C71)}$ | Ni-$S_{(C68)}$ | Ni-$S_{(C374)}$ | Fe-$H^-$ | Ni-$H^-$ |
|---|---|---|---|---|---|---|---|---|---|
| **2A** | 2.60 | 2.29 | 2.27 | 2.30 | 2.29 | 2.36 | 2.21 | na | na |
| **3A** | 2.59 | 2.28 | 2.27 | 2.29 | 2.29 | 2.36 | 2.21 | na | na |
| **3B** | 2.57 | 2.27 | 2.26 | 2.29 | 2.24 | 2.33 | 2.24 | na | na |
| **3D** | 2.55 | 2.29 | 2.26 | 2.35 | 2.26 | 2.23 | 2.17 | 1.70 | 1.60 |
| **3G** | 2.84 | 2.27 | 2.26 | 2.35 | 2.28 | 2.27 | 2.17 | na | 1.44 |
| **3G1** | 2.80 | 2.31 | 2.31 | 2.31 | 2.32 | 2.28 | 2.19 | na | na |
| **3J** | 2.62 | 2.29 | 2.27 | 2.30 | 2.29 | 2.35 | 2.21 | na | na |
| **3K** | 2.62 | 2.24 | 2.25 | 2.29 | 2.28 | 2.32 | 2.19 | na | na |
| **3X** | 2.85 | 2.30 | 2.22 | 2.88 | 2.23 | 2.26 | 2.20 | na | na |
| **3Y** | 2.84 | 2.26 | 2.30 | 2.35 | 2.37 | 2.37 | 2.25 | na | na |
| **3X-2** | 2.84 | 2.29 | 2.23 | 2.99 | 2.22 | 2.25 | 2.17 | na | na |
| **3Y-2** | 2.57 | 2.27 | 2.26 | 2.28 | 2.25 | 2.33 | 2.26 | na | na |
| **3E** | 2.55 | 2.29 | 2.26 | 2.35 | 2.26 | 2.23 | 2.17 | 1.70 | 1.60 |
| **3I** | 2.84 | 2.29 | 2.25 | 2.31 | 2.22 | 2.27 | 2.22 | na | 1.45 |
| **3H** | 2.84 | 2.27 | 2.25 | 2.36 | 2.27 | 2.28 | 2.18 | na | 1.44 |
| **3I1** | 2.83 | 2.36 | 2.31 | 2.41 | 2.32 | 2.22 | 2.11 | na | 1.51 |
| **3L** | 2.55 | 2.30 | 2.26 | 2.36 | 2.27 | 2.23 | 2.16 | 1.70 | 1.60 |

**Table S9. Mulliken spin populations of the key atoms in the Ni-L and Ni-C states of Mbh.** Mulliken spin-population of the key atoms in the core-structure of the [NiFe] cluster for all key models of the Ni-L state.

| Model | Fe | Ni | $S^{Cys377}$ | $S^{Cys374}$ | $S^{Cys68}$ | $S^{Cys71}$ | $H^-$ |
|---|---|---|---|---|---|---|---|
| **2A** | -0.50 | 1.06 | 0.14 | 0.11 | 0.07 | 0.04 | na |
| **3A** | -0.40 | 1.02 | 0.14 | 0.09 | 0.05 | 0.04 | na |
| **3B** | -0.21 | 0.91 | 0.15 | 0.03 | 0.04 | 0.03 | na |
| **3D** | ~0.0 | 0.64 | 0.25 | 0.12 | -0.02 | -0.01 | ~0.0 |
| **3G** | 0.06 | 0.58 | 0.04 | 0.08 | 0.20 | -0.02 | 0.02[t] |
| **3G1** | -0.32 | 0.94 | 0.13 | 0.12 | 0.03 | 0.04 | ~0.0 |
| **3J** | -0.52 | 1.08 | 0.14 | 0.11 | 0.07 | 0.04 | na |
| **3K** | -0.41 | 1.04 | 0.08 | 0.12 | 0.06 | 0.03 | na |
| **3X** | 0.29 | 0.67 | ~0.0 | 0.02 | 0.08 | -0.06 | na |
| **3Y** | -0.25 | 0.93 | 0.15 | 0.05 | 0.04 | 0.04 | na |
| **3X-2** | 0.35 | 0.63 | -0.01 | -0.01 | 0.08 | -0.06 | na |
| **3Y-2** | -0.24 | 0.92 | 0.15 | 0.04 | 0.04 | 0.04 | na |
| **3E** | ~0.0 | 0.65 | 0.26 | 0.10 | -0.02 | -0.02 | ~0.0 |
| **3I** | 0.07 | 0.58 | 0.03 | 0.03 | 0.28 | -0.01 | ~0.0[t] |
| **3H** | 0.06 | 0.60 | 0.03 | 0.06 | 0.22 | -0.02 | ~0.0 |
| **3I1** | ~0.0 | 0.58 | 0.24 | 0.18 | ~0.0 | ~0.0 | ~0.0 |
| **3L** | ~0.0 | 0.66 | 0.22 | 0.13 | -0.02 | -0.01 | 0.01 |

t: terminal hydride coordinated with Ni(III)

**Table S10. Adiabatic spin state gap between low and high spin form of the Ni-R state of Mbh.** The adiabatic low-spin minus high-spin gap (in kcal mol$^{-1}$) were computed using the B3LYP*/x2c-TZVPPall/x2c-TZVPall level of theory.

| Model | $\Delta E_{LS-HS}$ (kcal mol$^{-1}$) |
|---|---|
| **$4A^{LS} - 4A^{HS}$** | -1.7 |
| **$4B^{LS} - 4B^{HS}$** | -3.1 |
| **$4B1^{LS} - 4B1^{HS}$** | -24.8 |
| **$4X^{LS} - 4X^{HS}$** | -0.95 |
| **$5A^{LS} - 5A^{HS}$** | -3.45 |
| **$5B^{LS} - 5B^{HS}$** | -12.29 |
| **$5C^{LS} - 5C^{HS}$** | -26.44 |
| **$5E^{LS} - 5E^{HS}$** | +0.47 |
| **$5F^{LS} - 5F^{HS}$** | +1.53 |
| **$5C2^{LS} - 5C2^{HS}$** | -19.93 |
| **$5E2^{LS} - 5E2^{HS}$** | -3.49 |

**Table S11. Key bond distances of the DFT optimized Ni-R state of Mbh.** Key bond-distances of the Mbh [NiFe] active-site obtained from the DFT optimized quantum cluster models in the Ni-R state. The models were optimized in both singlet (S=0) and triplet (S=1) electronic configuration. The corresponding distances from the high-spin based geometry optimization are provided in brackets.

| Model | Ni-Fe | Fe-$S_{(C377)}$ | Fe-$S_{(C71)}$ | Ni-$S_{(C377)}$ | Ni-$S_{(C71)}$ | Ni-$S_{(C68)}$ | Ni-$S_{(C374)}$ | Fe-H- | Ni-$H^-$ |
|---|---|---|---|---|---|---|---|---|---|
| **$4A^{LS}$** **[$4A^{HS}$]** | 2.61 [2.62] | 2.33 [2.33] | 2.26 [2.34] | 2.74 [2.33] | 2.21 [2.42] | 2.22 [2.36] | 2.19 [2.26] | 1.70 [1.63] | 1.60 [1.71] |
| **$4B^{LS}$** **[$4B^{HS}$]** | 2.55 [2.56] | 2.34 [2.33] | 2.25 [2.37] | 2.58 [2.32] | 2.19 [2.37] | 2.22 [2.33] | 2.19 [2.33] | 1.70 [1.64] | 1.59 [1.69] |
| **$4B1^{LS}$** **[$4B1^{HS}$]** | 2.55 [2.56] | 2.29 [2.28] | 2.26 [2.32] | 2.36 [2.29] | 2.25 [2.46] | 2.23 [2.27] | 2.20 [2.22] | 1.73 [1.71] | 1.57 [1.61] |
| **$4D^{LS}$** **[$4D^{HS}$]** | 2.61 [2.92] | 2.24 [2.30] | 2.27 [2.23] | 2.29 [2.36] | 2.27 [2.42] | 2.27 [2.26] | 2.21 [2.17] | - | - |
| **$4B2^{LS}$** **[$4B2^{HS}$]** | 2.51 | 2.29 | 2.24 | 2.31 | 2.18 | 2.38 | 2.18 | na | 1.47 |
| **$4E^{LS}$** **[$4E^{HS}$]** | [2.78] | [2.33] | [2.32] | [2.33] | [2.40] | [2.27] | [2.19] | - | - |
| **$4X^{LS}$** **[$4X^{HS}$]** | 2.55 [2.56] | 2.34 [2.33] | 2.26 [2.34] | 2.68 [2.31] | 2.22 [2.39] | 2.16 [2.36] | 2.17 [2.21] | 1.73 [1.63] | 1.56 [1.68] |
| **$5A^{LS}$** **[$5A^{HS}$]** | 2.61 [2.63] | 2.34 [2.33] | 2.26 [2.34] | 2.74 [2.34] | 2.21 [2.43] | 2.23 [2.36] | 2.19 [2.24] | 1.70 [1.63] | 1.59 [1.71] |
| **$5B^{LS}$** **[$5B^{HS}$]** | 2.60 [2.66] | 2.30 [2.28] | 2.26 [2.33] | 2.68 [2.35] | 2.21 [2.42] | 2.23 [2.33] | 2.18 [2.23] | 1.69 [1.63] | 1.59 [1.72] |
| **$5C^{LS}$** **[$5C^{HS}$]** | 2.66 [2.62] | 2.35 [2.33] | 2.28 [2.31] | 3.44 [2.60] | 2.18 [2.39] | 2.23 [2.35] | 2.18 [2.25] | 1.68 [1.63] | 1.59 [1.70] |
| **$5E^{LS}$** **[$5E^{HS}$]** | 2.54 [2.56] | 2.34 [2.33] | 2.26 [2.34] | 2.54 [2.30] | 2.19 [2.37] | 2.23 [2.34] | 2.19 [2.33] | 1.71 [1.64] | 1.58 [1.69] |
| **$5F^{LS}$** **[$5F^{HS}$]** | 2.53 [2.55] | 2.33 [2.33] | 2.26 [2.34] | 2.50 [2.30] | 2.19 [2.36] | 2.23 [2.35] | 2.18 [2.31] | 1.71 [1.64] | 1.59 [1.69] |
| **$5C2^{LS}$** **[$5C2^{HS}$]** | 2.63 [2.64] | 2.29 [2.29] | 2.26 [2.31] | 2.86 [2.71] | 2.21 [2.40] | 2.22 [2.34] | 2.17 [2.19] | 1.70 [1.63] | 1.60 [1.70] |
| **$5E2^{LS}$** **[$5E2^{HS}$]** | 2.55 [2.56] | 2.33 [2.33] | 2.25 [2.34] | 2.57 [2.32] | 2.20 [2.36] | 2.23 [2.33] | 2.19 [2.34] | 1.70 [1.64] | 1.58 [1.70] |

**$4B1^{LS}$**: terminal Ni-$H^-$ is 1.44 Å, **$4B1^{HS}$**: terminal Ni-$H^-$ is 1.46 Å, **$4D^{LS}$**: Ni-$H_2$ is 1.59 Å, **$4E^{HS}$**: Fe-$H_2$ is 1.68 Å and Ni-$H_2$ is 2.08 Å.

**Table S12. Mulliken spin populations of the key atoms in the Ni-R state of Mbh.** Mulliken spin-population of the key atoms in the core-structure of the [NiFe] cluster for all key models of the Ni-R state (high-spin models, S=1).

| Model | Fe | Ni | $S^{Cys377}$ | $S^{Cys374}$ | $S^{Cys68}$ | $S^{Cys71}$ | $H^-$ |
|---|---|---|---|---|---|---|---|
| **$4A^{HS}$** | 0.14 | 1.31 | 0.15 | 0.14 | 0.11 | 0.12 | 0.03 |
| **$4B^{HS}$** | 0.11 | 1.32 | 0.18 | 0.08 | 0.10 | 0.16 | 0.04 |
| **$4B1^{HS}$** | 0.07 | 0.61 | 0.11 | 0.84 | 0.12 | 0.21 | 0.02 |
| **$4D^{HS}$** | 0.07 | 1.33 | 0.13 | 0.21 | 0.18 | 0.04 | na |
| **$4E^{HS}$** | 0.02 | 1.30 | 0.14 | 0.21 | 0.20 | 0.10 | na |
| **$4X^{HS}$** | 0.17 | 1.25 | 0.16 | 0.21 | 0.08 | 0.11 | 0.03 |
| **$5A^{HS}$** | 0.15 | 1.30 | 0.13 | 0.18 | 0.11 | 0.11 | 0.01 |
| **$5B^{HS}$** | 0.14 | 1.32 | 0.07 | 0.20 | 0.13 | 0.12 | 0.01 |
| **$5C^{HS}$** | 0.15 | 1.35 | 0.04 | 0.19 | 0.13 | 0.13 | 0.01 |
| **$5E^{HS}$** | 0.12 | 1.28 | 0.23 | 0.08 | 0.10 | 0.15 | 0.04 |
| **$5F^{HS}$** | 0.12 | 1.28 | 0.24 | 0.08 | 0.09 | 0.15 | 0.05 |
| **$5C2^{HS}$** | 0.16 | 1.28 | 0.02 | 0.29 | 0.13 | 0.11 | ~0.00 |
| **$5E2^{HS}$** | 0.12 | 1.32 | 0.17 | 0.08 | 0.11 | 0.16 | 0.04 |

**Table S13. $H_2$ binding free energy in the Ni-R state of Mbh.** The binding free energy (in kcal $mol^{-1}$) of the $H_2$ to the [NiFe] active site in Mbh and DvMF (soluble hydrogenases) is showcased. The binding energy is computed as $\Delta G_{complex}$ – ($\Delta G_{H2} + \Delta G_{apo\ [NiFe]}$). For Mbh, both LS and HS models with bound $H_2$ were investigated, whereas only the LS was studied for DvMF (see DFT model in Figure S1). The geometry optimizations for the complex ($H_2$ and apo forms) were performed using TPSSh level with the central core of the active site comprising of Fe, Ni, S, two CN- and CO assigned def2-TZVP basis-set, while where the rest of the atoms were described with the def2-SVP basis sets. The single-point energy computations were performed using the B3LYP* functional with x2c-TZVPP for Fe and Ni, and x2c-TZVP basis-set for rest of the atoms. The molecular Hessian used for estimation of free energies were computed numerically at the TPSSh/def2-TZVP/def2-SVP level of theory

| **Model** | $\Delta E_{Binding}$ | $\Delta(E+ZPE)_{Binding}$ | $\Delta G_{Binding}$ |
|---|---|---|---|
| | **Mbh** | | |
| **$4D^{LS}$** (Ni-$H_2$) | -4.7 | -2.1 | +7.2 |
| **$4E^{HS}$** (Fe-$H_2$) | -3.9 | -2.4 | +7.1 |
| | ***DvMF*** | | |
| **LS (Ni-$H_2$)*** | -11.4 | -7.8 | +2.1 |

*Ni-$H_2$ is 1.56 Å

**Table S14. Adiabatic spin state gap between low and high spin form of the Ni-SI$_a$ models in the presence of His75$_L^+$.** The adiabatic low-spin minus high-spin gap (in kcal mol$^{-1}$) were computed using the B3LYP*/x2c-TZVPPall/x2c-TZVPall level of theory.

| Model number | $\Delta E_{LS-HS}$ (kcal mol$^{-1}$) |
|---|---|
| **1A$^{LS}$His75$^+$ − 1A$^{HS}$His75$^+$** | +4.6 |
| **1B$^{LS}$His75$^+$ − 1B$^{HS}$His75$^+$** | +9.4 |
| **1D$^{LS}$His75$^+$ − 1D$^{HS}$His75$^+$** | +0.2 |
| **1E$^{LS}$His75$^+$ − 1E$^{HS}$His75$^+$** | +0.6 |
| **1G$^{LS}$His75$^+$ − 1G$^{HS}$His75$^+$** | +5.5 |

**Table S15. Key bond distances of the DFT optimized Ni-SI$_a$ state in the presence of His75$_L^+$.** Key bond-distances of the Mbh [NiFe] active-site obtained from the DFT optimized quantum cluster models in the Ni-SI$_a$ state. The models were optimized in both singlet (S=0) and triplet (S=1) electronic configuration. The corresponding distances from the high-spin based geometry optimization are provided in brackets.

| Model | Ni-Fe | Fe-S$_{(C377)}$ | Fe-S$_{(C71)}$ | Ni-S$_{(C377)}$ | Ni-S$_{(C71)}$ | Ni-S$_{(C68)}$ | Ni-S$_{(C374)}$ |
|---|---|---|---|---|---|---|---|
| **1A$^{LS}$His75$^+$ [1A$^{HS}$His75$^+$]** | 2.68 [2.90] | 2.20 [2.30] | 2.28 [2.23] | 2.18 [2.39] | 2.25 [2.40] | 2.26 [2.26] | 2.15 [2.15] |
| **1B$^{LS}$His75$^+$ [1B$^{HS}$His75$^+$]** | 2.68 [2.91] | 2.20 [2.30] | 2.28 [2.23] | 2.18 [2.39] | 2.25 [2.39] | 2.26 [2.26] | 2.15 [2.16] |
| **1D$^{LS}$His75$^+$ [1D$^{HS}$His75$^+$]** | 2.84 [2.85] | 2.22 [2.30] | 2.30 [2.24] | 2.19 [2.35] | 2.24 [2.28] | 2.20 [2.23] | 2.22 [2.26] |
| **1E$^{LS}$His75$^+$ [1E$^{HS}$His75$^+$]** | 2.82 [2.89] | 2.24 [2.30] | 2.31 [2.23] | 2.19 [2.35] | 2.28 [2.43] | 2.26 [2.28] | 2.12 [2.13] |
| **1G$^{LS}$His75$^+$ [1G$^{HS}$His75$^+$]** | 2.84 [2.94] | 2.22 [2.28] | 2.31 [2.24] | 2.20 [2.39] | 2.23 [2.32] | 2.19 [2.22] | 2.23 [2.29] |

**Table S16. Mulliken spin population of key atoms in the Ni-SI$_a$ state in the presence of His75$_L^+$.** Mulliken spin-population of the key atoms in the core-structure of the [NiFe] cluster (high-spin, S=1) for all key models of the Ni-SI$_a$ state.

| Model | Fe | Ni | $S^{Cys377}$ | $S^{Cys374}$ | $S^{Cys68}$ | $S^{Cys71}$ |
|---|---|---|---|---|---|---|
| **1A$^{HS}$His75$^+$** | 0.08 | 1.32 | 0.10 | 0.23 | 0.18 | 0.03 |
| **1B$^{HS}$His75$^+$** | 0.07 | 1.34 | 0.11 | 0.20 | 0.19 | 0.03 |
| **1D$^{HS}$His75$^+$** | 0.06 | 1.33 | 0.17 | 0.07 | 0.24 | 0.06 |
| **1E$^{HS}$His75$^+$** | 0.02 | 1.30 | 0.10 | 0.35 | 0.12 | 0.07 |
| **1G$^{HS}$His75$^+$** | 0.04 | 1.34 | 0.15 | 0.09 | 0.25 | 0.08 |

**Table S17. Key bond distances of the DFT optimized Ni-L and Ni-C states in the presence of His75$_L^+$.** Key bond-distances of the Mbh [NiFe] active-site obtained from the DFT optimized quantum cluster models in the Ni-L/Ni-C state. The models were optimized with S=1/2 electronic configuration with His75$^+$.

| Model | Ni-Fe | Fe-$S_{(C377)}$ | Fe-$S_{(C71)}$ | Ni-$S_{(C377)}$ | Ni-$S_{(C71)}$ | Ni-$S_{(C68)}$ | Ni-$S_{(C374)}$ | Fe-H$^-$ | Ni-H$^-$ |
|---|---|---|---|---|---|---|---|---|---|
| **2A-His75$^+$** | 2.62 | 2.29 | 2.27 | 2.30 | 2.29 | 2.35 | 2.21 | na | na |
| **3A-His75$^+$** | 2.60 | 2.28 | 2.26 | 2.30 | 2.29 | 2.36 | 2.21 | na | na |
| **3B-His75$^+$** | 2.57 | 2.27 | 2.25 | 2.29 | 2.24 | 2.33 | 2.25 | na | na |
| **3Y2-His75$^+$** | 2.57 | 2.27 | 2.26 | 2.28 | 2.25 | 2.32 | 2.27 | na | na |
| **3D-His75$^+$** | 2.55 | 2.30 | 2.26 | 2.36 | 2.27 | 2.23 | 2.16 | 1.70 | 1.60 |
| **3G-His75$^+$** | 2.84 | 2.28 | 2.26 | 2.37 | 2.28 | 2.27 | 2.17 | na | 1.44 |
| **3G1-His75$^+$** | 2.79 | 2.33 | 2.31 | 2.32 | 2.32 | 2.29 | 2.18 | na | na |
| **3E-His75$^+$** | 2.55 | 2.29 | 2.26 | 2.35 | 2.26 | 2.23 | 2.17 | 1.70 | 1.60 |
| **3I-His75$^+$** | 2.83 | 2.30 | 2.25 | 2.32 | 2.22 | 2.26 | 2.22 | na | 1.45 |
| **3H-His75$^+$** | 2.84 | 2.28 | 2.26 | 2.37 | 2.27 | 2.27 | 2.18 | na | 1.44 |
| **3I1-His75$^+$** | 2.83 | 2.37 | 2.31 | 2.42 | 2.32 | 2.22 | 2.11 | na | 1.51 |

**Table S18. Mulliken spin populations of the key atoms in the Ni-L and Ni-C state in the presence of His75$_L^+$.** Mulliken spin-population of the key atoms in the core-structure of the [NiFe] cluster for all key models of the Ni-L and Ni-C state.

| Model | Fe | Ni | $S^{Cys377}$ | $S^{Cys374}$ | $S^{Cys68}$ | $S^{Cys71}$ | $H^-$ |
|---|---|---|---|---|---|---|---|
| **2A-His75$^+$** | -0.52 | 1.08 | 0.14 | 0.11 | 0.07 | 0.04 | na |
| **3A-His75$^+$** | -0.43 | 1.04 | 0.13 | 0.09 | 0.06 | 0.04 | na |
| **3B-His75$^+$** | -0.23 | 0.93 | 0.14 | 0.03 | 0.04 | 0.03 | na |
| **3D-His75$^+$** | ~0.0 | 0.66 | 0.22 | 0.13 | -0.02 | -0.01 | 0.01 |
| **3G-His75$^+$** | 0.08 | 0.59 | 0.03 | 0.07 | 0.20 | -0.02 | ~0.0$^t$ |
| **3G1-His75$^+$** | -0.36 | 0.97 | 0.13 | 0.13 | 0.04 | 0.05 | ~0.0 |
| **3Y2-His75$^+$** | -0.26 | 0.94 | 0.14 | 0.04 | 0.04 | 0.04 | na |
| **3E-His75$^+$** | ~0.0 | 0.67 | 0.23 | 0.11 | -0.02 | -0.01 | ~0.0 |
| **3I-His75$^+$** | 0.08 | 0.58 | 0.02 | 0.03 | 0.29 | -0.02 | -0.01$^t$ |
| **3H-His75$^+$** | 0.08 | 0.61) | 0.02 | 0.05 | 0.22 | -0.02 | ~0.0 |
| **3I1-His75$^+$** | -0.01 | 0.59 | 0.21 | 0.20 | ~0.0 | ~0.0 | ~0.0 |

t: terminal hydride coordinated with Ni(III)

**Table S19. Adiabatic spin state gap between low and high spin forms of the Ni-R state in the presence of $His75_L^+$.** Adiabatic spin-state gap (low-spin *minus* high-spin, in kcal $mol^{-1}$) between Ni-R models using B3LYP*/x2c-TZVPPall/x2c-TZVPall level of theory.

| Model | $\Delta E_{LS-HS}$ (kcal $mol^{-1}$) |
|---|---|
| **$4A^{LS}His75^+ - 4A^{HS}His75^+$** | -2.9 |
| **$4B^{LS}His75^+ - 4B^{HS}His75^+$** | -4.4 |
| **$4B1^{LS}His75^+ - 4B1^{HS}His75^+$** | -24.3 |
| **$4X^{LS}His75^+ - 4X^{HS}His75^+$** | -6.5 |
| **$5E2^{LS}His75^+ - 5E2^{HS}His75^+$** | -4.47 |

**Table S20. Key bond distances of the DFT optimized Ni-R state in the presence of $His75_L^+$.** Key bond-distances of the Mbh [NiFe] active-site obtained from the DFT optimized quantum cluster models in the Ni-R state with protonated $His75_L^+$. The models were optimized in both singlet (S=0) and triplet (S=1) electronic configuration. The corresponding distances from the high-spin based geometry optimization are provided in brackets.

| Model | Ni-Fe | Fe-$S_{(C377)}$ | Fe-$S_{(C71)}$ | Ni-$S_{(C377)}$ | Ni-$S_{(C71)}$ | Ni-$S_{(C68)}$ | Ni-$S_{(C374)}$ | Fe-$H^-$ | Ni-$H^-$ |
|---|---|---|---|---|---|---|---|---|---|
| **$4A^{LS}His75^+$ [$4A^{HS}His75^+$]** | 2.61 [2.63] | 2.33 [2.33] | 2.26 [2.33] | 2.73 [2.33] | 2.21 [2.43] | 2.22 [2.36] | 2.19 [2.25] | 1.70 [1.63] | 1.59 [1.71] |
| **$4B^{LS}His75^+$ [$4B^{HS}His75^+$]** | 2.55 [2.57] | 2.34 [2.33] | 2.25 [2.34] | 2.59 [2.32] | 2.19 [2.37] | 2.22 [2.33] | 2.19 [2.33] | 1.70 [1.64] | 1.59 [1.69] |
| **$4B1^{LS}His75^+$ [$4B1^{HS}His75^+$]** | 2.55 [2.55] | 2.30 [2.29] | 2.26 [2.32] | 2.38 [2.29] | 2.26 [2.47] | 2.23 [2.27] | 2.20 [2.23] | 1.73 [1.71] | 1.57 [1.61] |
| **$4D^{LS}His75^+$ [$4D^{HS}His75^+$]** | 2.60 [2.91] | 2.25 [2.30] | 2.27 [2.23] | 2.31 [2.38] | 2.28 [2.41] | 2.26 [2.26] | 2.21 [2.16] | - | - |
| **$4E^{LS}His75^+$ [$4E^{HS}His75^+$]** | [2.79] | [2.34] | [2.32] | [2.34] | [2.41] | [2.27] | [2.19] | - | - |
| **$4X^{LS}His75^+$ [$4X^{HS}His75^+$]** | 2.85 [2.84] | 2.40 [2.43] | 2.31 [2.49] | 2.90 [2.37] | 2.25 [2.46] | 2.16 [2.34] | 2.15 [2.19] | 2.08 [1.96] | 1.53 [1.66] |
| **$5E2^{LS}His75^+$ [$5E2^{HS}His75^+$]** | 2.56 [2.57] | 2.33 [2.33] | 2.26 [2.34] | 2.58 [2.32] | 2.20 [2.37] | 2.22 [2.32] | 2.20 [2.35] | 1.70 [1.64] | 1.59 [1.70] |

**$4B1^{LS}His75^+$**: Ni-$H^-$ is 1.44 Å, **$4B1^{HS}His75^+$**: Ni-$H^-$ is 1.46 Å, **$4D^{LS}His75^+$**: Ni-$H_2$ is 1.56 Å, **$4E^{HS}His75^+$**: Fe-$H_2$ is 1.69 Å.

**Table S21. Mulliken spin populations of key atoms in the Ni-R state in the presence of $His75_L^+$.** Mulliken spin-population of the key atoms in the core-structure of the [NiFe] cluster for all key models of the Ni-R state (High-spin models, S=1).

| Model | Fe | Ni | $S^{Cys377}$ | $S^{Cys374}$ | $S^{Cys68}$ | $S^{Cys71}$ | $H^-$ |
|---|---|---|---|---|---|---|---|
| **$4A^{HS}His75^+$** | 0.14 | 1.31 | 0.14 | 0.15 | 0.12 | 0.12 | 0.02 |
| **$4B^{HS}His75^+$** | 0.11 | 1.33 | 0.16 | 0.09 | 0.11 | 0.16 | 0.04 |
| **$4B1^{HS}His75^+$** | 0.06 | 0.62 | 0.08 | 0.88 | 0.13 | 0.21 | 0.02 |
| **$4D^{HS}His75^+$** | 0.09 | 1.32 | 0.12 | 0.22 | 0.18 | 0.03 | na |
| **$4E^{HS}His75^+$** | 0.02 | 1.30 | 0.13 | 0.23 | 0.20 | 0.10 | na |
| **$4X^{HS}His75^+$** | 0.26 | 1.21 | 0.12 | 0.23 | 0.08 | 0.10 | 0.02 |
| **$5E2^{HS}His75^+$** | 0.12 | 1.33 | 0.16 | 0.08 | 0.12 | 0.16 | 0.03 |

**Table S22. Redox state of [NiFe] active-site in MD simulations.** Summary of the oxidation states of Ni and Fe in various [NiFe] catalytic states in the classical MD parametrization. Information on the protonation state of the $Cys374_L$ and bridging position (X) between Ni and Fe is also provided, including net spin and charge of the [NiFe] active site. $Ni^{II}$ is modelled in low-spin in both $Ni\text{-}SI_a$ and Ni-R states.

| Simulation | [NiFe] State | $Cys374_L$ | Ni | X | Fe | Spin | Charge |
|---|---|---|---|---|---|---|---|
| 1A | $Ni\text{-}SI_a$ | - | +2 | - | +2 | S=0 | -2 |
| 2A | $Ni\text{-}SI_a$ | - | +2 | - | +2 | S=0 | -2 |
| 3A | Ni-L | $H^+$ | +1 | - | +2 | S=1/2 | -2 |
| 4A | Ni-C | - | +3 | $H^-$ | +2 | S=1/2 | -2 |
| 5A | Ni-C | - | +3 | $H^-$ | +2 | S=1/2 | -2 |
| 6A | Ni-R | $H^+$ | +2 | $H^-$ | +2 | S=0 | -2 |

**Table S23. Complete list of classical MD simulations.** Summary of the classical MD simulations with modeled redox states of the FeS clusters (Fd1 and Fd2 from the Ferredoxin, and N6A, B6B and N2 from the Mbh peripheral arm; "ox" / "red" refer to the $2Fe^{3+}2Fe^{2+}/1Fe^{3+}3Fe^{2+}$ configuration), the protonation states of $His75_L$, $Glu20_L$, $Glu20_L$ and $Cys374_L$. Details on the oxidation state of the Ni and Fe and protonation state of the bridging position (X) is also provided (see Table S22). Simulation 2A_R and 5A_R are replicas of 2A and 5A, respectively.

| Sim | Fd1 | Fd2 | N6A | N6B | N2 | $H75_L$ | $E20_L$ | $E21_L$ | $C374_L$ | Ni | X | Fe | Spin | Time |
|---|---|---|---|---|---|---|---|---|---|---|---|---|---|---|
| 1A | red | red | ox | ox | ox | $H^\varepsilon$ | $H^+$ | $H^+$ | - | +2 | - | +2 | S=0 | 0.5 μs |
| 2A | ox | ox | red | ox | red | $H^\varepsilon$ | $H^+$ | $H^+$ | - | +2 | - | +2 | S=0 | 0.5 μs |
| 3A | ox | ox | ox | ox | red | $H^\varepsilon$ | $H^+$ | - | $H^+$ | +1 | - | +2 | S=1/2 | 0.5 μs |
| 4A | ox | ox | ox | ox | red | $H^\varepsilon$ | $H^+$ | - | - | +3 | $H^-$ | +2 | S=1/2 | 0.5 μs |
| 5A | ox | ox | ox | ox | red | $H^\varepsilon$ | - | $H^+$ | - | +3 | $H^-$ | +2 | S=1/2 | 0.5 μs |
| 6A | ox | ox | ox | ox | ox | $H^\varepsilon$ | - | - | $H^+$ | +2 | $H^-$ | +2 | S=0 | 0.5 μs |
| 2A_R | ox | ox | red | ox | red | $H^\varepsilon$ | $H^+$ | $H^+$ | - | +2 | - | +2 | S=0 | 0.25 μs |
| 5A_R | ox | ox | ox | ox | red | $H^\varepsilon$ | - | $H^+$ | - | +3 | $H^-$ | +2 | S=1/2 | 0.25 μs |
| | | | | | | | | | | | | | | 3.5 μs |

**Table S24. List of non-standard protonation states in the MD simulations.** Summary of non-standard protonation of residues used in the classical molecular dynamics simulations (Table S23). The protonation states were computed using Poisson-Boltzmann electrostatic calculations with Monte Carlo (28, 29) sampling approach as previously calculated [(30)]. Unless otherwise state, all histidine residues were modelled in their δ-neutral tautomer. Nomenclature ε denotes histidine protonated at only $N_\varepsilon$ atom (neutral). ε-δ denotes histidine protonated at both $N_\varepsilon$ and $N_\delta$ atoms (+1 charge).

| Subunits | Residues |
|---|---|
| MbhE | His49(ε-δ), Asp62 |
| MbhH | Lys354 |
| MbhK | His56(ε-δ), Glu114 |
| MbhL | His12(ε-δ), His192(ε-δ), His281(ε-δ), His75(ε), His325(ε), Glu187, Glu65, Glu20*, Glu21*, Cys374* |
| MbhM | His172(ε-δ), Glu141, Glu213 |
| MbhN | Glu27, Glu66, Glu97, Asp96 |

*The protonation states of these residues were systematically altered based on the redox states of [NiFe] cluster (see Table S23)

#### Supplementary movies

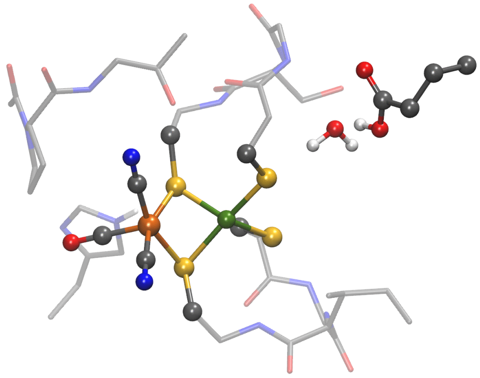

**Movie S1. Catalytic progression during the Ni-SI$_a$ → Ni-L transition.**

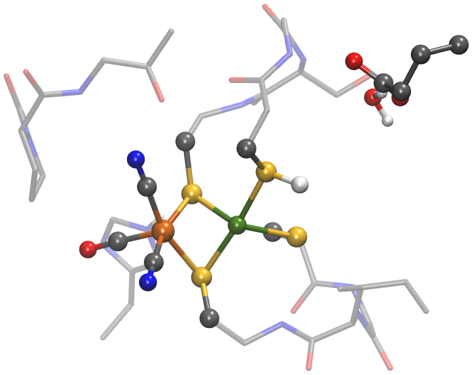

**Movie S2. Catalytic progression during the Ni-L → Ni-C transition.**

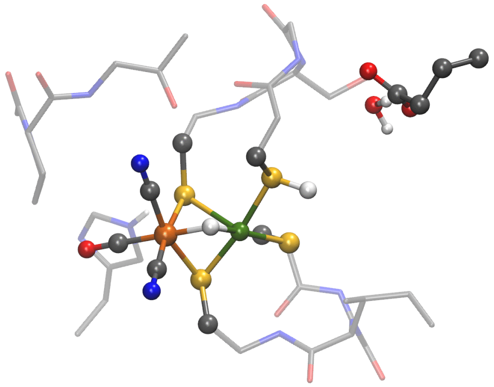

**Movie S3. $H_2$ formation in the Ni-R state.**
